# All-optical visualization of specific molecules in the ultrastructural context of brain tissue

**DOI:** 10.1101/2022.04.04.486901

**Authors:** Ons M’Saad, Ravikiran Kasula, Ilona Kondratiuk, Phylicia Kidd, Hanieh Falahati, Juliana E. Gentile, Robert F. Niescier, Katherine Watters, Robert C. Sterner, Seong Lee, Xinran Liu, Pietro De Camilli, James E. Rothman, Anthony J. Koleske, Thomas Biederer, Joerg Bewersdorf

## Abstract

Understanding the molecular anatomy and neural connectivity of the brain requires imaging technologies that can map the 3D nanoscale distribution of specific proteins in the context of brain ultrastructure. Light and electron microscopy (EM) enable visualization of either specific labels or anatomical ultrastructure, but combining molecular specificity with anatomical context is challenging. Here, we present pan-Expansion Microscopy of tissue (pan-ExM-t), an all-optical mouse brain imaging method that combines ∼24-fold linear expansion of biological samples with fluorescent pan-staining of protein densities (providing EM-like ultrastructural context), and immunolabeling of protein targets (for molecular imaging). We demonstrate the versatility of this approach by imaging the established synaptic markers Homer1, Bassoon, PSD-95, Synaptophysin, the astrocytic protein GFAP, myelin basic protein (MBP), and anti-GFP antibodies in dissociated neuron cultures and mouse brain tissue sections. pan-ExM-t reveals these markers in the context of ultrastructural features such as pre and postsynaptic densities, 3D nanoarchitecture of neuropil, and the fine structures of cellular organelles. pan-ExM-t is adoptable in any neurobiological laboratory with access to a confocal microscope and has therefore broad applicability in the research community.

**Highlights:**

- pan-ExM-t visualizes proteins in the context of synaptic ultrastructure
- Lipid labeling in pan-ExM-t reveals organellar and cellular membranes
- All-optical, easily accessible alternative to correlative light/electron microscopy
- High potential for high throughput connectomics studies

## Introduction

Three-dimensional microscopy techniques are instrumental to our understanding of brain organization with its complex morphology, spanning from the sub-synapse scale to neural circuitry maps. Despite significant advances in imaging, no microscopy method can provide a detailed molecular topography of the synapse (Kavalali and Jorgenson, 2014). While electron microscopy (EM) is the gold standard for ultrastructural analysis, localizing specific proteins still relies on immunogold labeling or electron-dense peroxidase substrates, neither of which are reliable nor quantitative (Gonda, 1998). Fluorescence microscopy, on the other hand, enables highly specific, multicolor labeling of proteins of interest. However, it fails to reveal the underlying ultrastructural context. Even with the advent of super-resolution microscopy, where single molecules can be imaged in 3D with spatial resolutions down to ∼10 nm (Zhang et al., 2020), delineating context is unattainable: fluorescent dyes are comparatively bulky (∼1 nm) and susceptible to quenching when densely packed, hindering imaging of crowded biomolecules (Conroy et al., 2016; Baddeley and Bewersdorf, 2018).

Expansion Microscopy (ExM) (Chen et al., 2015; Gallagher and Zhao, 2021), which physically expands tissue samples ∼4 to 20-fold by embedding them in swellable hydrogels, has resolved subcellular protein distributions of synaptic targets in mouse brain tissue (Chen et al., 2015; Tillberg et al., 2016; Ku et al., 2016; Chozinski et al., 2016; Chang et al., 2017; Murakami et al., 2018; Truckenbrodt et al., 2019; Park et al., 2019; Shen et al., 2020; Campbell et al., 2021; Gao et al., 2021; Park et al., 2021; Damstra et al., 2022), as well as map RNA transcripts within axons and dendrites in the brain (Alon et al., 2021). However, while the expansion-related decrowding avoids fluorescence quenching, ExM has traditionally capitalized almost exclusively on antibody labelings, and no ExM method has demonstrated correlative molecular and contextual imaging with synaptic resolution so far. Correlative light and electron microscopy (CLEM) (Hoffman et al., 2020) has been the only demonstrated option aligning specific molecular markers with sample ultrastructure. However, because of its operational complexity and challenge to label endogenous proteins specifically, especially in 3D imaging, its application has been very limited. We recently discovered a new principle for an optical contrast equivalent to EM heavy-metal stains, which allows for ultrastructural analysis using conventional light microscopy (M’Saad and Bewersdorf, 2020). By physically expanding a biological sample ∼20-fold in every dimension without losing the protein content, bulk staining of proteins can now reveal the totality of subcellular compartments down to size scales of ∼15 nm. We call this approach pan-ExM, referencing the philosophy of labeling the whole (Greek: pan) and the original concept of sample expansion in ExM (Chen et al., 2015). The 20^3^=8,000-fold volume expansion decrowds the cellular environment and thereby overcomes the permeability, quenching, and sampling limitations associated with bulk staining of conventional samples. In our previous work, we showed how this approach allows for conventional light microscopes to acquire EM-like images of adherent monolayer cells, revealing subcellular features such as mitochondria cristae and Golgi cisternae by their anatomical characteristics (M’Saad and Bewersdorf, 2020).

The expansion mechanism in pan-ExM differs from other ExM approaches in that the sample expands ∼20-fold while proteins are retained in the hydrogel for subsequent staining. To achieve this, pan-ExM avoids protease digestion and instead uses polymer entanglement (Edwards, 1967) to expand sample components twice without losing them: by embedding an already expanded sample prepared with a cleavable crosslinker in a second dense superabsorbent hydrogel, entanglements between polymer chains of the first and final hydrogels can physically interlock protein-polymer hybrids in this latter polymer network, thereby preserving the proteome while iteratively expanding it. The conceptual advance in pan-staining of highly expanded samples takes light microscopy to the realm of ultrastructural context imaging and generates EM heavy-metal stain-like images. In combination with well-established labeling methods in fluorescence microscopy, pan-staining provides nanoscale context to proteins of interest, analog to CLEM.

Here, we introduce pan-ExM-t, a pan-ExM protocol that enables contextual imaging of thick mouse brain tissue sections. Analogous to EM, we discovered that hallmark ultrastructural features such as pre- and postsynaptic densities can be identified by their morphological characteristics, allowing, for the first-time, ultrastructural imaging of putative synapses by light microscopy without specific labels. The developments we present in this paper give every neurobiologist the power to perform routine 3D pan-ExM-t imaging of brain tissue sections using their standard confocal microscope.

**Figure 1** shows an overview of our novel brain-tissue pan-ExM-t protocol. In brief, mice are transcardially perfused with fixative containing both formaldehyde (FA) and acrylamide (AAm) and their brains are extracted surgically and incubated in the same fixative overnight at 4°C. The brains are then sectioned at 50-100 µm thickness using a vibratome and stored in PBS until future use (**Fig. 1a**). Each tissue section to be expanded is embedded in a dense poly(acrylamide/sodium acrylate) co-polymer that is cross-linked with N,N′-(1,2-dihydroxyethylene)bisacrylamide (DHEBA), an acrylamide crosslinker with a cleavable amidomethylol bond (**Fig. 1b**). After polymerization, the now tissue-hydrogel hybrid is denatured with sodium dodecyl sulfate (SDS) in heated buffer (pH 6.8) for 4 hours (**Fig. 1c**) and expanded ∼5-fold in deionized water (**Fig. 1d**, blue). Next, a specific region of interest (ROI, ∼8x8 mm^2^) is cut and re-embedded first in a neutral polyacrylamide hydrogel cross-linked with DHEBA (**Fig. 1d**, green) and then in a poly(acrylamide/sodium acrylate) co-polymer cross-linked with N,N′-methylenebis(acrylamide) (BIS), a non-hydrolysable acrylamide crosslinker (**Fig. 1d**, orange). As we previously demonstrated (M’Saad and Bewersdorf, 2020), no secondary fixation of proteins before the re-embedding is required. The sample is then incubated in 200 mM sodium hydroxide to cleave DHEBA so that the crosslinked first and second hydrogel polymer networks are disentangled. After neutralization with multiple PBS washing steps, the sample is labeled with antibodies, pan-stained with fluorescent dyes to show protein-dense areas, washed with detergents, and expanded to its final size of ∼24-fold in ultrapure water (**Fig. 1e**). The expanded (and as a side-effect: optically cleared) sample is finally imaged on a standard confocal microscope (**Fig. 1f**) and can be stored at 4°C for months.

**Figure 1:**
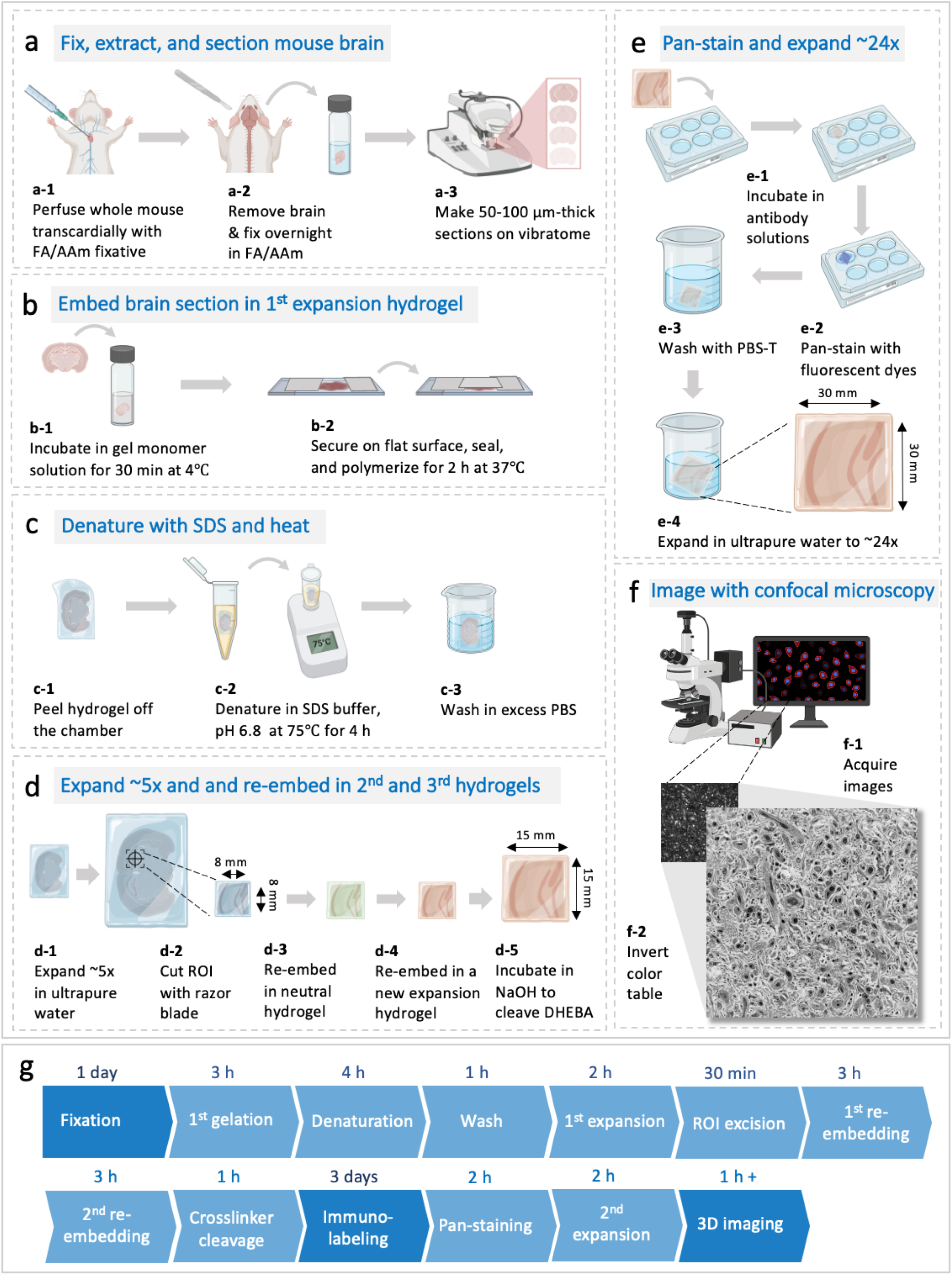
pan-ExM-t workflow for mouse brain tissue sections. (**a-f**) Experimental workflow. (**g**): Timeline summarizing the protocol. Abbreviations: FA: formaldehyde; AAm: acrylamide; NaOH: sodium hydroxide; DHEBA: N,N′-(1,2-dihydroxyethylene)bis-acrylamide; SDS: sodium dodecyl sulfate; PBS-T: 0.1% (v/v) TX-100 in PBS; ROI: region of interest.

## Results

### pan-ExM reveals synapse ultrastructure in dissociated neurons

Before experimenting with expanding mouse brain tissue sections, we tested pan-ExM in dissociated hippocampal rat and mice neurons using our published protocol with modifications in sample fixation (see **Methods**). We sought to test whether pan-ExM in neurons would reveal ultrastructural details previously too crowded or small to be efficiently resolved with conventional microscopy. A visual comparison of NHS ester bulk amine pan-staining of neurons that are non-expanded (**Fig. 2a**) or expanded with pan-ExM, (**Fig. 2d-k**, **Supplementary Videos 1 and 2**), confirms the validity of this approach: non-expanded synapses show essentially uniform staining, revealing little information, whereas expanding neurons ∼16-fold allows to spatially resolve synapses by their protein density patterns. Analogous to phosphotungstic acid- (PTA-) staining of neurons in EM (**Fig. 2l**) (Bloom and Aghajanian, 1966; Aghajanian and Bloom, 1967), now resolvable hallmark features such as dense projections (DP) of the presynaptic bouton (**Fig. 2m**, lime arrow) and the postsynaptic density (**Fig. 2m**, salmon arrow) allow for the identification of synapses by their morphological characteristics. We see spines that are stubby (**Fig. 2d,f,i**), mushroom shaped (**Fig. 2h,j**), and thin (**Fig. 2g,k**). Strikingly, we also observe hexagonal protein-dense patterns formed by presynaptic DPs (**Fig. 2e**), a discovery made in the 1960s (Gray, 1963; Pfenninger, 1972). A gallery of synapses imaged with pan-ExM is shown in **Supplementary Figure 1**.

**Figure 2:**
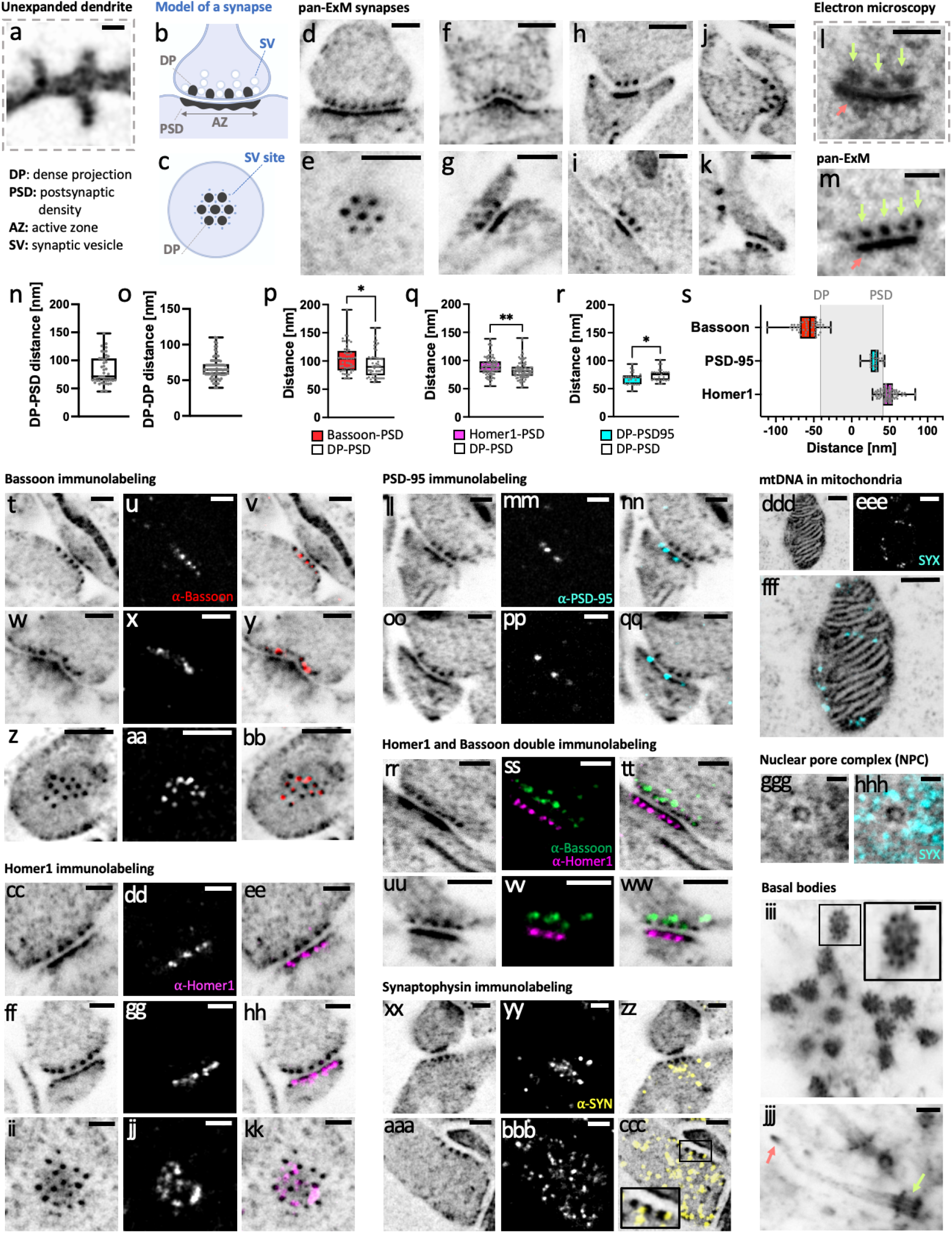
pan-ExM reveals synapse ultrastructure in dissociated neurons. **a**, NHS ester pan-stained dendrite in a non-expanded sample showing dendritic spines. **b**, axial view model of a synapse showing dense projections (DPs) and synaptic vesicles (SVs) in the presynaptic bouton, and the postsynaptic density (PSD), and the active zone (AZ) in the postsynaptic dendritic spine. **c**, top view of a synapse showing hexagonal dense projections (DPs) in the presynaptic bouton and synaptic vesicle (SV) attachment sites. **d**, **f-k**, pan-ExM processed and NHS ester pan-stained spines including mushroom (**h**, **j**), stubby (**d, f, i**), and thin (**g, k**) shapes. **e**, NHS ester pan-stained synapse showing hexagonally arranged DPs. **l**, transmission EM (TEM) image of a phosphotungstic acid (PTA) stained synapse showing prominent DPs (lime arrows) and a PSD (salmon arrow). **m**, pan-ExM processed and NHS ester pan-stained synapse for comparison, showing similar hallmark ultrastructural features. **n**, DP-PSD distances (n = 44 measurements from 4 independent experiments). **o**, DP-DP distances (n = 78 measurements from 6 independent experiments). **p**, Comparison of Bassoon-PSD and DP-PSD distances (n = 50 measurements from 3 independent samples). **q**, Comparison of Homer1-PSD and DP-PSD distances (n = 85 measurements from 4 independent samples). **r**, Comparison of PSD95-DP and DP-PSD distances (n = 25 measurements from 2 independent samples). *: p<0.05; **: p<0.01. **s**, Relative spatial distributions of Bassoon, PSD-95, and Homer1 along the trans-synaptic axis. **t**, **w**, **z**, axial (**t**, **w**) and top (**z**) views of synapses pan-stained with NHS ester. **u**, **x**, **aa**, Bassoon immunolabeling of the same areas. **v**, **y**, **bb**, respective overlays. **cc-kk**, **ll-qq**, **rr-ww**, and **xx-ccc**, same as **t-bb** in samples labeled for Homer1, labeled for PSD-95, double-labeled for Homer1 and Bassoon, and labeled for Synaptophysin (SYN), respectively. The inset in **ccc** shows SYN puncta, representing synaptic vesicles, intercalated between neighboring DPs. **ddd**, NHS ester image of a mitochondrion in a hippocampal rat neuron. **eee**, SYTOX Green (SYX) staining of the same area. **fff**, overlay. **ggg**, NHS ester image of a nuclear pore complex (NPC). **hhh**, overlay of **ggg** with a SYTOX Green image of the same area. **iii**, NHS ester image of basal bodies in a mouse neuron. The inset shows the familiar centriolar cartwheel structure. **jjj**, NHS ester image of a cilium in a mouse neuron. Lime and salmon arrows point to the basal body and ciliary tip, respectively. Gamma corrections: (**d, e**) γ=0.8; (**h**, **j**, **k**, **m**, **cc**, **ll**, **oo**, **rr**, **uu**, **xx**, **aaa**, **iii**, **jjj**) γ=0.7; (**i**) γ=0.6. **jjj** is a z-projection (intensity average) of 5 images. All scale bars are corrected for the expansion factor. Scale bars, (**a**) 800 nm, (**d-m**, **t-ccc**, **jjj**) 200 nm, (**ddd-fff**) 300 nm, (**ggg**, **hhh**) 50 nm, (**iii**) 100 nm.

To determine the achieved linear expansion factors, we imaged SYTOX Green-stained neuron nuclei in non-expanded samples and samples expanded using our standard protocol and compared the average nuclear cross-sectional area in both cases. On average, we obtained an expansion factor of 15.7 ± 0.3 (mean ± s.d.; N = 4 experiments; n = 6–13 nuclei per experiment). Dividing the measured distances between dense projections (DP) and the postsynaptic density (PSD) by the expansion factor determined from nuclei measurements, we obtained a value of 81.9 ± 25.8 nm (mean ± s.d.; N = 4 experiments; n = 44 synaptic profiles; **Fig. 2n**), which is consistent with the range of pre- and post-synaptic density distance measurements determined previously by super-resolution microscopy and electron tomography (Valtschanoff et al., 2001; Dani et al., 2010; Acuna et al., 2016; Hao et al., 2021; Yang and Annaert, 2021). Similarly, dividing the distance between neighboring DPs by the nuclear expansion factor, we obtained a value of 67.2 ± 15.4 nm (mean ± s.d.; N = 6 experiments; n = 78 DP profiles; **Fig. 2o**), consistent with earlier reports in EM (Harlow et al., 2001; Acuna et al., 2010). In all subsequent experiments, we used the DP-PSD distance as a metric for linear expansion factor calculation.

pan-ExM in dissociated neurons is compatible with immunofluorescence labeling as well as other established chemical stainings, enabling correlative studies which combine specific and contextual pan-staining approaches. Focusing on the synapse, **Figure 2t-ccc** shows the distributions of synaptic proteins Bassoon, Homer1, PSD-95, and Synaptophysin in the context of synaptic ultrastructure. We observe compartmentalization of active zone protein Bassoon into distinct puncta as dense projections (**Fig. 2t-bb**) with synaptic vesicle protein Synaptophysin intercalating in between neighboring dense projections (**Fig. 2xx-ccc**), supporting the model that DPs represent distinct sites for synaptic vesicle docking and fusion at the active zone. We also observe nanoclustering of postsynaptic density proteins Homer1 and PSD-95 along an otherwise macular and dense PSD, with Homer1 slightly offsetting the PSD further into the spine (**Fig. 2cc-hh,q**) and PSD-95 concentrating directly over the PSD (**Fig. 2ll-qq,r**), consistent with previous work (Xiao et al., 1998). The expansion-corrected distributions of Bassoon, Homer1, and PSD-95 within the axial DP-PSD distances are all in agreement with previously published studies (Dani et al., 2010; Hao et al., 2021). For instance, the distance between Bassoon and the PSD is 103.7 ± 24.4 nm (mean ± s.d.; N = 3 independent samples; n = 50 synaptic profiles; **Fig. 2p**); the distance between Homer1 and the DP is 82.86 ± 15.8 nm (mean ± s.d.; N = 4 independent samples; n = 85 synaptic profiles; **Fig. 2q**); and the distance between PSD-95 and the DP is 68.4 ± 11.0 nm (mean ± s.d.; N = 2 independent samples; n = 25 synaptic profiles; **Fig. 2r**). **Figure 2s** shows a plot of the positions of Bassoon, Homer1, and PSD-95 along the trans-synaptic axis defined as the center position between DP and PSD.

Double immunostainings are compatible with pan-ExM. **Figure 2rr-ww** and **Supplementary Video 3** show Bassoon-Homer1 immunofluorescence images at super-resolution and in their synaptic context. A gallery of images showing distributions of synaptic proteins Bassoon, Homer1, and PSD-95 is shown in **Supplementary Figs. 2-5**. This richness of information is inaccessible with conventional confocal microscopy of unexpanded or only ∼5-fold expanded samples (**Supplementary Fig. 6**).

Optimizing antibody labeling parameters to achieve efficient and background-free staining is critical for pan-ExM imaging. Because the hydrogels used in pan-ExM are entangled and dense (∼30% w/v monomer concentration), we suspect that many antibodies can become entrapped within this hydrogel matrix when used in high concentrations, often producing a granular background (**Supplementary Fig. 7b**). We found, for example, that lowering the concentrations of both Homer1 primary and dye-conjugated secondary antibodies from ∼4 µg/mL to ∼2 µg/mL strongly reduced the background without compromising signal levels (**Supplementary Fig. 7j**).

pan-ExM can clearly resolve subcellular structures in dissociated neurons that were previously inaccessible with standard confocal microscopy. **Figure 2ddd-fff** shows mtDNA inside a mitochondrion with visible cristae, **Fig. 2ggg-hhh** shows the hollow, circular structure of a nuclear pore complex, and **Fig. 2iii-jjj** reveals the cartwheel structure of basal bodies, their distal appendages, and the ciliary tip of a cilium. Furthermore, by combining NHS ester pan-staining with metabolic incorporation of palmitic acid azide, it becomes possible to examine the contact sites of membranous organelles, such as the tubules of the endoplasmic reticulum (ER) and mitochondria (**Supplementary Fig. 8**). A gallery of subcellular neuronal features is shown in **Supplementary Figs. 9-10**.

### pan-ExM-t reveals tissue ultrastructural features

Having established pan-ExM in dissociated neuron cultures, we adapted our technique to 70 µm-thick mouse brain tissue sections. Because of stark differences in thickness, lipid content, and presence of a highly connected extracellular matrix in tissue that is absent in dissociated neurons, brain fixation, sample denaturation, and antibody labeling parameters had to be optimized.

Ultrastructural analysis of brain tissue sections in EM has typically relied on formaldehyde (FA) and glutaraldehyde (GA) fixation to best preserve fine structures (Hayat, 1981; Sabatini et al., 1963). Because pan-ExM is an ultrastructural imaging method, structural preservation at the 10-nm scale is of utmost importance (M’Saad and Bewersorf, 2020). However, compatibility with immunolabeling is equally important to the core concept of our technique, and antigens are known to be masked by glutaraldehyde fixation (Migneault et al., 2004). To examine the effects of fixation and post-fixation on tissue preservation, we expanded 70 µm-thick mouse brain tissue sections that were fixed with 4% FA and not post-fixed (*Fix-1*), fixed with 4% FA and post-fixed with 0.7% FA + 1% acrylamide (AAm) (*Fix-2*), fixed with 4% FA and post-fixed with 4% FA + 20% AAm (*Fix-3*), fixed with 4% FA + 0.1% GA and post-fixed with 4% FA + 20% AAm (*Fix-4*), and fixed with 4% FA and post-treated with 0.1 mg/mL acryloyl-X SE (AcX) (*Fix-5*). **Supplementary Figs. 11-12** show that tissue treated with *Fix-1* has no distinguishable structural features beyond the outline of cell nuclei, suggesting little protein retention; tissue treated with *Fix-2* expands 18.5 ± 1.3-fold (mean ± s.d.; N = 6 fields of view; n = 38 synaptic profiles, **Supplementary Fig. 12p**) and shows resolved synapses and cell bodies, but no distinguishable neurites; tissue treated with *Fix-3* expands 16.0 ± 1.0-fold (mean ± s.d.; N = 9 fields of view; n = 81 synaptic profiles) and shows distinguishable neurites, but many detached cell bodies; tissue treated with *Fix-4* expands 11.6 ± 1.1-fold (mean ± s.d.; N = 8 fields of view; n = 77 synaptic profiles) and exhibits adequate neuropil and cell body preservation, and finally; tissue treated with *Fix-5* expands 11.8 ± 1.9-fold (mean ± s.d.; N = 9 fields of view; n = 74 synaptic profiles) and shows poor cell body preservation and multiple artifactual gaps in neuropil. These results suggest that the expansion factor increases with lower post-fixation strength and decreases with the addition of GA in the fixative. They also suggest that using AcX, a common acryloylated and amine-reactive reagent in ExM (Tillberg et al., 2016), results in lower expansion factors as well as multiple artifacts in tissue ultrastructure when coupled with nonenzymatic homogenization.

Moreover, to examine tissue preservation further, we determined the extracellular space and lipid membrane (ECS+) fraction of neuropil across the different fixation schemes presented. The ECS fraction in live organotypic brain slices, determined by STED microscopy, ranges from 5% to 36% (Tonnesen et al., 2018). We therefore expect the ECS+ fraction to be within that range or higher by ∼10% to account for unlabeled lipid membrane boundaries. **Supplementary Fig. 13** shows that indeed lowering the strength of the post-fixative (*Fix-2*) or using AcX (*Fix-5*) result in large ECS gaps (ECS+>60%) that are likely artifactual, while using stronger post-fixatives (*Fix-3* and *Fix-4*) result in an ECS+ fraction within an acceptable range (∼30%). It is worth noting that in EM of chemically fixed tissue, the ECS fraction is notoriously underestimated because of tissue shrinkage from excessive fixation (ECS shrinkage is ∼6 fold), giving the false notion that neurites are tightly apposed to one another (Harreveld et al., 1965; Korogod et al., 2015).

We conclude based on our assessment of fixation effects on ultrastructural preservation in expanded brain tissue that strong post-fixation with FA and AAm (*Fix-3* and *Fix-4*) is necessary for good neuropil and ECS preservation. However, these still cause artifactual gaps in neuropil, and in the case of FA+GA-fixed tissue, result in a low expansion factor of ∼11 fold. Therefore, we hypothesized that excessive interprotein crosslinking occurs in the initial fixation stage and predicted that transcardial perfusion with both FA and AAm along with post-fixation in the same fixative would result in a more uniform expansion. Inspired by CLARITY and MAP (Chung et al., 2013; Ku et al., 2016) where FA is combined with acrylamide to simultaneously quench inter-protein crosslinking and functionalize proteins with an AAm group, we used 4% FA + 20% AAm in both the transcardial perfusion and overnight post-fixation solutions (*Fix-6*). We obtained an expansion factor of 24.1± 1.4-fold (mean ± s.d.; N = 10 fields of view from 3 independent experiments; n = 254 synaptic profiles, **Fig. 3i**; **Supplementary Fig. 12p**) as well as good neuropil preservation, very little artifactual tissue perforations (**Fig. 3a-c**; **Supplementary Figs. 11-12**), and an acceptable ECS+ fraction of ∼38% (**Fig. 3j-l**; **Supplementary Fig. 13**). We therefore decided to use this fixation strategy for all subsequent experiments.

**Figure 3:**
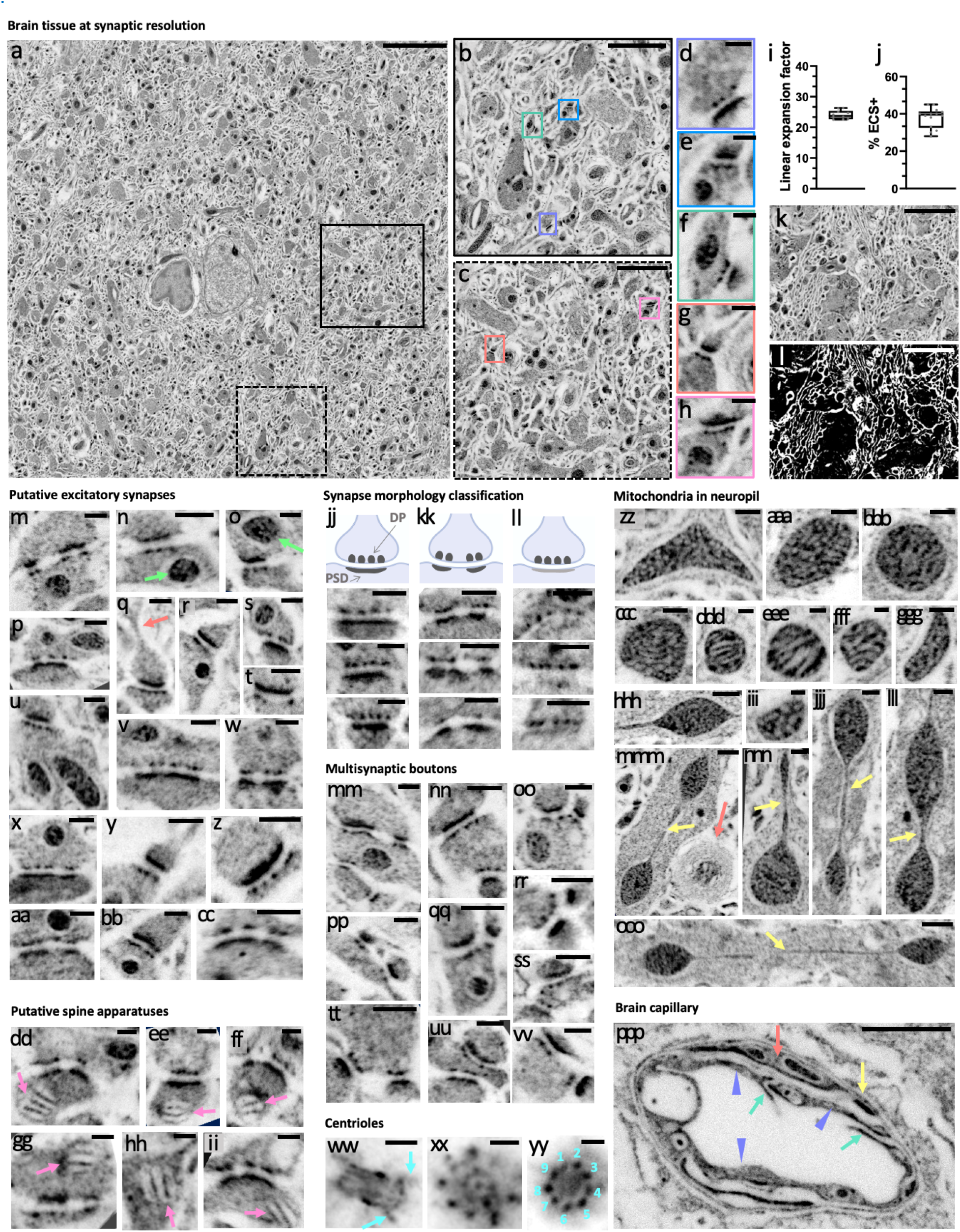
pan-ExM-t reveals mouse brain tissue ultrastructure. **a**, NHS ester pan-stained tissue section of the mouse cortex. **b**, **c**, magnified areas in the black and dotted black boxes in **a**, respectively. **d-h**, magnified areas identified by the correspondingly colored boxes in **b**, **c** showing putative excitatory synapses. **i**, linear expansion factor (n = 254 measurements from 3 independent experiments). **j**, ECS + lipid membrane (ECS+) fraction (n = 14 measurements from 3 independent experiments). **k**, image of neuropil in the hippocampus. **l**, same area as **k** where white pixels represent the ECS+. **m-cc**, putative excitatory synapses defined by a prominent PSD. Lime arrows in **n** and **p** point to mitochondria in the presynaptic bouton and in the postsynaptic compartment, respectively. The salmon arrow in **q** points to a spine neck. **dd-ii**, putative spine apparatuses (pink arrows) in the postsynaptic compartment defined by a characteristic lamellar arrangement. **jj-ll**, classification of synapses based on the patterns and intensity of the PSD; **jj**, class 1: the PSD is prominent and macular; **kk**, class 2: the PSD is prominent and perforated; **ll**, class 3: the PSD, unlike DPs, is barely visible. **mm-vv**, NHS ester pan-stained multisynaptic boutons and their postsynaptic partners. **ww**, lateral view of a centriole showing distal and proximal ends as well as distal appendages (turquoise arrows). **xx**, **yy**, top views of centrioles showing the cartwheel structure (**xx**) and the 9-fold symmetry of microtubule triplets (**yy**). **zz-ooo**, mitochondria with vesicular cristae (**zz-ccc**, **ggg**, **iii**), lamellar cristae (**ddd-fff**), and teardrop-shaped with tubular extensions (yellow arrows) (**mmm-ooo**). Salmon arrow in **mmm** points to putative myelinated sheaths. **ppp**, brain capillary showing endothelial cells (lavender arrow heads), putative tight junctions (TJs; teal arrows) that link neighboring endothelial cells, putative pericyte branch (salmon arrow), and the basement membrane (BM; yellow arrow). Gamma corrections: (**a**, **ww-yy**, **ppp**) γ=0.7; (**m-vv**, **zz-ooo**) γ=0.8. All scale bars are corrected for the expansion factor. Scale bars, (**a**) 5 µm, (**b**, **c**, **ppp**) 1 µm, (**d-h**, **m-cc**, **jj-vv**, **xx**, **ww**, **zz-ccc**, **ggg**, **jjj**, **lll**) 200 nm, (**k**, **l**) 3 µm, (**dd-ii**, **xx**, **yy**, **ddd-fff**, **iii**, **nnn**) 100 nm, (**hhh**, **mmm**, **ooo**) 400 nm.

In the process of optimizing protein denaturation conditions, we found that increasing the denaturation temperature above 90 °C or using alkaline buffers resulted in hydrogel disintegration, possibly because of the lability of the DHEBA crosslinker molecule (Brady and O’Connell, 1976). We therefore adopted the pH 6.8 SDS buffer of pan-ExM and used 75 °C as the temperature for denaturation, similar to 73 °C used in the original protocol (M’Saad and Bewersdorf, 2020). We also found that tissue-hydrogel hybrids denatured for 4 h were macroscopically flat, while gels denatured for fewer than 4 h exhibited crumbled macrostructures, signifying incomplete mechanical homogenization. We therefore assessed denaturation times of 4 h (*Denat-4*), 6 h (*Denat-6*), and 8 h (*Denat-8*) for both expansion factor and relative protein retention (measured by reporting peak DP intensity). We found that *Denat-4* results in an expansion factor of 26.9 ± 1.2 (mean ± s.d.; N = 3 fields of view; n = 50 synaptic profiles, **Supplementary Fig. 14j**) and DP peak intensity of 1506.8 ± 243.8 (mean ± s.d.; N = 3 fields of view; n = 50 intensity measurements, **Supplementary Fig. 14k**), *Denat-6* yields a similar expansion factor of 26.0 ± 1.3 (mean ± s.d.; N = 3 fields of view; n = 42 synaptic profiles) and slightly lower DP peak intensity of 1353.3 ± 248.3 (mean ± s.d.; N = 3 fields of view; n = 42 intensity measurements), and finally *Denat-8* shows an increase in expansion factor to 33.7 ± 1.6 (mean ± s.d.; N = 3 fields of view; n = 100 synaptic profiles) and partial protein loss with DP peak intensity of 1014.3 ± 209.1 (mean ± s.d.; N = 2 fields of view; n = 48 intensity measurements). Since we prioritize protein retention and argue that expansion factors of ∼24 fold are sufficient to resolve ultrastructural features of interest, we denatured our samples for 4 h at 75°C in all subsequent experiments.

Equipped with a pan-ExM-t protocol that preserves ultrastructure well and allows for ∼24-fold linear expansion, we imaged a wide variety of tissue nanostructures across both hippocampal and cortical regions of the mouse brain. **Figure 3a** shows a tiled ∼1x1 mm^2^ image (corresponding to ∼42x42 µm^2^ after correction for the expansion factor) of NHS ester pan-stained cortical tissue at synaptic resolution. Hallmark synaptic features such as pre- and postsynaptic densities can now be imaged and with a standard confocal microscope deep inside a brain tissue section (**Fig. 3d-h**; **Supplementary Figs. 15-17**; **Supplementary Videos 4** and **5**). For example, we could observe that putative excitatory synapses, defined by a prominent PSD (**Fig. 3m-cc**), often featured mitochondria in the vicinity of axonal boutons (**Fig. 3n**, arrow) and sometimes in the postsynaptic partner (**Fig. 3o**, arrow). Intriguingly, we can discern densely stained stacked structures in some postsynaptic compartments, suggestive of spine apparatuses (**Fig. 3dd-ii**) (Harris and Weinberg, 2012). Moreover, the obtained resolution allowed us to classify synapses based on variations in PSD patterns and intensities into three classes inspired by the previously established (Gray, 1959; Klemann and Roubos, 2011; Harris and Weinberg, 2012; Nahirne and Trembley, 2021) Gray classification of synapses: Class 1, where the PSD is dense and macular (**Fig. 3jj**); Class 2, where the PSD is dense and perforated (**Fig. 3kk**); and Class 3, where the PSD is barely visible, despite prominent DPs in the presynaptic bouton (**Fig. 3ll**). Furthermore, pan-ExM-t images clearly reveal the spine neck (**Fig. 3q**, arrow) and multisynaptic boutons **(Fig. 3mm-vv**).

Centrioles in the perikarya are distinguished by a bright NHS ester pan-staining showing clear distal appendages (**Fig. 3ww**) and, in top views, the cartwheel structure (**Fig. 3xx**) and nine-fold symmetry (**Fig. 3zz**). The nine-fold symmetry is also visible in multiciliated ependymal epithelia lining the lateral ventricles of the mouse brain (**Fig. 5l**, **Supplementary Video 6**, **Supplementary Fig. 18**). Moreover, we also observe that mitochondria morphologies vary strongly across neuropil (**Fig. 3zz-ooo**), some featuring clearly resolvable lamellar cristae, and shapes ranging from vesicular to teardrops with tubular extensions (**Fig. 3mmm-ooo**, arrows), reported to correlate with disease and synaptic performance (Zhang et al., 2016; Cserép et al., 2018). A closer look at brain capillaries (**Fig. 3ooo**, **Supplementary Fig. 19**) reveals clearly discernible endothelial cells, tight junctions, pericytes, and the basement membrane (Nahrine and Trembley, 2021).

### pan-ExM-t is compatible with antibody labeling of synapses in thick brain tissue sections

A particular strength of pan-ExM-t is its ability to localize specific proteins to sub-compartments revealed in the contextual pan-stained channel. Our protocol must therefore allow for efficient antibody staining, while also maintaining good ultrastructural preservation. Because hydrogels with high monomer concentrations are known to impede antibody diffusion, we investigated whether lowering the monomer concentration in the different interpenetrating hydrogels would result in better signal-to-noise antibody stainings. We assessed both antibody-labeling efficiency and ultrastructural preservation in hydrogels synthesized with different sodium acrylate (SA) monomer concentrations. We hypothesized that lowering this concentration would allow for larger effective hydrogel mesh sizes and therefore better antibody penetration. In summary, we used four different hydrogel monomer combinations with either 19% (w/v) or 9% (w/v) SA in the first and second expansion hydrogels, referred to as *19SA/19SA*, *19SA/9SA*, *9SA/19SA*, and *9SA/9SA*. Comparing expansion factors, we found that *19SA/19SA* gels expand as we previously reported 24.1± 1.4-fold (mean ± s.d.; N = 10 fields of view from 3 independent experiments; n = 254 synaptic profiles, **Supplementary Fig. 20e**), *19SA/9SA* gels expand 20.3± 1.2-fold (mean ± s.d.; N = 12 fields of view from 2 independent experiments; n = 267 synaptic profiles), *9SA/19SA* gels expand 20.5± 1.8-fold (mean ± s.d.; N = 6 fields of view from 2 independent experiments; n = 88 synaptic profiles), and *9SA/9SA* gels expand 18.5± 1.7-fold (mean ± s.d.; N = 6 fields of view from 2 independent experiments; n = 86 synaptic profiles). While expansion factors are not radically affected by variations in SA monomer concentrations, we noticed visible ultrastructural differences. Judging tissue preservation by the frequency of gaps in neuropil and detachments of cell bodies from the underlying tissue, we observe equally good tissue integrity in *19SA/19SA*, *19SA/9SA*, and *9SA/19SA* gels (**Supplementary Fig. 20a-c)**, but significantly distorted neuropil and dissociated cell bodies in *9SA/9SA* gels (**Supplementary Fig. 20d**). We suspect that this distortion effect is due to a decrease in hydrogel mechanical sturdiness when hydrogels of lower monomer concentrations, like the ones in ExR (Sarkar et al., 2020), are used. We therefore caution against synthesizing hydrogels that are mechanically too brittle to properly preserve the integrity of biological structures. As for antibody-staining efficiency, we observed equally efficient labeling in *19SA/19SA*, *19SA/9SA*, and *9SA/19SA* hydrogels for most antigens, with reduced background following synaptic protein immunolabeling in *19SA/9SA* hydrogels. Therefore, in all subsequent immunoassays, we used *19SA/9SA* gels for synaptic protein immunolabeling and *19SA/19SA* gels in immunolabeling structural markers.

With a now robust pan-ExM-t protocol capable of localizing specific proteins in their tissue ultrastructural context, we performed several immunostainings against commonly studied synaptic targets. **Figure 4** shows the distributions of synaptic proteins Homer1, Bassoon, PSD-95 (**Supplementary Video 6**), and Synaptophysin in the context of tissue and synaptic ultrastructure. The expansion-corrected distributions of Homer1, PSD-95, and Bassoon within the axial DP-PSD distances are all in agreement with published values as well as with values obtained from cultured neuron measurements in this work. For example, the distance between Bassoon and the PSD is 93.7 ± 16.9 nm (mean ± s.d.; N = 3 fields of view; n = 74 synaptic profiles; **Fig. 4ooo**); the distance between Homer1 and DP is 93.3 ± 16.1 nm (mean ± s.d.; N = 5 fields on views; n = 120 synaptic profiles; **Fig. 4ppp**); and the distance between PSD-95 and the DP is 76.4 ± 11.8 nm (mean ± s.d.; N = 3 fields of view; n = 113 synaptic profiles, **Fig. 4qqq**). **Figure 4rrr** shows a plot of the axial positions of Homer1, PSD-95, and Bassoon along the trans-synaptic axis defined as the center position between DP and PSD. In agreement with previous work, Homer1 is on average localized further inside the spine and away from the cleft than the PSD center, while PSD-95 is concentrated directly onto the PSD (Xiao et al., 1998). Interestingly, in Class 3 synapses, defined by the absence of a protein-dense PSD, we found that while Bassoon scaffold protein is present (**Figs. 4q-s**) PSD proteins Homer1 (**Figs. 4jj-ll**) and PSD-95 (**Figs. 4ddd-fff)** are not, reminiscent of inhibitory synapses.

**Figure 4:**
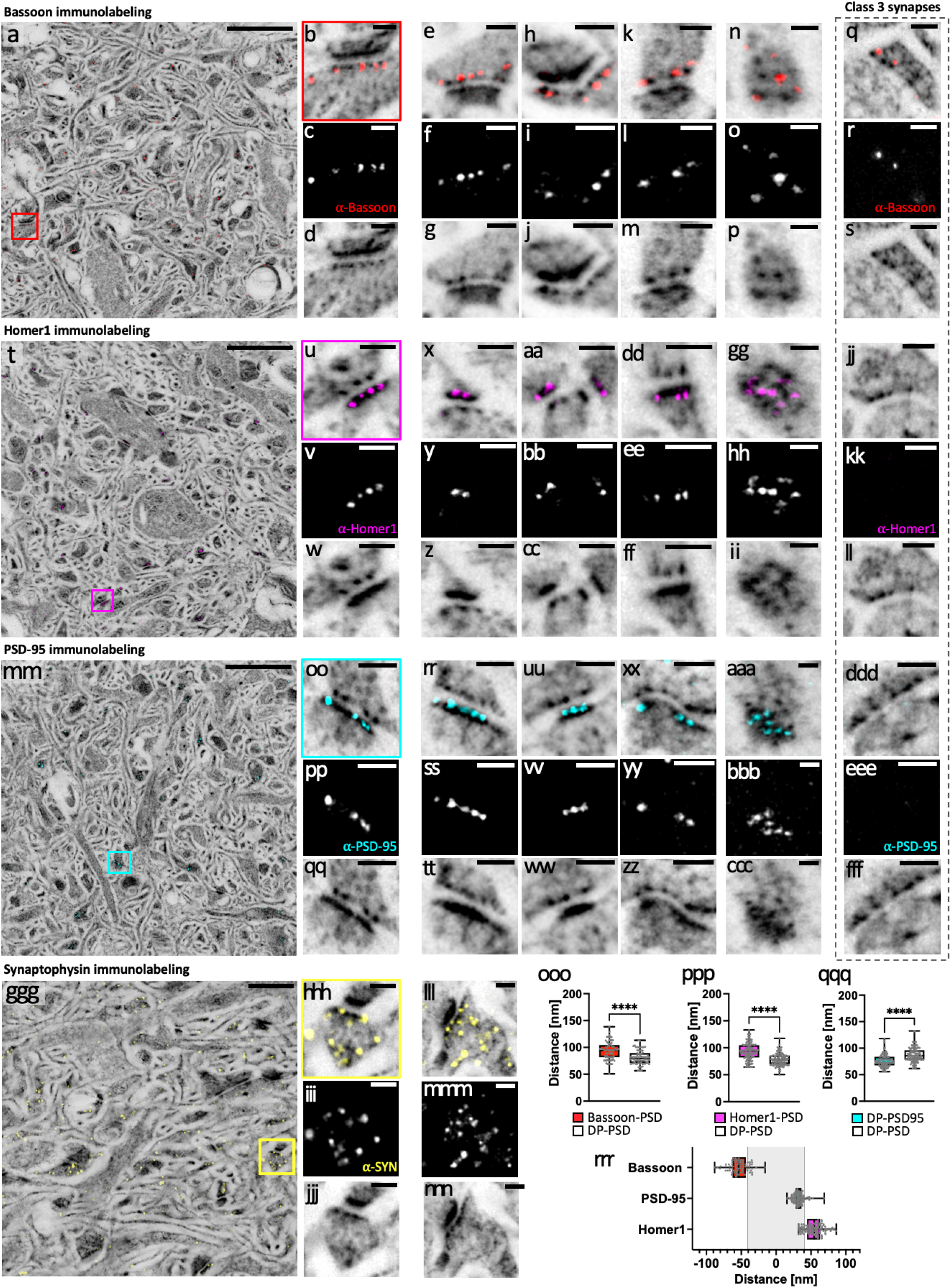
pan-ExM-t is compatible with antibody labeling of synaptic proteins in brain tissue. **a**, NHS ester (grayscale) pan-stained and Bassoon (red) immunolabeled brain tissue section. **b**, magnified area in the red box in **a** showing a synapse. **c**, Bassoon immunolabeling of the same area as in **b**. **d**, NHS ester pan-staining of the same area as in **b**. **e-p**, additional examples similar to **b-d** showing lateral (**e-m**) and top (**n-p**) views of Bassoon-immunolabeled synapses. **q-s**, lateral view of a Class 3 synapse pan-stained with NHS ester and immunolabeled with Bassoon antibody indicating that Class 3 synapses are positive for Bassoon protein. **t**, NHS ester (grayscale) pan-stained and Homer1 (magenta) immunolabeled brain tissue section. **u**, magnified area in the magenta box in **t** showing a synapse. **v**, Homer1 immunolabeling of the same area as in **u**. **w**, NHS ester pan-staining of the same area as in **u**. **x-ii**, additional examples similar to **u-w** showing lateral (**x-ff**) and top (**gg-ii**) views of Homer1-immunolabeled synapses. **jj-ll**, lateral view of a Class 3 synapse pan-stained with NHS ester and immunolabeled with Homer1 antibody indicating that Class 3 synapses are negative for Homer1 protein. **mm**, NHS ester (grayscale) pan-stained and PSD-95 (cyan) immunolabeled brain tissue section. **oo**, magnified area in the cyan box in **mm** showing a synapse. **pp**, PSD-95 immunolabeling of the same area as in **oo**. **qq**, NHS ester pan-staining of the same area as in **oo**. **rr-ccc**, additional examples similar to **oo-qq** showing lateral (**rr-zz**) and top (**aaa-ccc**) views of PSD-95-immunolabeled synapses. **ddd-fff**, lateral view of a Class 3 synapse pan-stained with NHS ester and immunolabeled with PSD-95 antibody indicating that Class 3 synapses are negative for PSD-95 protein. **ggg**, NHS ester (grayscale) pan-stained and Synaptophysin (SYN; yellow) immunolabeled brain tissue section. **hhh**, magnified area in the yellow box in **ggg** showing a synapse. **iii**, Synaptophysin immunolabeling of the same area as in **hhh**. **jjj**, NHS ester pan-staining of the same area as in **hhh**. **lll-nnn**, additional example similar to **hhh-jjj**. **ooo**, Comparison between Bassoon-PSD and DP-PSD distances (n = 74 measurements from 3 fields of view (FOVs) in one independent experiment). **ppp**, Comparison between Homer1-PSD and DP-PSD distances (n = 120 measurements from 5 FOVs in 2 independent experiments). **qqq**, Comparison between PSD95-DP and DP-PSD distances (n = 113 measurements from 3 FOVs in one independent experiment). ****: p<0.0001. **rrr**, relative spatial distributions of Bassoon, PSD-95, and Homer1 along the trans-synaptic axis. All images (with the exception of **ggg-nnn**) are z-projections (intensity average) of 2 images. All NHS ester images were Gamma-corrected with γ=0.7 with the exception of the Synaptophysin images (γ=0.6). All scale bars are corrected for the expansion factor. Scale bars (**a**, **t**, **mm**) 2 µm, (**b-m**, **q-s**, **u-ff**, **jj-ll**, **oo-zz**, **ddd-fff**, **hhh-nnn**) 200 nm, (**n-p**, **gg-ii**, **aaa-ccc**) 100 nm, (**ggg**) 1 µm.

### pan-ExM-t is compatible with antibody labeling of tissue structures

pan-ExM-t is not limited to providing context to synaptic proteins. We found that SYTOX Green produces a bright nuclear staining that overlays perfectly with NHS ester pan-stained cell nuclei (**Fig. 5a-d**). Furthermore, we can resolve individual glial fibrillary acidic protein (GFAP) filaments in astrocytes (**Fig. 5e-k**) including near multiciliated ependymal epithelia (**Supplementary Fig. 18**) and engulfing blood vessels (**Fig. 5k; Supplementary Video 7**). Interestingly, the distinctive pan-stain pattern of astrocyte nuclei and their thick and fibrous cytoplasmic branches allow for their identification without GFAP, analogous to heavy metal-stains in EM (**Supplementary Fig. 21**; **Supplementary Video 8**) (Luse, 1956; Nahirne and Tremblay, 2021). We also observe that myelin basic protein (MBP-) labeled structures are devoid of surrounding pan-staining, suggesting these represent myelinated axons (**Fig. 5m-s**). Moreover, when we immunolabeled GFP in Thy1-GFP transgenic mice, we noticed the sparsity of this marker in both neuropil and neuron somas (**Fig. 5t-ii**; **Supplementary Video 9**). Originally designed to resolve individual dendrites and axons in densely packed neuropil (Feng et al., 2000), GFP-Thy1 imaged with pan-ExM-t reveals these transgenic neurons in their ultrastructural context. pan-ExM-t clearly shows the GFP surrounding, but being excluded from, mitochondria (**Fig. 5v-x**). The ability to distinguish GFP-positive dendritic spine heads (**Fig. 5y-aa**) and axonal boutons (**Fig. 5bb-gg**) from their synaptic partners suggests that pan-ExM-t is especially suited for light-based neural tracing and connectomics.

**Figure 5:**
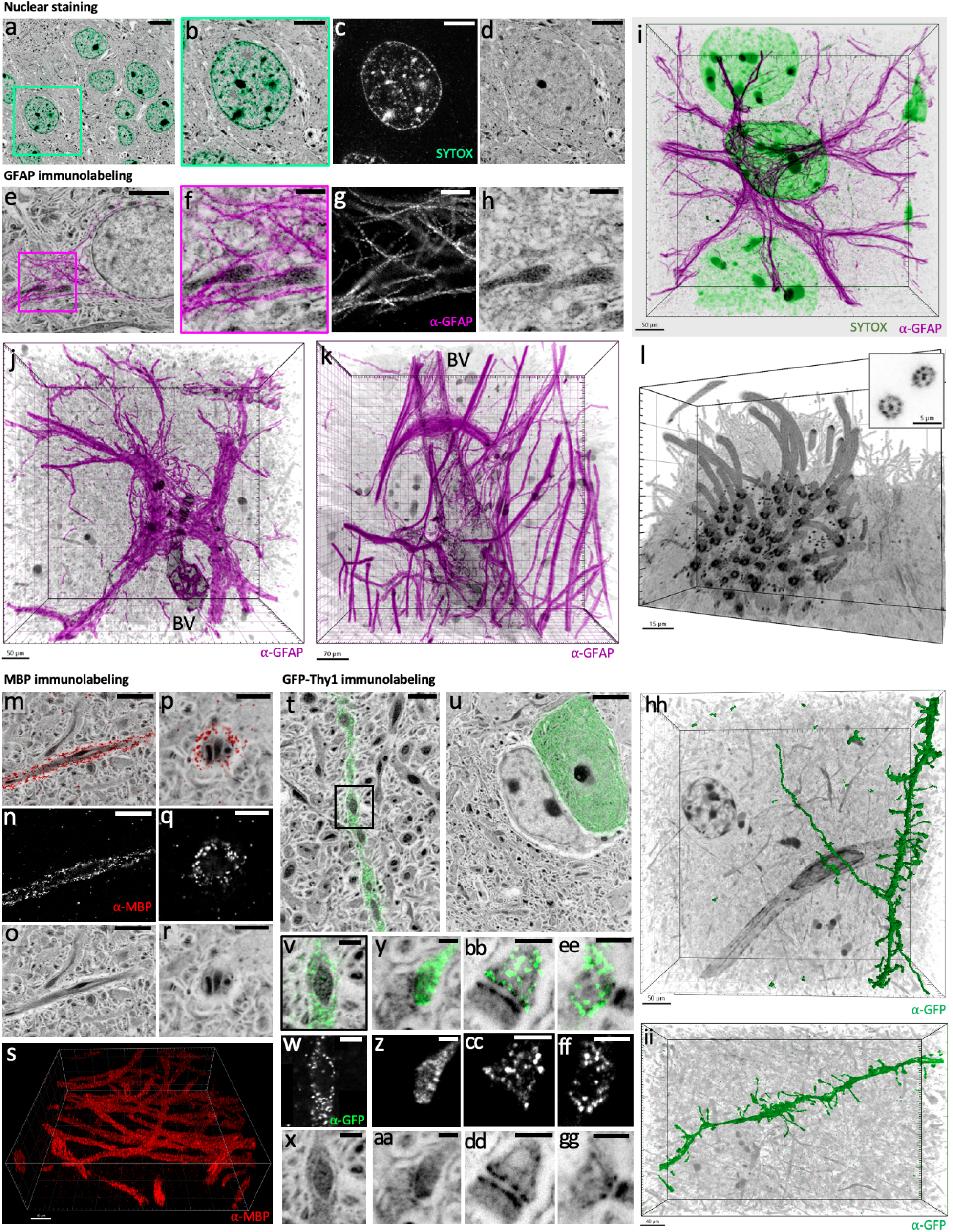
pan-ExM-t is compatible with antibody labeling of brain tissue structural markers. **a**, cortical neurons pan-stained with NHS ester (grayscale) and stained with SYTOX green (teal) showing a neuron nucleus. **b-d**, magnified view of the area in the green box in **a** and the SYTOX Green (**c**) and NHS ester (**d**) channels shown separately. **e**, astrocyte pan-stained with NHS ester (grayscale) and immunolabeled with glial fibrillary acidic protein antibody (⍺-GFAP; magenta). **f-h**, magnified view of the area in the magenta box in **e** showing GFAP filaments surrounding mitochondria, and the ⍺-GFAP (**g**) and NHS ester (**h**) channels shown separately. **i**, 3D rendering of ⍺-GFAP (magenta), SYTOX Green (green), and NHS ester (grayscale). **j**, **k**, 3D renderings of ⍺-GFAP (magenta) and blood vessels (BV) pan-stained with NHS ester (grayscale). **l**, multiciliated ependymal epithelia pan-stained with NHS ester (grayscale). Inset shows 9-fold symmetry of cilia. **m-r**, lateral (**m-o**) and top (**p-r**) views of axons pan-stained with NHS ester (grayscale) and immunolabeled with myelin basic protein antibody (⍺-MBP; red). ⍺-MBP (**n**, **q**) and NHS ester (**o**, **r**) channels of the areas shown in **m** and **p**. **s**, 3D rendering of anti-MBP showing multiple axons. **t**, **u**, neuropil pan-stained with NHS ester (grayscale) and anti-GFP (green) showing that only a sparse subset of cell bodies and neurites express GFP-Thy1. **v-x**, magnified area in the black box in **t** showing a neurite pan-stained with NHS ester (**x**) and immunolabeled with anti-GFP (**w**) where GFP-Thy1 is not expressed inside mitochondria. **y-gg**, a dendritic spine (**y-aa**) and axonal boutons (**bb-gg**) pan-stained with NHS ester and immunolabeled with anti-GFP. **z**, **cc**, **ff**, anti-GFP channels of the areas shown in **y**, **bb**, and **ee**, respectively. **aa**, **dd**, **gg**, NHS ester channels of the areas shown in **y**, **bb**, and **ee**, respectively. **hh**, 3D representation of a surface-rendered dendrite immunolabeled with anti-GFP (green) and NHS ester pan-staining (grayscale). **ii**, 3D rendering of a dendrite immunolabeled with anti-GFP (green) and NHS ester pan-staining (grayscale). Gamma corrections: (**a**, **e**, **m**, **p**, **t**, **u**, **x**, **aa**, **dd**, **gg**) γ=0.7. 3D renderings were processed with Imaris with gamma corrections applied. 3D image processing details for images **i-l**, **s**, **hh**, and **ii** are found in the **Methods** section. Scale bars in the 3D renderings **i-l**, **s**, **hh**, and **ii** are not corrected for the expansion factor. All other scale bars are corrected for the expansion factor. Scale bars (**a**) 5 µm, (**b-d**, **e, m-o**, **u**) 2 µm, (**f-h**, **v-x**) 500 nm, (**p-r**, **t**) 1 µm, (**y-gg**) 200 nm.

### pacSph enables lipid labeling of tissue in pan-ExM-t

Neural connectomics analysis in EM generally relies on contrast from lipid membranes to delineate cellular boundaries. One way to achieve this in pan-ExM-t would be to preserve the lipid content. Since there are roughly 5 million lipids per 1 μm^2^ of membrane surface and 150-times more lipids than proteins on a typical cell membrane (Quinn et al., 1984), we expect that even following detergent extraction with SDS, a fraction of these lipids would remain in the hydrogel. Interestingly, when we labeled SDS-denatured and ∼4-fold expanded brain tissue with BODIPY TR Methyl Ester (BDP), we achieved a distinct pan-staining (**Supplementary Fig. 22**). Here, BDP highlights the nuclear envelope, organelles like mitochondria, the Golgi complex, the ER, as well as myelinated axons. All these structures resemble those shown in mExM (Karagiannis et al., 2019), but were acquired without using any specialized probes. Overlaid with sulfonated NHS ester dye pan-staining, we observe a striking differential hydrophilic-lipophilic pan-staining pattern (**Supplementary Fig. 23**), with significantly higher BDP staining levels achieved in tissue fixed with 4% FA + 0.1% GA than in tissue fixed with 4% FA (**Supplementary Fig. 24**). Similarly, in pan-ExM-t expanded tissue, we achieve distinctive and bright BDP labeling in tissue fixed with 4% FA + 0.1% GA (**Supplementary Fig. 25**), but almost no staining in tissue fixed with 4% FA. We also see major structural artifacts when supplementing the fixatives with 0.01% osmium tetroxide, known to better preserve lipids (Palade, 1952; Litman and Barrnett, 1972) (**Supplementary Fig. 24s-aa**; **Supplementary Fig. 26**). These results confirm that GA plays a major role in stabilizing and crosslinking the lipid content (Hayat, 1981), and suggest that monoaldehydes like FA alone do not provide adequate lipid preservation when preceded by detergent extraction.

Next, we asked whether replacing the anionic detergent SDS in our denaturation buffer with the chaotropic reagent guanidine hydrochloride (G-HCl) would better preserve the lipid content and enable superior lipid staining in 4% FA + 0.1% GA-fixed, pan-ExM-t expanded tissue. Unlike SDS, hyperhydration with urea or guanidine HCl is known to disrupt hydrogen bonding in proteins without solubilizing lipids (Huerta-Viga and Woutersen, 2013). EM images of brain tissue samples fixed with only 4% FA and chemically cleared with the functionally similar chaotropic molecule urea show intact membrane ultrastructure and up to 70% lipid retention (Hama et al., 2015; Oz et al., 2021). We chose to work with G-HCl and not urea, because the latter is known to decompose and carbamylate proteins at the elevated temperatures required for efficient denaturation (Poulsen et al., 2013). **Supplementary Fig. 27** shows that BDP signal is in fact higher in G-HCl-denatured neuron and tissue samples, giving credence to this approach. However, our data also show that BDP does not exclusively stain cell membranes and subjecting samples to no denaturation at all results in lower BPD signal. This observation is more evident in ∼20-fold expanded samples where easily distinguishable neurite boundaries are not specifically stained with BDP (**Supplementary Fig. 28**). All these results suggest that lipophilic dyes like BDP do not exclusively label lipid membranes in denatured samples, but also label the hydrophobic core of proteins exposed following denaturation. In fact, in thermal denaturation assays, environmentally sensitive fluorescent dyes that bind to the hydrophobic core of proteins are commonly used to determine the temperature at which proteins denature (e.g., in differential scanning fluorimetry (DSF)) (Senisterra et al., 2012; Vedadi et al., 2006; Hofman et al., 2016). Protein denaturation, however, is a necessary step in enzyme-free ExM methods, since electrostatic interactions between proteins must be nulled for isotropic sample expansion.

Since strategies to preserve the lipid content for post-homogenization lipophilic labeling do not result in exclusive cell membrane delineation and require antigen-masking GA fixation (Passagot et al., 1987), we decided to capitalize on pre-denaturation lipid labeling strategies. Here, proteins are not unfolded, and the lipid content may require less harsh fixation since we do not need to preserve it after the homogenization process. To label lipid membranes in ExM, all approaches have so far used trifunctional lipid probes in either live or fixed samples. In these probes, one functional group is a membrane-intercalating lipid, the second is hydrogel-anchorable, and the third is a reporter molecule. In the live-labeling approaches, cells have been labeled with acryolated and fluorescent 1,2-distearoyl-sn-glycero-3-phosphoethanolamine (DSPE) (Wen et al., 2020), clickable palmitic acid (M’Saad and Bewersdorf, 2020), aminated α-NH2-ω-N3-C6-ceramide (Götz et al., 2020), or clickable choline (Sun et al., 2020). The only strategy available to stain lipids in tissue with ExM after it is fixed is to use aminated (poly-lysine) membrane-intercalating probes such as pG5kb (Karagiannis et al., 2019) and mCling (Damstra et al., 2022; Revelo et al., 2014). These survive detergent disruption by covalent attachment to the hydrogel matrix and contain a biotin group available for conjugation with dye-labeled streptavidin after expansion. When we tested mCling in 5-fold expanded brain tissue sections, we found that although it labeled tissue hydrophobic components, this labeling was limited to the tissue surface, likely because of diffusion limitations (**Supplementary Fig. 29**). When we tested mCling in pan-ExM-t samples, we observed little to no labeling. mCling (Revelo et al., 2014) as well as an ExM-optimized version of it, pG5kb (Karagiannis et al., 2019), have 5 to 7 lysine groups making them relatively highly charged and bulky (**Supplementary Fig. 30a-b**). mCling and pG5kb also require that the tissue is fixed with GA, which we previously found decreases synaptic protein antigenicity as well as limits the expansion factor to ∼12 fold, and so we did not explore this avenue any further.

We therefore developed a new pan-ExM-t lipid labeling strategy where we use a commercially available photocrosslinkable and clickable sphingosine pacSph (Haberkant et al., 2015) as a lipid intercalating reagent in tissue fixed only with 4% FA + 20% AAm (to preserve antigenicity) (**Fig. 6n**). In our protocol, we (1) label 70 µm-thick tissue sections with pacSph prior to hydrogel embedding; (2) irradiate the sample with UV light (365 nm) to photo-crosslink the probes onto neighboring proteins (from both directions to compensate for depth-dependent UV light absorption in the 70-µM thick tissue); and (3) probe the ligands with fluorescent dyes after homogenization and expansion. It is worth noting that pacSph is smaller (335 Da) than mCling (1445 Da) or pG5kb (1158 Da), polar, and partially positively charged (amine pKa ∼6.6), allowing for high probe packing and sampling. Sphingosines are also membrane bilayer-rigidifying: they stabilize gel domains in membranes, raise their melting temperature, and induce membrane permeabilization without micellar extraction of proteins or lipids (Contreras et al., 2006; Jiménez-Rojo et al., 2014). We therefore expect sphingosine to diffuse efficiently in thick tissue. Furthermore, pacSph has a photoactivatable diazirine group that forms reactive carbene intermediates capable of insertion in C-H or N-H, and O-H bonds of tissue components upon UV irradiation (Gomes and Gozzo, 2010; Krieg et al., 1986). Since diazirine is on the lipid chain (**Fig. 6n**, pink diazirine), we expect that two pacSph lipids located on opposing sides of a membrane bilayer will photo-crosslink the embedded hydrogel polymer chains at the center of the hydrophobic phase of the bilayer, minimizing the linker error. This error is inevitable if we were to photo-crosslink the probes at the polar head group. Finally, retained lipids are conjugated with a hydrophobic azido dye at the lipid tail extremity (**Fig. 6n**, green alkyne). We believe all these factors make pacSph ideally suited for membrane delineation in pan-ExM-t.

**Figure 6:**
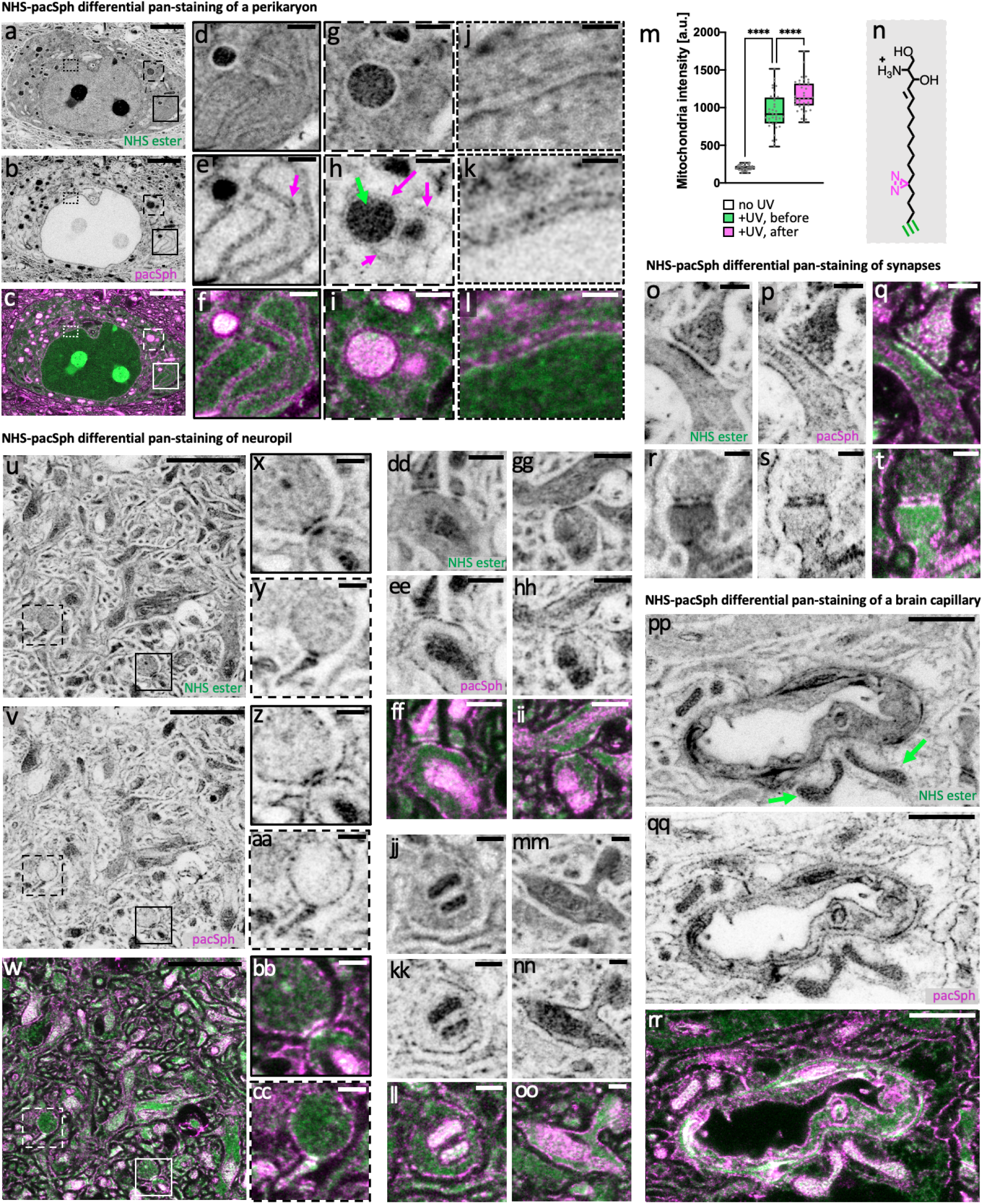
pacSph enables lipid membrane pan-staining of brain tissue in pan-ExM-t. **a-c**, perikaryon pan-stained both with NHS ester (**a**) and pacSph (**b**) (overlay shown in **c**). **d-l**, magnified areas of the solid boxes in **a-c** showing the two sides of ER tubules (magenta arrow; **d-f**), the dashed boxes in **a-c** showing a mitochondrion (green arrow) in contact with the ER (magenta arrow; **g-i**), and the dotted boxes in **a-c** showing the nuclear envelope (**j-l**). **m**, mitochondria signal levels in tissue pan-stained with pacSph and not irradiated with UV (white; n = 35 measurements from 5 FOVs in one independent experiment); pan-stained with pacSph and irradiated with UV before hydrogel embedding (green; n = 37 measurements from 5 FOVs in one independent experiment); and pan-stained with pacSph and irradiated with UV after hydrogel embedding (pink; n = 39 measurements from 5 FOVs in one independent experiment). ****: p<0.0001. **n**, chemical structure of pacSph. The probe is photocrosslinked via its diazirine group (magenta) and labeled via its alkyne group (green). **o-t**, synapses pan-stained both with NHS ester (**o, r**) and pacSph (**p, s**) (overlays shown in **q, t**).**u-w**, neuropil pan-stained both with NHS ester (**u**) and pacSph (**v**) (overlay shown in **w**). **x-cc**, magnified areas of the solid (**x, z, bb**) and dashed (**y, aa, cc**) boxes in **u-w** showing synapses. **dd-oo**, neurites and synapses pan-stained both with NHS ester (**dd, gg, jj, mm**) and pacSph (**ee**, **hh**, **kk**, **nn**) (overlays shown in **ff**, **ii**, **ll**, **oo**). **pp-rr**, brain capillary pan-stained both with NHS ester (**pp**) and pacSph (**qq**) (overlay shown in **rr**). Green arrows point to mitochondria. Gamma corrections: (**a**) γ=0.7; (**p**, **r**, **u**, **dd**, **gg**, **jj**, **mm**, **pp**) γ=0.5. All scale bars are corrected for the expansion factor. Scale bars (**a-c**) 3 µm, (**d-i**, **dd-ii**) 500 nm, (**u-w**) 2 µm, (**j-l**, **o-t**, **x-cc**, **jj-oo**) 250 nm, (**pp-rr**) 1 µm.

**Figure 6** confirms the strength of our approach. In the perikarya of brain tissue fixed with 4% FA + 20% AAm and pan-stained with pacSph, we can clearly discern both sides of ER tubules (**Fig. 6d-f**), ER and mitochondria contact sites (**Fig. 6g-i**), and both sides of the nuclear envelope (**Fig. 6j-l**). In neuropil, lipid membrane boundaries are now discernable **(Fig. 6o-oo)**, revealing the boundaries of neurites and synaptic compartments. In brain capillaries, the different membranes are also differentially highlighted relative to NHS ester pan-staining (**Fig. 6pp-rr**). Moreover, when we experimented with irradiating pacSph pan-stained tissue with UV light before or after the first hydrogel embedding, we found that UV irradiation after hydrogel embedding results in significantly higher probe retention, suggesting that diazirine is being photo-crosslinked to the dense hydrogel mesh in addition to surrounding proteins (**Fig. 6m**; **Supplementary Fig. 31**). In fact, a recent study showed that diazirine has an affinity for reacting with proteins containing large negative electrostatic surfaces in cells (e.g., carboxyl groups) (West et al., 2021). We suspect that diazirine in our probe is preferentially photo-crosslinked to carboxyl groups on the poly(acrylamide/acrylate) mesh, offering better retention of the lipid content. While benefiting from future investigation, our data shows that photo-crosslinking lipid-intercalating ceramides to the expansion hydrogel represents an important strategy in labeling lipids in ∼24-fold expanded and formaldehyde fixed brain tissue sections. A gallery of pacSph pan-stained tissue is found in **Supplementary Fig. 32**.

## Discussion

The data we present in this paper demonstrates that pan-ExM-t can clearly resolve molecular targets in the context of brain tissue ultrastructure. By combining ∼24-fold linear sample expansion with novel pan-stainings, we were able to (1) resolve and identify synapses and their substructures by their morphological characteristics, (2) localize protein markers to their subcellular compartments, and (3) delineate cellular membranes.

pan-ExM-t achieves this level of detail by taking advantage of an opportunity inaccessible to optical super-resolution techniques: the relative size of labels becomes irrelevant since labels appear shrunken by the linear expansion factor and the relative radius of a fluorescent dye (∼ 1 nm) approaches ∼1 nm/20 = ∼50 pm, a size comparable to that of an osmium atom used in heavy-metal EM stainings (∼200 pm). The distance between two proteins that originally were in close proximity (∼2 nm) grows to ∼40 nm, well beyond the Förster radius (∼5 nm) of typical fluorescent dyes. Bulk fluorescence staining of decrowded neuropil is therefore no longer limited by size, sampling and quenching restrictions of fluorescent dyes, and ultrastructural details, previously accessible only by EM, can now be examined on standard light microscopes.

We have shown that brain tissue structures such as neurons, glia, axons, dendrites, blood capillaries, pericytes, the endothelium, basal bodies, as well as subcellular features like DPs, PSDs, mitochondria cristae, ER tubules, myelin sheaths, multisynaptic boutons, spine necks, spine apparatuses, and the cartwheel structure of centrioles can all be identified in the pan-stain channel alone, analogous to classical EM techniques. We have also demonstrated, for the first time, that chemical synapses can be distinguished in a light microscopic image by their characteristic protein density patterns. These patterns could be characterized analogously to the Gray classification established for PTA and osmium stained synapses in EM (Gray, 1959; Klemann and Roubos, 2011; Harris and Weinberg, 2012). Taking advantage of pan-ExM-t’s strength of directly correlating synapse ultrastructure with immunolabeling against specific proteins, we observed that Class 1 and Class 2, but not Class 3, synapses are positive for Homer1 and PSD-95, supporting the functional Gray synapse concept that synapses with dense PSDs are likely excitatory. This example demonstrates our approach’s capability to map the molecular makeup of different synapse types within the context of their distinct ultrastructural properties, which could lead to a systematic exploration of synapse diversity and a determination of synapse sub-types that are vulnerable to diseases.

To achieve the highest ultrastructural preservation of tissue while maximizing sample expansion, we have shown that it is important to optimize fixation and gelation parameters. NHS ester pan-staining in this context serves as a readily accessible readout to assess the preservation and retention of proteins. We have demonstrated that reducing interprotein crosslinking enables 26-fold expansion without resorting to protease treatments or harsh denaturation conditions. We have also shown that hydrogels of high monomer concentrations (∼30% w/v) and sufficient crosslinker density (∼0.1% w/v) must be used to ensure that proteins are adequately sampled by the polymer matrix. In this context, it is not surprising that pan-ExM-t gels are stiffer than the gels prepared using the standard ExM protocol (Chen et al., 2015). Moreover, we found that ECS preservation is sensitive to the strength of fixatives used, with higher fixation resulting in lower ECS fractions. In all cases, the ECS is significantly more prominent in pan-ExM-t than in EM, emphasizing that the familiar appearance of tightly apposed neurites, typically seen in EM, is artifactual, and consistent with a previous live-STED microscopy characterization of brain ECS (Tonnesen et al., 2018).

In this work, we also explored the feasibility of imaging lipid membranes in 24-fold expanded brain tissue sections. We reasoned that attempts to preserve the endogenous lipid content and probe it after expansion are counterproductive. After all, since the free energy of lipid bilayers must be minimized, we hypothesize that even if we were to avoid detergent extraction, lipid membranes will eventually collapse into micelles following sample expansion. Our data shows that although lipophilic dyes in protein-denatured samples produce a distinctive pan-stain, these likely also stain the hydrophobic core of proteins. Our strategy was therefore to imprint lipid membranes onto the hydrogel before mechanical homogenization of the sample. We have shown here that pacSph, a photocrosslinkable and clickable sphingosine, enables neuronal membrane imaging in tissue fixed only with formaldehyde. We were able to delineate cellular organelles, synaptic anatomy, as well as membrane boundaries. Our new contrast complements our established protein pan-stain, and can potentially facilitate segmentation and neuronal tracing algorithms for connectomics analysis.

Tracing lipid boundaries in neuropil using EM has enabled reconstruction of the neural wiring diagrams of a 1,500-μm^3^ large volume in the mouse brain (Kasthuri et al., 2015), the *Drosophila* larval mushroom body (Eichler et al., 2017), and the *Caenorhabditis elegans* connectome (Cook et al., 2019). To date, however, no light microscopy method has been able to reconstruct a brain connectome, as these lack the ability of EM to reveal neurite ultrastructure without selective staining. Interestingly, one theoretical study supports the feasibility of reconstructing entire connectomes using lipid membrane labeling and *in-situ* molecular barcoding in 20-fold expanded tissue (Yoon et al., 2017). Based on our data, however, capitalizing on protein (instead of membrane) pan-staining in >20-fold expanded tissue might be a more feasible step towards this goal. Our data shows that cell boundaries can be detected easily by the absence of protein pan-staining. In contrast to lipids, which need to be negatively imprinted onto the expansion hydrogel first, proteins can be retained well through the expansion process. While we did not reconstruct neural connectivity in this work, individual neighboring neurites and pre- and post-synaptic compartments can be resolved relatively easily, laying the groundwork to computational tracing of neurons. We envision that combining pan-ExM-t with microscopes optimized for high-resolution and large field of view imaging with molecular optical barcoding techniques that are nucleic acid- (Askary et al., 2020) or protein-based (Wroblewska et al., 2018) will revolutionize this active area of research.

Similar to the early days of EM, we believe that many future optimizations will enhance this work. Sample preservation could be further improved, for example by using cryo-fixation (Laporte et al., 2022), lipid membrane delineation would benefit from systematic optimizations, and more experiments could be designed to characterize the multitude of ultrastructural features revealed by the presented pan-stainings. However, our core concept holds true regardless: that with adequate sample preservation and expansion, pan-staining brings light microscopy to the realm of ultrastructural context imaging.

No longer in the dark, we now see the brain in its complex makeup and the totality of its diverse neural processes: from the microstructure of cells and neuropil down to the nanoarchitecture of synaptic densities. No longer in the void, fluorescing antigens are now localized to their nanoscopic compartments, providing a functional dimension to the underlying structures they embody.

## Methods

### Key resources table

Please see **Supplementary Tables 1-5** for an overview of the reagents and materials used in this work.

### Resource availability

#### Lead contact

Further information and requests for resources and reagents should be directed to the lead contact, Joerg Bewersdorf.

#### Materials availability

This study did not generate any new materials.

### Experimental model and subject details

#### Neuron culture

Rat neuron cultures used to generate NHS ester, anti-Homer1, anti-Bassoon, and anti-PSD-95 images were cultured as described previously (Scheiffele and Biederer, 2007). In short, primary neuron cultures were prepared from hippocampal regions dissected from E18 Sprague Dawley Rat pups (Charles River Laboratories). The brains were removed, and hippocampi were isolated on ice in HBS medium (Sigma; catalog no. H2387) containing 10 mM HEPES (Gibco; catalog no. 15630-080). The hippocampal neurons were then dissociated via incubation at 37°C for 30 minutes in 2% papain solution (Worthington; catalog no. LS003127). Following incubation, hippocampi were washed and triturated in Neurobasal medium (Gibco; catalog no. 21103-049) supplemented with 5% fetal bovine serum (Atlanta; S11150), 2% B27 (Invitrogen; catalog no. 17504-044), 1% Glutamax (Invitrogen; catalog no. 35050-061) and gentamycin 1ng/ul (Gibco; catalog no. 1570060). The cells were plated at a density of 20,000 cells per Poly-DL-Ornithine (Sigma; catalog no. P0421-100MG) and Laminin (Corning; catalog no. #354232) coated 12-mm glass coverslip (Assistant; catalog no. 633009). To reduce glia growth, cultures were treated with 10 µM 5-Fluoro-2’-deoxyuridine (Sigma; catalog no. F0503). After 14 days in vitro, neurons were fed with Neurobasal medium (Gibco; catalog no. 21103-049) supplemented with 2% B27 (Invitrogen; catalog no. 17504-044), 1% Glutamax (Invitrogen; catalog no. 35050-061), 10 µM 5-Fluoro-2’-deoxyuridine (Sigma; catalog no. F0503) and gentamycin 1ng/ul (Gibco; catalog no. 1570060). Experiments were performed on 15–21 DIV cultures.

Rat neuron cultures used to generate NHS ester and anti-Synaptophysin images were cultured as follows: hippocampal CA3-CA1 regions were isolated and dissected from E18 Sprague Dawley Rats (Charles River Laboratories) in the presence of chilled hibernate E media (BrainBits). Hippocampal neurons were then incubated at 37°C for 25 minutes in 0.25% trypsin (Corning; catalog no. 25–053 CI) and immediately washed with DMEM (Thermofisher; catalog no. 11965–118) containing 10% fetal bovine serum (Thermofisher; catalog no. 16000044). Following trypsinization, hippocampi were triturated and immediately plated in DMEM containing 10% fetal bovine serum on poly-D-lysine (Thermofisher; catalog no. ICN10269491) coated 18 mm glass coverslips (Warner instruments; catalog no. 64–0734 [CS-18R17]). After 1 hour of incubation at 37°C, DMEM was removed and replaced with Neurobasal-A (Thermofisher; catalog no. 10888–022) medium containing 2% B-27 (Thermofisher; catalog no. 17504001) and 2 mM Glutamax (Gibco; catalog no. 35050061). Experiments were performed 15-17 days after plating.

Mouse hippocampal cultures used to generate NHS ester images were performed on C57BL/6 E17-E18 embryos, in accordance with IACUC protocol number 2019-07912. Pregnant animals were sacrificed with CO2 asphyxiation. The embryos were isolated from uterus horns through Cesarean section, followed by decapitation and hippocampal dissection. Hippocampi were dissociated using papain dissociation kit (Worthington LK003176) for 10 minutes at 30°C. Neurons were triturated and plated onto 12 mm Poly-D-lysine coated coverslips in a 24 well dish at a density of 150,000 cells per well in Neurobasal Plus media (Thermo-Fisher; catalog no. A3653401) supplemented with 2% B27 plus (Thermo-Fisher; catalog no. A3582801), and 50ug/ml of gentamicin (Thermo-Fisher; catalog no. 15750060). Fixation was performed at DIV 14-15.

Mouse hippocampal cultures used to generate 5-fold expansion images were performed on BalbC P0 pups, in accordance with IACUC protocol number 2019-07912. Hypothermia was induced in pups using ice, followed by decapitation and hippocampal dissection. Hippocampi were dissociated using 200 units Papain (Worthington LS003124, 200 units) for 40 minutes at 37°C. Neurons were triturated and plated onto 12 mm Poly-D-lysine coated coverslips in a 24 well dish at a density of 150,000 cells per well in Neurobasal/B27 Plus media (Thermo-Fisher; catalog no. A3653401) supplemented with 10% FBS and 1% Penicillin/Streptomycin (Thermo-Fisher; catalog no. 15140163). Following 3 hours of incubation at 37°C and 5% CO2, the media was replaced with Neurobasal/B27 Plus media (Thermo-Fisher; catalog no. A3653401) supplemented with 1% Penicillin/Streptomycin (Thermo-Fisher; catalog no. 15140163)). Fixation was performed at DIV 14-15.

#### Experimental models

Brain tissue experiments were conducted in 2 wild-type (C57BL/6) adult male mice purchased from the Charles River Laboratories and 2 wild-type (C57BL/6) adult male mice and 1 Thy1-EGFP mouse (strain Tg(Thy1-EGFP)MJrs/J, 4-8 weeks), purchased from the Jackson Laboratory.

### Method details

#### Neuron fixation

Neurons immunolabeled with Homer1, Bassoon, or PDS-95 were fixed with 4% formaldehyde in 1× PBS (Thermofisher; catalog no. 10010023) for 15 min at RT. Neurons immunolabeled with Synaptophysin were fixed with 3% FA and 0.1% glutaraldehyde in 1× PBS for 15 min at RT. Images showing nucleoids in mitochondria were from samples fixed with 3% FA + 0.1% GA in 1× PBS for 15 min at RT. Images showing centrioles and NPCs were from samples fixed with 4% FA in 1× PBS for 15 min at RT. After fixation, all samples were rinsed three times with 1× PBS and processed according to the pan-ExM protocol immediately after. Formaldehyde (FA; catalog no. 15710) and glutaraldehyde (GA; catalog no. 16019) were purchased from Electron Microscopy Sciences.

### Mice perfusion and brain fixation

All experiments were carried out in accordance with National Institutes of Health (NIH) guidelines and approved by the Yale IACUC.

All wild-type (C57BL/6) adult male mice were between P21 and P35 and maintained in the vivarium with a 12-h light–dark cycle, stable temperature at 22 ± 1 °C and humidity between 20 and 50%. The mice were anesthetized by a mixture of Ketamine (ketaset, Zoetis Inc, Cat # 010177) and Xylazine (Anased Injection, Akorn Animal health, NDC: 59399-110-20) with a 4:1 ratio. This mixture was then diluted 5x in saline, and administered IP. Mice were transcardially perfused first with ice-cold 1× PBS and then with either 4% FA, 4% FA + 0.1% FA, or 4% FA + 20% acrylamide (AAm; Sigma; catalog no. A9099) (all in 1× PBS, pH 7.4). Brains were isolated and post-fixed overnight in the same perfusion solution at 4 °C. Brains were subsequently washed and stored in PBS at 4 °C. Brains were mounted in ice-cold PBS and coronally sectioned at 70 µm using a vibrating microtome (Vibratome 1500, Harvard Apparatus). Medial hippocampal slices were selected and washed four times for 15 min in PBS at room temperature. Sections were stored in PBS at 4 °C for up to 3 months.

Thy1-EGFP mouse was anesthetized with ketamine (100 mg/kg, Butler Schein: 9952949) and xylazine (10 mg/kg, Akorn: NDC 59399-110-20) in saline (Hudson RCI: 200-59) soon after delivery. The mouse was transcardially perfused first with ice cold 1× PBS (Fisher; BP2944-100) and then with 4% FA (J.T. Baker; S898-07) + 20% AAm in 1× PBS, pH 7.4. Brain was isolated and postfixed overnight in 4% FA + 20% AAm in 1× PBS at 4°C. Brain was then washed in PBS and coronally sectioned at 70 μm using vibrating microtome (Leica VT1000, Leica Biosystems, Nussloch, Germany). Sections were stored in 1× PBS with 0.01% sodium azide (Sigma-Aldrich, St. Louis, MO) at 4°C for up to 3 months.

### pan-ExM-t reagents

Acrylamide (AAm; catalog no. A9099), N,N′-methylenebis(acrylamide) (BIS; catalog no. 146072) were purchased from Sigma-Aldrich. N,N’-(1,2-Dihydroxyethylene)bisacrylamide (DHEBA) was purchased from Sigma-Aldrich (catalog no. 294381) and Santa Cruz Biotechnologies (catalog no. sc-215503), with the latter being of better purity. Sodium acrylate (SA) was purchased from both Sigma Aldrich (catalog no. 408220) and Santa Cruz Biotechnologies (catalog no. sc-236893C). To verify that SA was of acceptable purity, 38% (w/v) solutions were made in water and checked for quality as previously reported (Asano et al., 2018). Only solutions that were light yellow were used. Solutions that were yellow and/or had a significant precipitate were discarded. Solutions with a minimum precipitate were centrifuged at 4000 rpm for 5 min and the supernatant was transferred to a new bottle and stored at 4 °C until use. Ammonium persulfate (APS) was purchased from both American Bio (catalog no. AB00112) and Sigma-Aldrich (catalog no. A3678). N,N,N′,N′-tetramethylethylenediamine (TEMED; catalog no. AB02020) was purchased from American Bio. 10X phosphate buffered saline (10X PBS; catalog no. 70011044) was purchased from Thermofisher. Sodium Dodecyl Sulfate (SDS; catalog no. AB01922) was purchased from American Bio and Guanidine hydrochloride (G-HCl; catalog no. G3272) was purchased from Sigma-Aldrich

### pan-ExM gelation chamber for dissociated neurons

The gelation chamber was constructed using a glass microscope slide (Sigma-Aldrich, catalog no. S8400) and two spacers, each consisting of a stack of two no. 1.5 22 × 22 mm coverslips (Fisher Scientific, catalog no. 12-541B). The spacers were superglued to the microscope slide on both sides of the neuron-adhered coverslip, with this latter coverslip glued in between. A no. 1.5 22 × 22 mm coverslip was used as a lid after addition of the gel solution. This geometry yielded an initial gel thickness size of ∼170 µm.

### pan-ExM-t gelation chamber for brain tissue sections

The gelation chamber was constructed using a glass microscope slide (Sigma-Aldrich, catalog no. S8400) and two spacers, each consisting of one no. 1.5 22 × 22 mm coverslip (Fisher Scientific, catalog no. 12-541B). The spacers were superglued to the microscope slide spaced ∼ 1 cm from one another. After incubation in activated first expansion gel solution (as described below in **First round of expansion for brain tissue sections**), the 70 µm-thick brain tissue section was placed and flattened on an additional no. 1.5 22 × 22 mm coverslip which was used as a chamber lid after addition of more activated first expansion gel solution on the microscope slide in-between the spacers. This geometry yielded an initial gel thickness size of ∼170 µm.

### First round of expansion for dissociated neurons

Neurons, previously fixed as described in the **Neuron fixation** section, were incubated in post-fixation solution (0.7% FA + 1% AAm (w/v) in 1× PBS) for 6–7 h at 37 °C. Next, the neurons were washed twice with 1× PBS for 10 min each on a rocking platform and embedded in the first expansion gel solution (18.5% (w/v) SA + 10% AAm (w/v) + 0.1% (w/v) DHEBA + 0.25% (v/v) TEMED + 0.25% (w/v) APS in 1× PBS). Gelation proceeded for 10-15 min at room temperature (RT) and then for 1.5 h at 37 °C in a humidified chamber. Coverslips with hydrogels were then incubated in ∼4 mL denaturation buffer (200 mM SDS + 200 mM NaCl + 50 mM Tris in MilliQ water, pH 6.8) in 6-well plates for 45 min at 37 °C. Gels were then transferred to denaturation buffer-filled 1.5 mL Eppendorf tubes and incubated at 73 °C for 1 h. Next, the gels were washed twice with PBS for 20 min each and stored in PBS overnight at 4 °C. Gels were then cut and placed in 6-well plates filled with MilliQ water for the first expansion. Water was exchanged twice every 30 min and once for 1 h. Gels expanded between 4.0× and 4.5× according to SA purity (see **pan-ExM-t reagents**).

### First round of expansion for brain tissue sections

Brain tissue, previously fixed and sectioned to 70 µm as described in the **Brain perfusion** section, were first incubated in inactivated first expansion gel solution (19% (w/v) SA + 10% AAm (w/v) + 0.1% (w/v) DHEBA in 1× PBS) for 30-45 min on ice and then in activated first expansion gel solution (19% (w/v) SA + 10% AAm (w/v) + 0.1% (w/v) DHEBA + 0.075% (v/v) TEMED + 0.075% (w/v) APS in 1× PBS) for 15-20 min on ice before placing in gelation chamber. The tissue sections were gelled for 15 min at RT and 2 h at 37 °C in a humidified chamber. Next, the tissue-gel hybrids were peeled off of the gelation chamber and incubated in ∼4 mL denaturation buffer (200 mM SDS + 200 mM NaCl + 50 mM Tris in MilliQ water, pH 6.8) in 6-well plates for 15 min at 37 °C. Gels were then transferred to denaturation buffer-filled 1.5 mL Eppendorf tubes and incubated at 75 °C for 4 h. The gels were washed twice with 1× PBS for 20 min each and once overnight at RT. Gels were optionally stored in 1× PBS at 4 °C. The samples were then placed in 6-well plates filled with MilliQ water for the first expansion. Water was exchanged three times every 1 h. Gels expanded between ∼5× according to SA purity (see **pan-ExM-t reagents**).

### Re-embedding in neutral hydrogel

Expanded hydrogels (of both dissociated neuron and tissue samples) were incubated in fresh re-embedding neutral gel solution (10% (w/v) AAm + 0.05% (w/v) DHEBA + 0.05% (v/v) TEMED + 0.05% (w/v) APS) two times for 20 min each on a rocking platform at RT. Immediately after, the residual gel solution was removed by gentle pressing with Kimwipes. Each gel was then sandwiched between one no. 1.5 coverslip and one glass microscope slide. Gels were incubated for 1.5 h at 37 °C in a nitrogen-filled and humidified chamber. Next, the gels were detached from the coverslips and washed two times with 1× PBS for 20 min each on a rocking platform at RT. The samples were stored in 1× PBS at 4 °C. No additional post-fixation of samples after the re-embedding step was performed.

### Second round of expansion

Re-embedded hydrogels (of both dissociated neuron and tissue samples) were incubated in fresh second hydrogel gel solution (19% (w/v) SA + 10% AAm (w/v) + 0.1% (w/v) BIS + 0.05% (v/v) TEMED + 0.05% (w/v) APS in 1× PBS) two times for 15 min each on a rocking platform in 4 °C. Each gel was then sandwiched between one no. 1.5 coverslip and one glass microscope slide. Gels were incubated for 1.5 h at 37 °C in a nitrogen-filled and humidified chamber. Next, to dissolve DHEBA, gels were detached from the coverslips and incubated in 200 mM NaOH for 1 h on a rocking platform at RT. Gels were afterwards washed three to four times with 1× PBS for 30 min each on a rocking platform at RT or until the solution pH reached 7.4. The gels were optionally stored in 1× PBS at 4 °C. Subsequently, the gels were labeled with antibodies and pan-stained with NHS ester dyes. Finally, the gels were placed in 6-well plates filled with MilliQ water for the second expansion. Water was exchanged at least three times every 1 h at RT. Gels expanded ∼ 4.5× according to SA purity (see **pan-ExM-t Reagents**) for a final expansion factor of ∼16× (dissociated neurons) and ∼24× (brain tissue).

### Antibody labeling of neurons post-expansion

Samples immunolabeled with Homer1, Bassoon, and PSD-95 were fixed with 4% FA in 1× PBS for 15 min as described in the **Neuron fixation** section, processed with pan-ExM, and incubated overnight (∼15 h) with monoclonal anti-Homer1 antibody (abcam, catalog no. ab184955), monoclonal anti-Bassoon antibody (abcam, catalog no. ab82958), or monoclonal anti-PSD-95 antibody (antibodiescinc, catalog no. 75-028) diluted 1:500 in antibody dilution buffer (0.05% TX-100 + 0.05% NP-40 + 0.2% BSA in 1× PBS). All primary antibody incubations were performed on a rocking platform at 4 °C. Gels were then washed with 0.1% (v/v) TX-100 in 1× PBS twice for 20 min each on a rocking platform at RT and once overnight (∼15 h) at 4 °C. Next, samples were incubated overnight with ATTO594-conjugated anti-mouse antibodies (Sigma-Aldrich, catalog no. 76085) or ATTO594-conjugated anti-rabbit antibodies (Sigma-Aldrich, catalog no. 77671) diluted 1:500 in antibody dilution buffer. All secondary antibody incubations were performed on a rocking platform at 4 °C. The gels were subsequently washed in 0.1% (v/v) TX-100 in 1× PBS twice for 20 min each at RT and once overnight at 4 °C. Gels were stored in PBS at 4 °C until subsequent treatments. Bovine serum albumin (BSA; catalog no. 001-000-162) was purchased from Jackson ImmunoResearch and NP-40 (catalog no. T8786) and Triton X-100 (TX-100; catalog no. T8787) was purchased from Sigma-Aldrich.

Samples immunolabeled with both Homer1 and Bassoon simultaneously were fixed with 4% FA in 1× PBS for 15 min as described in the **Neuron fixation** section, processed with pan-ExM, and incubated overnight (∼15 h) with both monoclonal anti-Homer1 antibody (abcam, catalog no. ab184955) and monoclonal anti-Bassoon antibody (abcam, catalog no. ab82958) diluted 1:500 in antibody dilution buffer (0.05% TX-100 + 0.05% NP-40 + 0.2% BSA in 1× PBS). Primary antibody incubations were performed on a rocking platform at 4 °C. Gels were then washed with 0.1% (v/v) TX-100 in 1× PBS twice for 20 min each on a rocking platform at RT and once overnight (∼15 h) at 4 °C. Next, samples were incubated overnight (∼15 h) with both ATTO594-conjugated anti-rabbit antibodies (Sigma-Aldrich, catalog no. 77671) and ATTO647N-conjugated anti-mouse antibodies (Sigma-Aldrich, catalog no. 50185) diluted 1:500 in antibody dilution buffer. Secondary antibody incubations were performed on a rocking platform at 4 °C. The gels were subsequently washed in 0.1% (v/v) TX-100 in 1× PBS twice for 20 min each at RT and once overnight at 4 °C. Gels were stored in PBS at 4 °C until subsequent treatments.

Samples immunolabeled with Synaptophysin were fixed with 3% FA + 0.1% GA 1× PBS as described in the **Neuron fixation** section, processed with pan-ExM, and incubated for ∼36-40 h at 37° C with rabbit anti-synaptophysin (Cell signaling technology, catalog no. 36406) diluted 1:250 in antibody dilution buffer (2% (w/v) BSA in 1× PBS). Gels were then washed in PBS-T (0.1% (v/v) Tween 20 in 1× PBS) three times for 20 min each on a rocking platform at RT. Next, samples were incubated for ∼16-20 h at 37°C with donkey anti-rabbit CF568 (Biotium, catalog no. 20098) diluted 1:250 in antibody dilution buffer. The gels were washed in PBS-T three times for 20 min each at RT each before subsequent pan-staining.

### Antibody labeling of brain tissue samples post-expansion

Brain tissue samples immunolabeled with Homer1, Bassoon, PSD-95, and Synaptophysin were previously fixed with 4% FA + 20% AAm in 1× PBS overnight at 4 °C and sectioned to 70 µm as described in the **Brain perfusion** section. Next, they were washed three times with 1× PBS for 30 min each on a rocking platform at RT and processed with the pan-ExM-t protocol with this modification: the monomer composition of the second expansion hydrogel was 10% (w/v) AAm + 9% SA (w/v). Samples were subsequently incubated for ∼30 h with monoclonal anti-Homer1 antibody (abcam, catalog no. ab184955), monoclonal anti-Bassoon antibody (abcam, catalog no. ab82958), monoclonal anti-PSD-95 antibody (antibodiescinc, catalog no. 75-028), or anti-Synaptophysin antibody (SYSY, catalog no. 101 011) diluted 1:250 in antibody dilution buffer (0.05% TX-100 + 0.05% NP-40 + 0.2% BSA in 1× PBS). All primary antibody incubations were performed on a rocking platform at 4 °C. Next, samples were incubated for ∼30 h with ATTO594-conjugated anti-mouse antibodies (Sigma-Aldrich, catalog no. 76085) or ATTO594-conjugated anti-rabbit antibodies (Sigma-Aldrich, catalog no. 77671) diluted 1:250 in antibody dilution buffer. Gels were then washed in 0.1% (v/v) TX-100 in 1× PBS four times for 30 min to 1 h each on a rocking platform at RT and once overnight at 4 °C. All secondary antibody incubations were performed on a rocking platform at 4 °C. The gels were subsequently washed in 0.1% (v/v) TX-100 in 1× PBS four times for 30 min to 1 h each on a rocking platform at RT and once overnight at 4 °C. Gels were stored in PBS at 4 °C until subsequent treatments.

Brain tissue samples immunolabeled with GFAP, MBP, and GFP were previously fixed with 4% FA + 20% AAm in 1× PBS overnight at 4 °C and sectioned to 70 µm as described in the **Brain perfusion** section. Next, they were washed three times with 1× PBS for 30 min each on a rocking platform at RT and processed with the pan-ExM-t protocol. Samples were subsequently incubated for ∼30 h with polyclonal anti-GFAP antibody (Thermofisher, catalog no. PA1-10019), monoclonal anti-MBP (BioLegend, catalog no. 808401), or polyclonal anti-GFP (Thermofisher, catalog no. A-1122) diluted 1:500 in antibody dilution buffer (0.05% TX-100 + 0.05% NP-40 + 0.2% BSA in 1× PBS). All primary antibody incubations were performed on a rocking platform at 4 °C. Gels were then washed in 0.1% (v/v) TX-100 in 1× PBS four times for 30 min to 1 h each on a rocking platform at RT and once overnight at 4 °C. Next, samples were incubated for ∼30 h with ATTO594-conjugated anti-mouse antibodies (Sigma-Aldrich, catalog no. 76085) or ATTO594-conjugated anti-rabbit antibodies (Sigma-Aldrich, catalog no. 77671) diluted 1:500 in antibody dilution buffer. Gels were then washed in 0.1% (v/v) TX-100 in 1× PBS four times for 30 min to 1 h each on a rocking platform at RT and once overnight at 4 °C. All secondary antibody incubations were performed on a rocking platform at 4 °C. The gels were subsequently washed in 0.1% (v/v) TX-100 in 1× PBS four times for 30 min to 1 h each on a rocking platform at RT and once overnight at 4 °C. Gels were stored in PBS at 4 °C until subsequent treatments.

### NHS ester pan-staining of neurons

After antibody labeling, gels were incubated for 1.5 h with 20 µg/mL NHS ester-ATTO532 (Sigma-Aldrich, catalog no. 88793) (Homer1, Bassoon, PSD-95, centriole, and NPC samples), or 20 µg/mL NHS ester-ATTO488 (Sigma-Aldrich, catalog no. 41698) (Synaptophysin samples), or 20 µg/mL NHS ester-ATTO594 (Sigma-Aldrich, catalog no. 08741) (mitochondria samples), dissolved in 100 mM sodium bicarbonate solution (Sigma-Aldrich, catalog no. SLBX3650) on a rocking platform at RT. The gels were subsequently washed four to six times in either 1× PBS or PBS-T for 20 min each on a rocking platform at RT.

### NHS ester pan-staining of brain tissue sections

Gels were incubated for 2 h with either 30 µg/mL NHS ester-ATTO594 (Sigma-Aldrich, catalog no. 08741) (ultrastructural samples in **Fig. 3**) or 30 µg/mL NHS ester-ATTO532 (Sigma-Aldrich, catalog no. 88793) (all immunolabeled and lipid-stained samples), dissolved in 100 mM sodium bicarbonate solution on a rocking platform at RT. The gels were subsequently washed four to six times in either 1× PBS or PBS-T for 20 min each on a rocking platform at RT.

### Palmitate pan-staining of neurons

Live and dissociated 80%-confluent hippocampal neurons were incubated with 50 µM azide-functionalized palmitate (Thermofisher, catalog no. C10265) diluted in delipidated medium (DMEM + 10% charcoal-stripped FBS; Thermofisher, catalog no. A3382101) for 5 h at 37 °C and 5% CO2. Next, the neurons were fixed with 3% FA + 0.1% GA in 1× PBS for 15 min at RT and processed according to the pan-ExM protocol. Prior to NHS ester pan-staining with ATTO532, CuAAC (Copper(I)-catalyzed Azide-Alkyne Cycloaddition) was performed using the Click-iT Cell Reaction Buffer Kit (Thermo Fisher, catalog no. C10269) according to manufacturer instructions. Alkyne-functionalized ATTO590 dye (Sigma-Aldrich, catalog no. 93990) was used at a concentration of 5 µM. After CuAAC, the gels were washed twice with 2% (w/v) delipidated BSA (Sigma Aldrich, catalog no. A4612) in 1× PBS for 20 min each on a rocking platform at RT and twice with 1× PBS for 20 min each at RT.

### SYTOX Green staining post-expansion

pan-ExM and pan-ExM-t processed gels were incubated with SYTOX Green (Invitrogen, catalog no. S7020) diluted 1:5,000 (dissociated neuron samples) and 1:3,000 (brain tissue samples) in 1× PBS for 30 min on a rocking platform at RT. The gels were then washed three times with PBS-T for 20 min each on a rocking platform at RT.

### pan-ExM and pan-ExM-t sample mounting

After expansion, the gels were mounted on glass-bottom dishes (35 mm; no. 1.5; MatTek). A clean 18-millimeter diameter coverslip (Marienfeld, catalog no. 0117580) was put on top of the gels after draining excess water using Kimwipes. The samples were then sealed with two-component silicone glue (Picodent Twinsil, Picodent, Wipperfürth, Germany). After the silicone mold hardened (typically 15–20 min), the samples were stored in the dark at 4 °C until they were imaged.

### Image acquisition

All confocal images were acquired using a Leica SP8 STED 3X equipped with a SuperK Extreme EXW-12 (NKT Photonics) pulsed white light laser as an excitation source. The STED imaging mode of the instrument was not used in this work. Images were acquired using either an HC FLUOTAR L 25×/0.95 NA water objective, APO 63×/1.2 NA water objective, or an HC PL APO 86×/1.2 NA water CS2 objective. Application Suite X software (LAS X; Leica Microsystems) was used to control imaging parameters. ATTO532 was imaged with 532-nm excitation. ATTO594 was imaged with 585-nm excitation. CF568 was imaged with 568-nm excitation. ATTO647N was imaged with 647-nm excitation. ATTO488 and SYTOX Green were imaged with 488-nm excitation.

### Image processing

Images were visualized, smoothed, gamma-, and contrast-adjusted using FIJI/ImageJ software. STED and confocal images were smoothed for display with a 0.4 to 1.5-pixel sigma Gaussian blur. Minimum and maximum brightness were adjusted linearly for optimal contrast.

All line profiles were extracted from the images using the Plot Profile tool in FIJI/ImageJ. Stitching of tiled images was performed using the Pairwise Stitching tool in FIJI/ImageJ (Preibisch et al., 2009).

### 3D data visualization

3D data stacks were processed with the microscopy image analysis software Imaris (Oxford Instruments; Imarix x64 versions 9.6 or 9.8). Data sets with low signal-to-noise ratios were smoothed with Gaussian filters. Gamma correction and displayed color range were adjusted for contrast to highlight features of interest. Generated videos were further processed with Adobe Premiere Pro (version 22.2.0) to add headers and link videos .

### Phosphotungstic acid (PTA) staining and electron microscopy

Brain tissue was fixed in 2.5% GA in phosphate buffer (pH 7.4) for one hour at room temperature, it was then cut into 100-µm thick slices by a vibratome and dehydrated in a graded series of ethanol to 95%. The tissue sections were incubated in 1% phosphotungstic acid (Sigma, catalog no. P6395) in 0.5% ethanol for 2 hours at room temperature. After rinsing with propylene oxide (Electron Microscopy Sciences (EMS), catalog no. 75569), the sections embedded in EMbed 812 resin (EMS, catalog no. 14120), polymerized in a 60 °C oven overnight. Thin sections (65 nm) were cut by a Leica ultramicrotome (UC7) and directly collected on the sample grids without further metal staining. EM images were acquired by a FEI Tecnai transmission electron microscope at an accelerating voltage of 80 kV, an Olympus Morada CCD camera and iTEM imaging software.

### Neuron expansion factor calculation

Images of hippocampal mouse and rat neuron cell nuclei in non-expanded and pan-ExM expanded samples stained with SYTOX Green (1:5,000) were acquired with a Leica SP8 STED 3X microscope using a HCX PL Fluotar 10×/0.30 dry objective. Average nuclear cross-sectional areas were determined using FIJI/ImageJ software. To calculate the expansion factor, the average nuclear cross-sectional area in pan-ExM samples was divided by the average nuclear cross-sectional area of non-expanded samples. The square root of this ratio represents an estimate of the linear expansion factor. For nuclei cross-section measurements in pan-ExM samples, 38 nuclei were analyzed from 4 independent experiments. For nuclei cross-section measurements in non-expanded samples, 279 nuclei were analyzed from 4 independent experiments.

Using 2-pixel thick line profiles in FIJI, the peak-to-peak distances between the intensity distributions of the dense projection (DP) and postsynaptic density (PSD) NHS ester signals in the same samples were measured and divided by the nuclear expansion factor determined above. For this measurement, 44 line profiles were drawn from 4 independent experiments. The DP-PSD value was determined to be 81.9 nm and was used to convert DP-PSD measurements to expansion factors in all subsequent experiments. Results are summarized in **Fig. 2n**.

### General neuron and brain tissue expansion factor calculation

Using 2-pixel thick line profiles in FIJI, the peak-to-peak distances between the intensity distributions of the DP and PSD NHS ester signals were measured and divided by 81.9 nm, the DP-PSD length determined in **Neuron expansion factor calculation**.

### Distance between individual DPs calculation

Using 2-pixel thick line profiles in FIJI, the peak-to-peak distances between the intensity distributions of individual dense projections (DPs) were measured and divided by the determined expansion factor. For this measurement, 78 line profiles were drawn from 41 synapses in 6 independent experiments. Results are summarized in **Fig. 2o**.

### Measurements of synaptic protein distributions in neurons

Using 2-pixel thick line profiles in FIJI, the peak-to-peak distances between the intensity distributions between Homer1 and PSD, PSD-95 and DP, and Bassoon and PSD were measured and divided by the determined experiment expansion factor. For Bassoon-PSD measurements in neurons, 50 line profiles were drawn from 3 independent samples. For Homer1-DP measurements in neurons, 85 line profiles were drawn from 4 independent samples. For PSD95-DP measurements in neurons, 25 line profiles were drawn from 2 independent samples. Results are summarized in **Fig. 2p-s**.

### Measurement of anti-Homer1 signal across dilution factors

Neuron samples were labeled with anti-Homer1 primary antibody and ATTO594-conjugated anti-rabbit antibodies as described in **Antibody labeling of neurons post-expansion** with this modification: Samples were labeled with primary and secondary antibodies both diluted 1:250 or 1:500 or 1:1000.

Relative protein retention was measured by comparing the peak intensity of Homer1 ATTO504 signal from 2-pixel thick line profiles in FIJI drawn across the punctate signal. For *1:250* measurements, 45 peak intensities were recorded from 7 FOVs. For *1:500* measurements, 13 peak intensities were recorded from 3 FOVs. For *1:1000* measurements, 6 peak intensity measurements were recorded from 1 FOV. Results are summarized in **Supplementary Fig. 7j**.

### Assessment of fixation effects in brain tissue ultrastructural preservation

*Fix-1* brain tissue samples were previously fixed with 4% FA in 1× PBS overnight at 4 °C and sectioned to 70 µm as described in the **Brain perfusion** section. Next, they were washed three times with 1× PBS for 30 min each on a rocking platform at RT and processed with the pan-ExM-t protocol. *Fix-2* brain tissue samples were previously fixed with 4% FA in 1× PBS overnight at 4 °C and sectioned to 70 µm as described in the **Brain perfusion** section. Next, they were post-fixed in 0.7% FA + 1% AAm in 1× PBS for 7 h at 37 °C, washed three times with 1× PBS for 30 min each on a rocking platform at RT, and processed with the pan-ExM-t protocol. *Fix-3* brain tissue samples were previously fixed with 4% FA in 1× PBS overnight at 4 °C and sectioned to 70 µm as described in the **Brain perfusion** section. Next, they were post-fixed in 4% FA + 20% AAm in 1× PBS for 7 h at 37 °C, washed three times with 1× PBS for 30 min each on a rocking platform at RT, and processed with the pan-ExM-t protocol. *Fix-4* brain tissue samples were previously fixed with 4% FA + 0.1% GA in 1× PBS overnight at 4 °C and sectioned to 70 µm as described in the **Brain perfusion** section. Next, they were post-fixed in 4% FA + 20% AAm in 1× PBS for 12 h at 4°C, washed three times with 1× PBS for 30 min each on a rocking platform at RT, and processed with the pan-ExM-t protocol. *Fix-5* brain tissue samples were previously fixed with 4% FA in 1× PBS overnight at 4 °C and sectioned to 70 µm as described in the **Brain perfusion** section. Next, they were treated with 0.1 mg/mL acryloyl-X, SE (AcX; Thermofisher, catalog no. A20770) in 1× PBS for 3 h at RT, washed three times with 1× PBS for 30 min each on a rocking platform at RT, and processed with the pan-ExM-t protocol. *Fix-6* brain tissue was previously fixed with 4% FA + 20% AAm in 1× PBS overnight at 4 °C and sectioned to 70 µm as described in the **Brain perfusion** section. Next, they were washed three times with 1× PBS for 30 min each on a rocking platform at 4°C and processed with the pan-ExM-t protocol.

Expansion factors were calculated as described in the **General neuron and brain tissue expansion factor calculation** section. For *Fix-2* DP-PSD measurements, 38 line profiles were drawn from 6 FOVs. For *Fix-3* DP-PSD measurements, 81 line profiles were drawn from 9 FOVs. For *Fix-4* DP-PSD measurements, 77 line profiles were drawn from 8 FOVs. For *Fix-5* DP-PSD measurements, 74 line profiles were drawn from 9 FOVs. For *Fix-6* DP-PSD measurements, 254 line profiles were drawn from 10 FOVs in 3 independent experiments. Results are summarized in **Supplementary Fig. 12**.

To calculate the ECS + lipid membrane (ECS+) fraction, ∼250 x 250 µm^2^ large-FOV images that show predominantly neuropil pan-stained with NHS ester, were processed with a Gaussian blur of 1 sigma (*Fix-1*, *Fix-2* and *Fix-5*) or 0.5 sigma (*Fix-3*, *Fix-4* and *Fix-6*) in Fiji. A mask created with manual thresholding was used to exclude large gaps in the image that do not represent neurites. The images were then thresholded using the Otsu method (Otsu, 1979) with 100%, 70%, 50%, and 30% of the automatically determined threshold. The ECS+ fraction was calculated by dividing the total thresholded pixels by the total area represented by the masked pixels. For *Fix-2*, 8 FOVs from 1 independent experiment were processed. For *Fix-3*, 8 FOVs from 1 independent experiment were processed. For *Fix-4*, 8 FOVs from 1 independent experiment were processed. For *Fix-5*, 8 FOVs from 1 independent experiment were processed. For *Fix-6*, 14 FOVs from 3 independent experiments were processed. Results are summarized in **Supplementary Fig. 13**.

### Assessment of denaturation effects in brain tissue expansion factor

*Denat-4, Denat-6,* and *Denat-8* brain tissue samples were previously fixed with 4% FA + 20% AAm in 1× PBS overnight at 4 °C and sectioned to 70 µm as described in the **Brain perfusion** section. Next, they were washed three times with 1× PBS for 30 min each on a rocking platform at RT and processed with the pan-ExM-t protocol with these modifications: *Denat-6* samples were denatured for 6 h and *Denat-8* samples were denatured for 8 h.

Expansion factors were calculated as described in the **General neuron and brain tissue expansion factor calculation section**. For *Denat-4* DP-PSD measurements, 50 line profiles were drawn from 3 FOVs. For *Denat-6* DP-PSD measurements, 42 line profiles were drawn from 3 FOVs. For *Denat-8* DP-PSD measurements, 100 line profiles were drawn from 3 FOVs. Results are summarized in **Supplementary Fig. 14**.

Relative protein retention was measured by comparing the peak intensity of DP NHS ester signal from 2-pixel thick line profiles in FIJI. For *Denat-4* measurements, 50 DP peak intensities were recorded from 3 FOVs. For *Denat-6* measurements, 42 DP peak intensities were recorded from 3 FOVs. For *Denat-8* measurements, 48 DP peak intensity measurements were recorded from 3 FOVs. The intensity values were multiplied by the cube of the expansion factor determined for every denaturation condition. Results are summarized in **Supplementary Fig. 14**.

### Assessment of SA monomer concentration effects in brain tissue preservation

*19SA/19SA, 19SA/9SA, 9SA/19SA,* and *9SA/9SA* brain tissue samples were previously fixed with 4% FA + 20% AAm in 1× PBS overnight at 4 °C and sectioned to 70 µm as described in the **Brain perfusion** section. Next, they were washed three times with 1× PBS for 30 min each on a rocking platform at RT and processed with the pan-ExM-t protocol with these modifications: the monomer composition of the second expansion hydrogel in *19SA/9SA* samples was 10% (w/v) AAm + 9% SA (w/v); the monomer composition of the first expansion hydrogel in *9SA/19SA* samples was 10% (w/v) AAm + 9% SA (w/v); and the monomer composition of the first and second expansion hydrogels in *9SA/9SA* samples was 10% (w/v) AAm + 9% SA (w/v).

Expansion factors were calculated as described in the **General neuron and brain tissue expansion factor calculation section**. For *19SA/19SA* DP-PSD measurements, 254 line profiles were drawn from 10 FOVs in 3 independent experiments. For *19SA/9SA* DP-PSD measurements, 267 line profiles were drawn from 12 FOVs in 2 independent experiments. For *9SA/19SA* DP-PSD measurements, 88 line profiles were drawn from 6 FOVs in 2 independent experiments. For *9SA/9SA* DP-PSD measurements, 86 line profiles were drawn from 6 FOVs in 2 independent experiments. All results are summarized in **Supplementary Fig. 20e**.

### Measurements of synaptic protein distributions in brain tissue

Using 2-pixel thick line profiles in FIJI, the peak-to-peak distances between the intensity distributions between Homer1 and PSD, PSD-95 and DP, and Bassoon and PSD were measured and divided by the determined experiment expansion factor. For Homer1-DP measurements in brain tissue, 120 line profiles were drawn from 5 FOVs in 2 independent experiments. For PSD-95-DP measurements in brain tissue, 113 line profiles were drawn from 3 FOVs in one independent experiment. For Bassoon-PSD measurements in brain tissue, 74 line profiles were drawn from 3 FOVs in one independent experiment. Results are summarized in **Fig. 4ooo-rrr**.

### Measurement of BDP signal in 4-fold expanded brain tissue across fixation conditions

To compare the effect of fixation on lipid retention, brain tissue was fixed with either 4% FA in 1× PBS (*FA-fix*) or 4% FA+ 0.1% GA in 1× PBS (*FA/GA-fix*) overnight at 4 °C and sectioned to 70 µm as described in the **Brain perfusion** section. Several *FA/GA-fix* tissue sections were post-fixed with 0.01% osmium tetroxide (4% osmium solution; EMS, catalog no. 19150) in 1× PBS for 1h at RT (*FA/GA/OsO4-fix*), washed three times with 1× PBS at RT, and stored in 4 °C until subsequent treatment. All tissue sections were post-fixed in 4% FA + 20% AAm in 1× PBS for 15 h at 4 °C, washed three times with 1× PBS for 30 min each on a rocking platform at 4°C, and processed only with the **First round of expansion for brain tissue** protocol with this modification: tissue sections were denatured in SDS buffer for 1 h. For NHS ester pan-staining, gels were incubated with 10 µg/mL NHS ester ATTO532 (Sigma-Aldrich, catalog no. 88793) in 1× PBS for 30 min on a rocking platform at RT and washed three times with 1× PBS for 30 min each. For BDP pan-staining, gels were incubated with 10 µM BODIPY TR Methyl Ester (Thermofisher, catalog no. C34556) in 1× PBS for 1 h on a rocking platform at RT and washed three times with 1× PBS for 30 min each. Gels were subsequently expanded in MilliQ water and imaged.

BDP pan-staining signal in axons was measured by recording the peak intensity of BDP signal from 10-pixel thick line profiles drawn across the width of axons in FIJI. For *FA-fix* measurements, 46 peak intensities were recorded from 5 FOVs. For *FA/GA-fix* measurements, 46 peak intensities were recorded from 5 FOVs. For *FA/GA/OsO4-fix* measurements, 38 peak intensities were recorded from 5 FOVs. Results are summarized in **Supplementary Fig. 24bb**.

### Measurement of BDP signal in 4-fold expanded brain tissue across denaturation conditions

To compare the effect of fixation on lipid retention, brain tissue was fixed with 4% FA+ 0.1% GA in 1× PBS overnight at 4 °C and sectioned to 70 µm as described in the **Brain perfusion** section. All tissue sections were post-fixed in 4% FA + 20% AAm in 1× PBS for 15 h at 4 °C, washed three times with 1× PBS for 30 min each on a rocking platform at 4°C, and processed only with the **First round of expansion for brain tissue** protocol with this modification: tissue sections were denatured with either SDS buffer at 75°C for 1 h (*SDS-1h*), or SDS buffer at 75°C for 4 h, (*SDS-4h*), non-denatured (*PBS*), or denatured with guanidine hydrochloride (*G-HCl*) buffer (6 M G-HCl + 5 mM DTT + 50 mM Tris in MilliQ water, pH 7.5) at 42°C overnight. For NHS ester pan-staining, gels were incubated with 10 µg/mL NHS ester ATTO532 (Sigma-Aldrich, catalog no. 88793) in 1× PBS for 30 min on a rocking platform at RT and washed three times with 1× PBS for 30 min each. For BDP pan-staining, gels were incubated with 10 µM BODIPY TR Methyl Ester (Thermofisher, catalog no. C34556) in 1× PBS for 1 h on a rocking platform at RT and washed three times with 1× PBS for 30 min each. Gels were subsequently expanded in MilliQ water and imaged.

BDP pan-staining signal in axons was measured by recording the peak intensity of BDP signal from 10-pixel thick line profiles drawn across the width of axons in FIJI. For *SDS-1h* measurements, 40 peak intensities were recorded from 4 FOVs. For *SDS-4h* measurements, 20 peak intensities were recorded from 1 FOV. For *PBS* measurements, 41 peak intensities were recorded from 2 FOVs. For *G-HCl* measurements, 24 peak intensities were recorded from 1 FOV. Results are summarized in **Supplementary Fig. 27k**.

BDP pan-staining signal in mitochondria was measured by recording the peak intensity of BDP signal from 10-pixel thick line profiles drawn across the width of mitochondria in FIJI. For *SDS-1h* measurements, 40 peak intensities were recorded from 4 FOVs. For *SDS-4h* measurements, 18 peak intensities were recorded from 1 FOV. For *PBS* measurements, 41 peak intensities were recorded from 2 FOVs. For *G-HCl* measurements, 20 peak intensities were recorded from 1 FOV. Results are summarized in **Supplementary Fig. 27l.**

### Measurement of pacSph pan-staining signal across UV irradiation conditions

To compare the effect of fixation on lipid retention, brain tissue was fixed with 4% FA+ 20% AAm in 1× PBS overnight at 4 °C and sectioned to 70 µm as described in the **Brain perfusion** section. Sections were incubated in 50 µM pacSph (PhotoClick Sphingosine; Avanti Polar Lipids, catalog no. 900600) in 1× PBS overnight at 4 °C, washed three times in 1× PBS for 1 h each at 4 °C, and either stored at 4 °C (*noUV*) or photo-crosslinked. Samples that were photo-crosslinked were kept on ice and irradiated with an 8 Watt, 365 nm UV light source (Analytik Jena, UVL-18) 1 cm away from a lamp surface for 30 min on both section surface sides, either before hydrogel embedding (*UV-before*) or immediately after **First round of expansion for brain tissue** while the tissue-hydrogel hybrid was still sandwiched between a glass microscope slide and glass coverslip (*UV-after*). Samples were processed with the remaining steps of the pan-ExM-t protocol with this modification: before NHS ester pan-staining, CuAAC was performed using the Click-iT Protein Reaction Buffer Kit (Thermo Fisher, catalog no. C10276) according to manufacturer instructions. Azide-functionalized ATTO590 dye (ATTO-TEC, catalog no. AD 590-101) was used at a concentration of 5 µM. After CuAAC, the gels were washed once with 2% (w/v) delipidated BSA (Sigma Aldrich, catalog no. A4612) in 1× PBS for 20 min each and three times with 1× PBS for 20 min each on a rocking platform at RT.

pacSph pan-staining signal in mitochondria was measured by recording the peak intensity of ATTO590 signal from 10-pixel thick line profiles drawn across the width of mitochondria in FIJI. For *noUV* measurements, 35 peak intensities were recorded from 5 FOVs in one independent experiment. For *UV-before* measurements, 37 peak intensities were recorded from 5 FOVs in one independent experiment. For *UV-after* measurements, 39 peak intensities were recorded from 5 FOVs in one independent experiment. Results are summarized in **Fig. 6m**.

### Quantification and statistical analysis

For all quantitative experiments, the number of samples and independent reproductions are listed in the figure legends. An unpaired two-tailed t-test in GraphPad Prism 9 was used to analyze the data.

## Data availability

The datasets generated and/or analyzed during the current study are available from the corresponding author on reasonable request.

## Acknowledgements

We want to thank Mark Lessard for assisting with vibratome sectioning. This work was supported by grants from the Wellcome Trust (203285/B/16/Z), NIH (P30 DK045735; S10 OD020142; R56MH122449 and MH115939 to A.J.K.; NS036251 to P.D.C) and a Yale Kavli Institute for Neuroscience Faculty Award (to J.B. and A.J.K.). H.F. was supported by a HHMI/LSRF postdoctoral fellowship. R.C.S. was supported by an NIH Individual Predoctoral MD/PhD Fellowship (MH124284).

## Author contributions

O.M. developed and optimized the sample preparation protocols. I.K., H.F., J.S., R.N., R.C.S. and S.L. cultured and fixed dissociated neurons. I.K., H.F., R.N. and K.W. provided fixed mouse brain samples. O.M. and R.K. expanded, labeled, and image neuron samples. O.M. expanded and imaged brain tissue samples. O.M. and P.K. labeled brain tissue samples. X.L. provided EM images. O.M. and J.B. visualized the data. O.M. analyzed and quantified all the data. All authors interpreted the data. O.M. and J.B. wrote the manuscript with input from all authors.

