## Supplementary Tables and Figures for "All-optical visualization of specific molecules in the ultrastructural context of brain tissue"

### SUPP TABLE. 1

**Supplementary Table 1:** Polymerization, buffer, and fixative reagents used

| Step | Reagent | Acronym | Storage | Catalog number | Vendor |
|---|---|---|---|---|---|
| Polymerizations | 40% Acrylamide | AAm | 4°C | A9099 | Sigma-Aldrich |
| | N,N'-(1,2-Dihydroxyethylene)bisacrylamide | DHEBA | 4°C | 294381 | Sigma-Aldrich |
| | N,N'-(1,2-Dihydroxyethylene)bisacrylamide | DHEBA | 4°C | sc-215503 | SCBT* |
| | Sodium acrylate | SA | -20°C desiccated | 408220 | Sigma-Aldrich |
| | Sodium acrylate | SA | -20°C desiccated | sc-236893C | SCBT* |
| | N,N′-Methylenebis(acrylamide) | BIS | RT | 146072 | Sigma-Aldrich |
| | Ammonium persulfate | APS | RT desiccated | AB00112 | American Bio |
| | Ammonium persulfate | APS | RT desiccated | A3678 | Sigma-Aldrich |
| | N,N,N′,N′-Tetramethylethylenediamine | TEMED | RT desiccated | AB02020 | American Bio |
| | 10X Phosphate buffered saline (Gibco) | 10X PBS | RT | 70011044 | Thermofisher |
| Buffers | Sodium hydroxide | NaOH | RT | S8045 | Sigma-Aldrich |
| | 20% Sodium dodecyl sulfate solution | SDS | RT | AB01922 | American Bio |
| | 5 M NaCl solution | NaCl | RT | AB1915 | American Bio |
| | Sodium Chloride | NaCl | RT | 3624-01 | J.T. Baker |
| | 1 M Tris solution, pH 8 | Tris | RT | AB14043 | American Bio |
| | Tris [hyroxymethyl] aminomethane | Tris | RT | AB02000 | American Bio |
| | Triton X-100 | TX-100 | RT | T8787 | Sigma-Aldrich |
| | Tween 20 | Tween 20 | RT | P7949 | Sigma-Aldrich |
| | NP-40 | NP-40 | RT | I8896 | Sigma-Aldrich |
| | 1X Phosphate buffered saline (Gibco) | 1X PBS | RT | 10010023 | Thermofisher |
| | Sodium Bicarbonate | - | RT | S5761 | Sigma-Aldrich |
| | Bovine Serum Albumin | BSA | 4°C | 001-000-162 | Jackson IR |
| | Normal Goat Serum | NGS | 4°C | 005-000-121 | Jackson IR |
| | Guanidine hydrochloride | G-HCl | RT | G3272 | Sigma-Aldrich |
| Fixatives | 16% Paraformaldehyde | FA | RT | 15710 | EMS** |
| | 8% Glutaraldehyde | GA | 4°C | 16019 | EMS** |

* Santa Cruz Biotechnology; ** Electron Microscopy Sciences

### SUPP TABLE. 2

**Supplementary Table 2:** Materials used

| Materials | Vendor | Catalog number |
|---|---|---|
| No. 1.5 12-mm round glass coverslips | Electron Microscopy Sciences | 72230-01 |
| Glass microscope slide | Sigma-Aldrich | S8400 |
| No. 1.5 22 x 22 mm square cover glass coverslips | Fisher Scientific | 12-541BP |
| No. 1.5 18-mm round coverslip | Marienfeld | 0117580 |
| 50 mm MatTek dish, No. 1.5 coverslip 30-mm diameter | MatTek Life Sciences | P50G-1.5-30-F |
| Picodent Twinsil | Picodent | 1300 1000 |

### SUPP TABLE. 3

**Supplementary Table 3:** Primary antibodies used to immunolabel neuron and brain tissue samples

| Primary antibody | Target | Host species | Dilution | Catalog number | Vendor |
|---|---|---|---|---|---|
| Recombinant anti-Homer1 antibody [EPR15309] | Homer 1 | Rb mAb | 1:500 (**N**); 1:250 (**B**) | ab184955 | abcam |
| Anti-Bassoon/BSN antibody [SAP7F407] | Bassoon | Ms mAb | 1:500 (**N**); 1:250 (**B**) | ab82958 | abcam |
| Anti-PSD-95 Antibody [K28/43] | PSD-95 | Ms mAb | 1:500 (**N**) 1:250 (**B**) | 75-028 | antibodiesinc |
| Anti-Synaptophysin Antibody (D8F6H) | Synaptophysin | Rb mAb | 1:250 (**N**) | 36406 | CST |
| Anti-Synaptophysin 1 antibody | Synaptophysin | Ms mAb | 1:250 (**N**) | 101 011 | SYSY |
| Anti-GFAP polyclonal antibody | GFAP | Rb pAb | 1:500 (**B**) | PA1-10019 | Thermofisher |
| Purified anti-Myelin Basic Protein Antibody | MBP | Ms mAb | 1:500 (**B**) | 808401 | BioLegend |
| Anti-GFP polyclonal antibody | GFP | Rb pAb | 1:500 (**B**) | A-11122 | Thermofisher |

**N**: cultured neuron samples, **B**: brain tissue section samples

### SUPP TABLE. 4

**Supplementary Table 4:** Secondary antibodies used to immunolabel neuron and brain tissue samples

| Secondary dye | Species reactivity | Host | Dilution | Ex/Em [nm] | Catalog number | Vendor |
|---|---|---|---|---|---|---|
| ATTO594 | Mouse | Goat | 1:250-1:500 | 603/626 | 76085 | Sigma-Aldrich |
| ATTO594 | Rabbit | Goat | 1:250-1:500 | 603/626 | 77671 | Sigma-Aldrich |
| ATTO647N | Mouse | Goat | 1:250-1:500 | 646/664 | 50185 | Sigma-Aldrich |
| ATTO647N | Rabbit | Goat | 1:250-1:500 | 646/664 | 40839 | Sigma-Aldrich |
| CF568 | Rabbit | Donkey | 1:250 | 562/583 | 20098 | Biotium |

### SUPP TABLE. 5

**Supplementary Table 5:** Fluorescent pan-stains and lipid probes used

| Stain/Pan-stain | Concentration | Ex/Em [nm] | Catalog number | Vendor |
|---|---|---|---|---|
| ATTO532, NHS ester | 20 µg/mL (**N**); 30 µg/mL (**B**) | 532/552 | 88793 | Sigma-Aldrich |
| ATTO532, NHS ester | 20 µg/mL (**N**); 30 µg/mL (**B**) | 532/552 | AD 532-31 | ATTO-TEC |
| ATTO594, NHS ester | 20 µg/mL (**N**); 30 µg/mL (**B**) | 603/626 | 08741 | Sigma-Aldrich |
| ATTO594, NHS ester | 20 µg/mL (**N**); 30 µg/mL (**B**) | 603/626 | AD 594-31 | ATTO-TEC |
| ATTO488, NHS ester | 20 µg/mL (**N**) | 500/520 | 41698 | Sigma-Aldrich |
| SYTOX Green | 1 µM (**N**); 1.7 µM (**B**) | 504/523 | S7020 | Thermofisher |
| BODIPY TR Methyl Ester | 10 µM (**B**) | 588/621 | C34556 | Thermofisher |
| ATTO590, alkyne | 5 µM (**N**) | 594/624 | 93990 | Sigma-Aldrich |
| ATTO590, azide | 5 µM (**N**) | 593/622 | AD 590-101 | ATTO-TEC |
| Palmitic acid azide | 50 µM (**N**) | NA | C10265 | Thermofisher |
| PhotoClick Sphingosine (pacSph) | 50 µM (**B**) | NA | 900600 | Avanti Polar Lipids |
| Click-IT™ Cell Reaction Buffer Kit | NA | NA | C10269 | Thermofisher |

**N**: cultured neuron samples, **B**: brain tissue section samples

SUPP FIG. 1

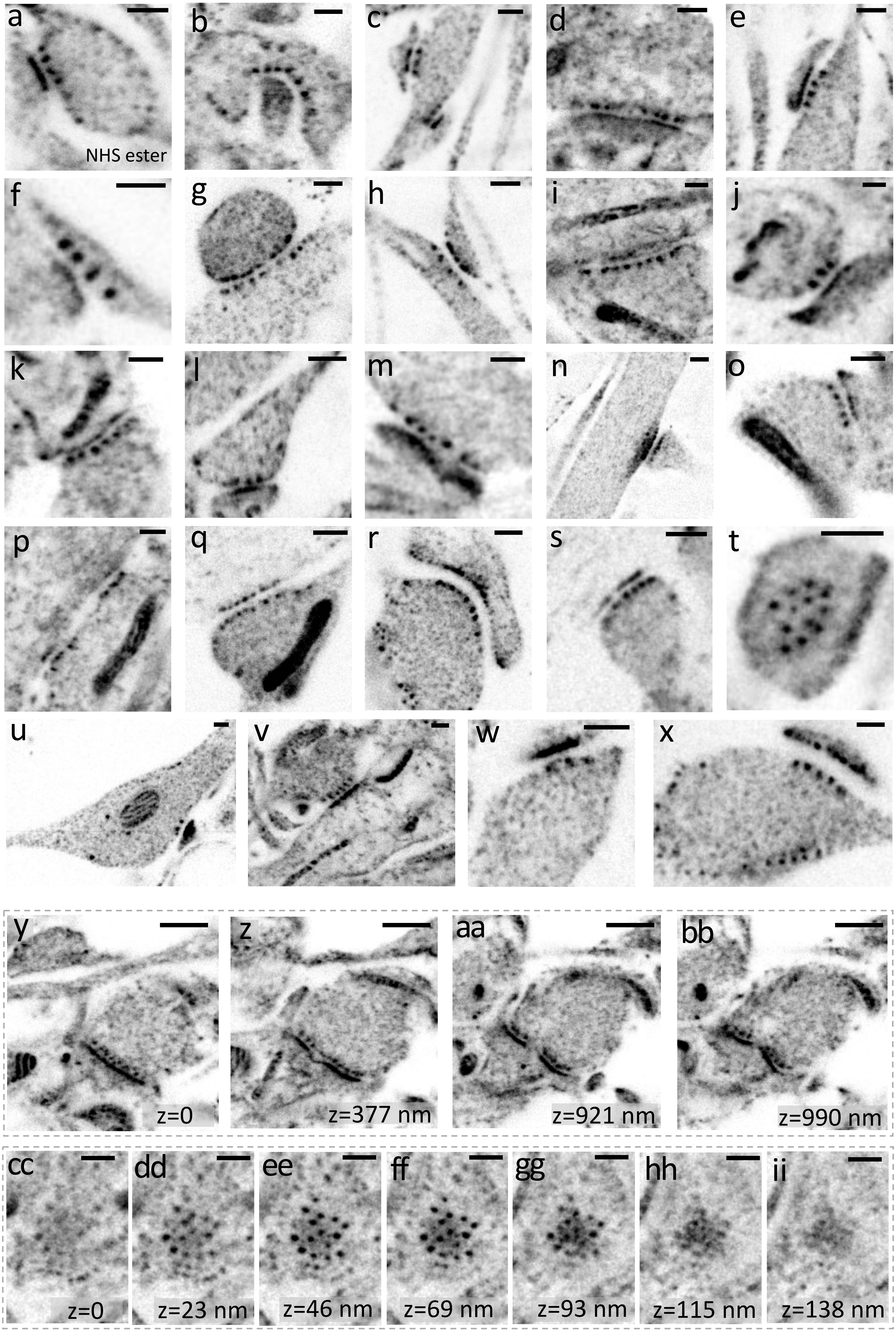
a
b
c
d
e
NHS ester
f
g
h
i
j
k
l
m
n
o
p
q
r
s
t
u
v
w
x
y
z
aa
bb
z=0
z=377 nm
z=921 nm
z=990 nm
cc
dd
ee
ff
gg
hh
ii
z=0
z=23 nm
z=46 nm
z=69 nm
z=93 nm
z=115 nm
z=138 nm

**Supplementary Figure 1: Gallery of NHS ester pan-stained synapses in dissociated mouse and rat hippocampal neurons. a-z,** axial or top views of synapses showing presynaptic dense projections (DPs) and postsynaptic densities (PSDs). **y-bb,** z-stack of a synapse (side view) spanning 990 nanometers. **cc-ii**, z-stack of a synapse (top view) spanning 138 nanometers. Image **d** was processed with a gamma value ɣ=0.5; image **l** with ɣ=0.6; images **b**, **g, i,** and **u** with ɣ=0.7; images **h**, **j**, **k**, **m**, **n**, **o**, **q**, **s**, **t**, **v**, **w**, **x**, **cc-ii** with ɣ=0.8; and image **r** with ɣ=0.9. Scale bars are corrected for the expansion factor, (**a-ii**) 200 nm.

SUPP FIG. 2

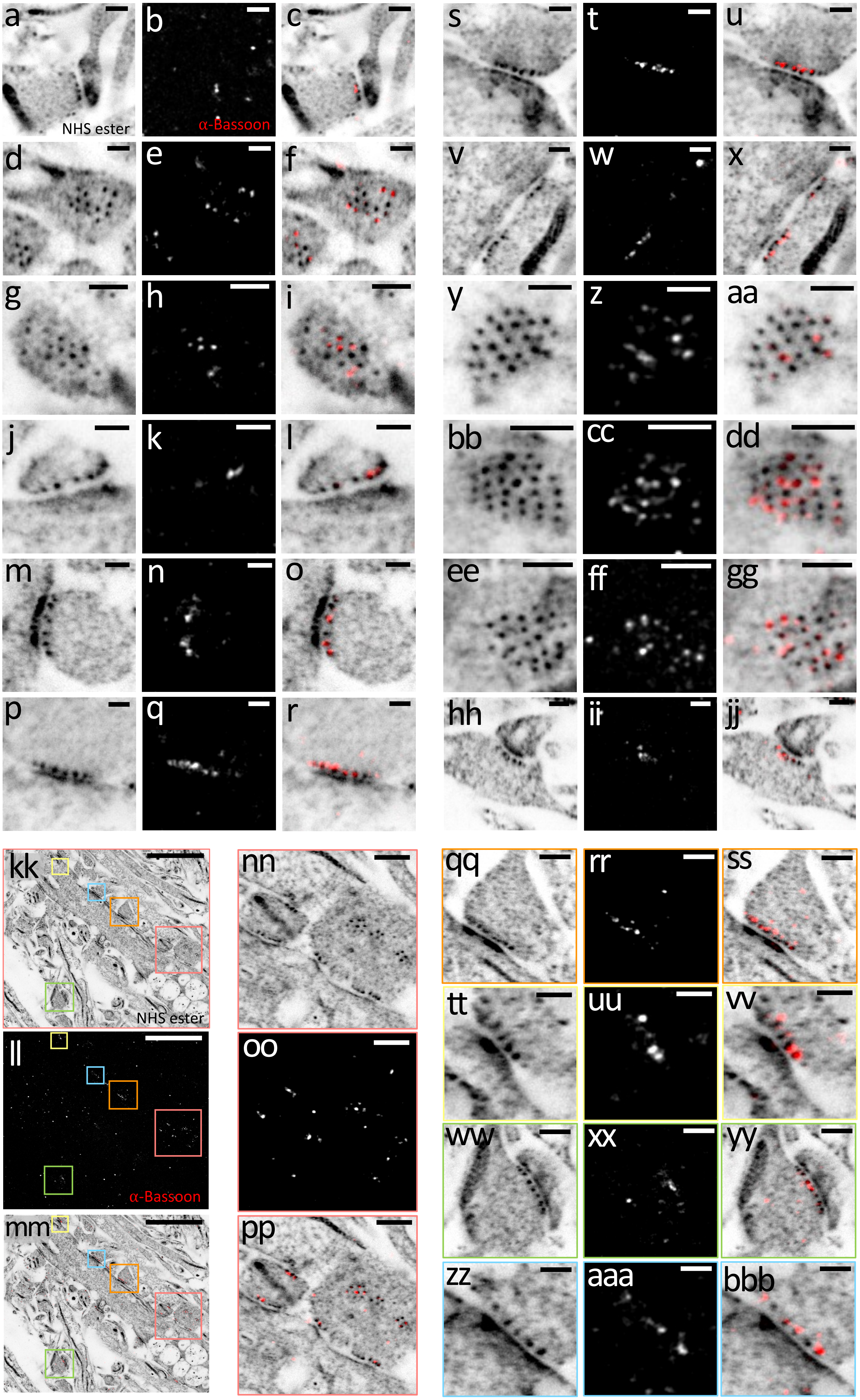
a
b
c
d
e
f
g
h
i
j
k
l
m
n
o
p
q
r
s
t
u
v
w
x
y
z
aa
bb
cc
dd
ee
ff
gg
hh
ii
jj
kk
ll
mm
nn
oo
pp
qq
rr
ss
tt
uu
vv
ww
xx
yy
zz
aaa
bbb
NHS ester
α-Bassoon
NHS ester
α-Bassoon

**Supplementary Figure 2: Gallery of Bassoon-immunolabeled and NHS ester pan-stained synapses in dissociated mouse hippocampal neurons.** **a**, **d**, **g**, **j**, **m**, **p**, **s**, **v**, **y**, **bb**, **ee**, **hh**, axial or top views of synapses pan-stained with NHS ester. **b**, **e**, **h**, **k**, **n**, **q**, **t**, **w**, **z**, **cc**, **ff**, **ii**, Bassoon immunolabeling distributed near dense projections. **c**, **f**, **i**, **l**, **o**, **r**, **u**, **x**, **aa**, **dd**, **gg**, **jj**, overlay. **kk**, ~35 µm FOV image of NHS ester pan-stained neurites. **ll**, Bassoon immunolabeling of the same area. **mm**, overlay of **kk** and **ll**. **nn**, **oo**, **pp**, magnified views of the areas outlined in the salmon boxes in **kk**, **ll**, and **mm**, respectively. **qq**, **rr**, **ss**, magnified views of the areas outlined by the orange boxes in **kk, ll,** and **mm,** respectively. **tt**, **uu**, **vv**, magnified views of the areas outlined by the yellow boxes in **kk**, **ll**, and **mm**, respectively. **ww**, **xx**, **yy**, magnified views of the areas outlined by the lime boxes in **kk**, **ll**, and **mm**, respectively. **zz**, **aaa**, **bbb**, magnified views of the areas outlined by the light blue boxes in **kk**, **ll**, and **mm**, respectively. Images **d**, **v**, **hh**, **kk**, **nn**, **qq**, **tt**, **ww**, and **zz**, were processed with a gamma value ɣ=0.7 and image **a** with ɣ=0.8. Scale bars are corrected for the expansion factor, (**a-jj**, **nn-pp**) 200 nm. (**kk-mm**) 5 µm.

### SUPP FIG. 3

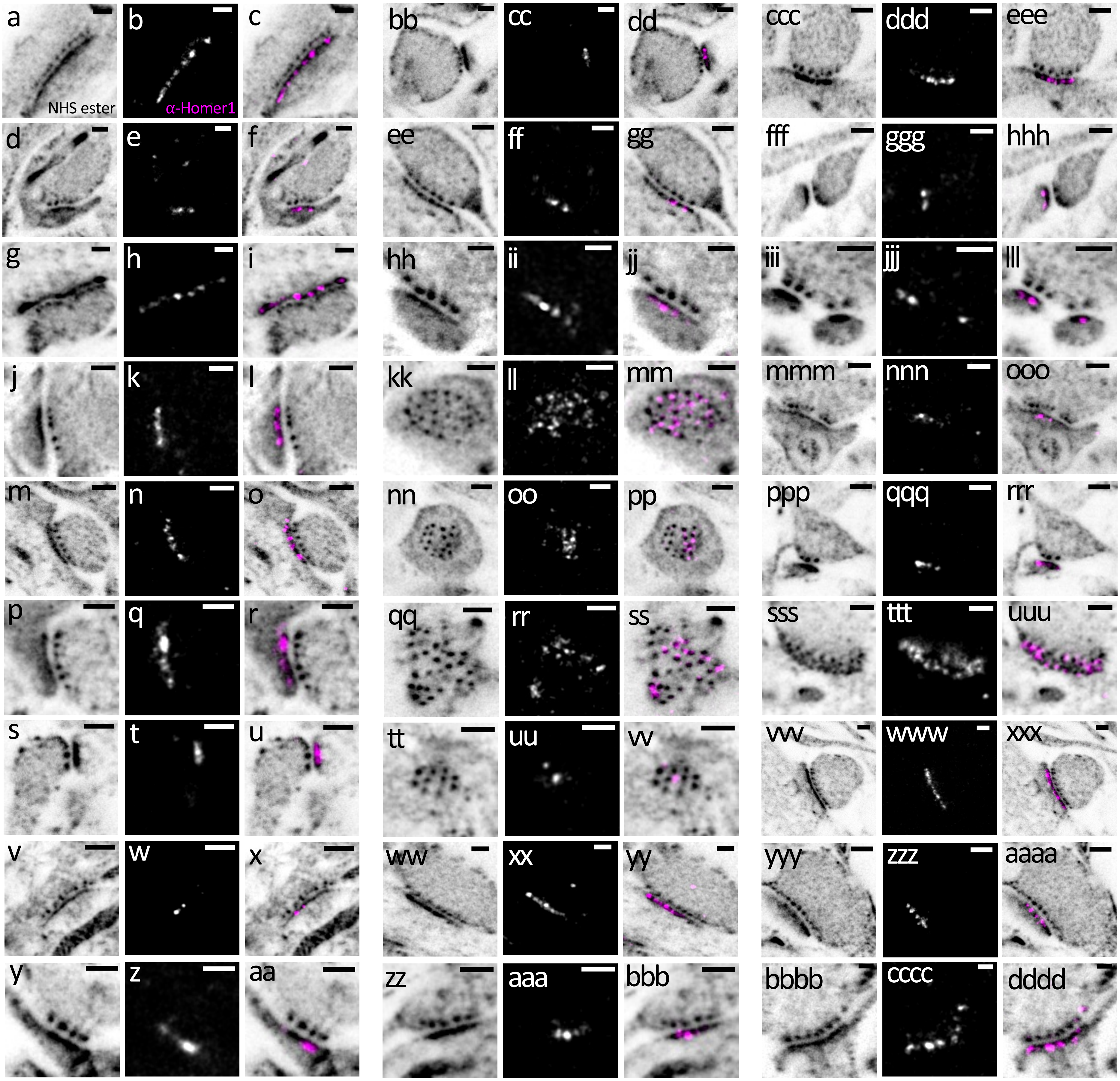


**Supplementary Figure 3: Gallery of Homer1-immunolabeled and NHS ester pan-stained synapses in dissociated mouse neurons. a, d, g, j, m, p, s, v, y, bb, ee, hh, kk, nn, qq, tt, ww, zz, ccc, fff, iii, mmm, ppp, sss, vvv, yyy, bbbb**, axial or top views of synapses pan-stained with NHS ester. **b, e, h, k, n, q**, t, **w, z, cc, ff, ii, ll, oo, rr, uu, xx, aaa, ddd, ggg, jjj, nnn, qqq, ttt, www, zzz, cccc**, Homer1 immunolabeling localized in the postsynaptic density. **c, f, i, l, o, r, u, x, aa, dd, gg, jj, mm, pp, ss, vv, yy, bbb, eee, hhh, lll, ooo, rrr, uuu, xxx, aaaa, dddd**, overlay. Images **j, nn, tt, ccc, zz, ppp, fff, sss**, and **yyy** were processed with a gamma value ɣ=0.8. Scale bars are corrected for the expansion factor, (**a-dddd**) 200 nm.

### SUPP FIG. 4

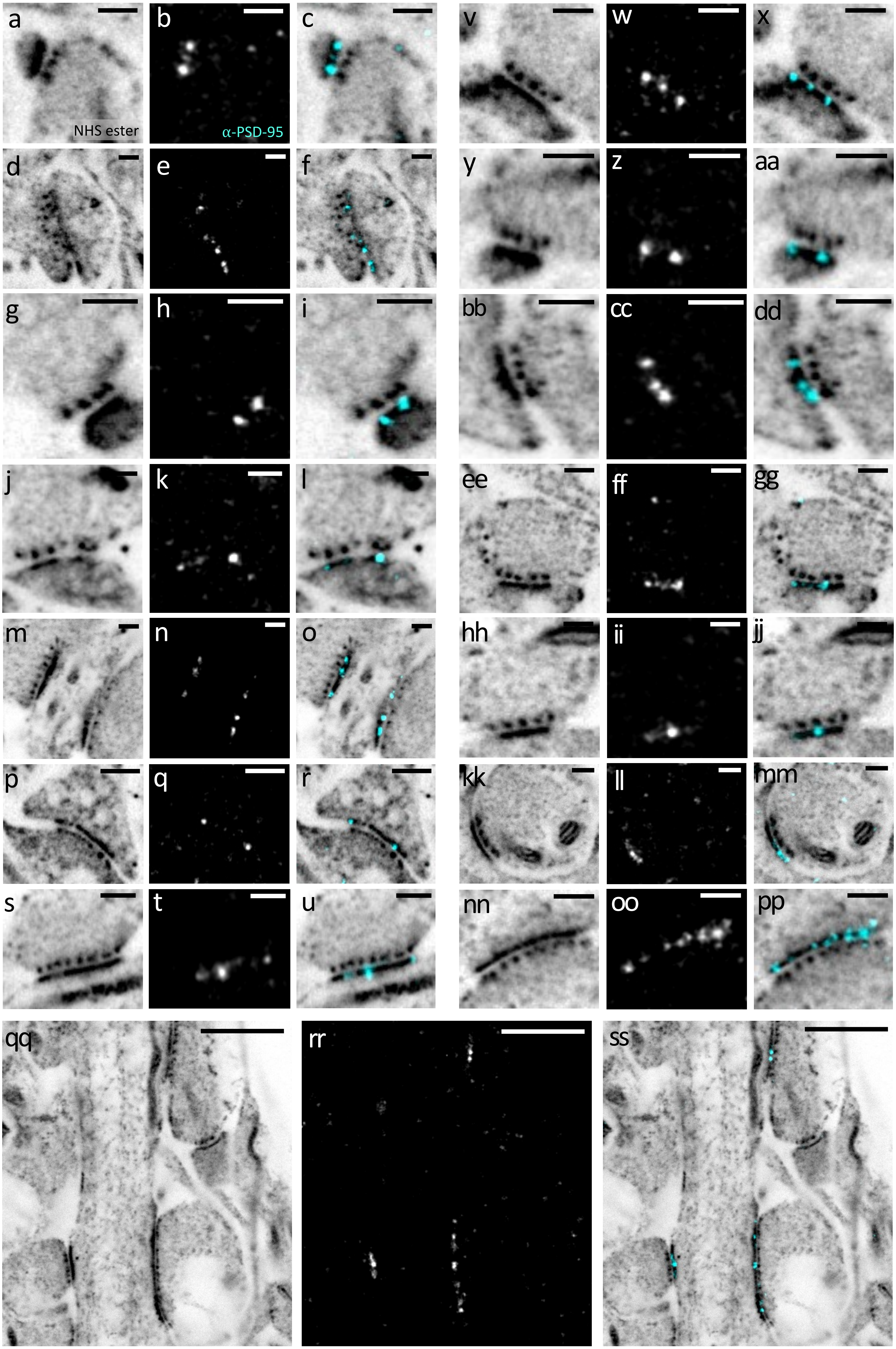
a
b
c
v
w
x
NHS ester
α-PSD-95
d
e
f
y
z
aa
g
h
i
bb
cc
dd
j
k
l
ee
ff
gg
m
n
o
hh
ii
jj
p
q
r
kk
ll
mm
s
t
u
nn
oo
pp
qq
rr
ss

**Supplementary Figure 4: Gallery of PSD-95 immunolabeled and NHS ester pan-stained synapses in dissociated mouse hippocampal neurons.** **a**, **d**, **g**, **j**, **m**, **p**, **s**, **v**, **y**, **bb**, **ee**, **hh**, **kk**, **nn**, **qq**, axial or top views of synapses pan-stained with NHS ester. **b**, **e**, **h**, **k**, **n**, **q**, **t**, **w**, **z**, **cc**, **ff**, **ii**, **ll**, **oo**, **rr**, PSD-95 immunolabeling localized near dense projections. **c**, **f**, **i**, **l**, **o**, **r**, **u**, **x**, **aa**, **dd**, **gg**, **jj**, **mm**, **pp**, **ss**, overlay. Images **m** and **kk** were processed with a gamma value ɣ=0.7 and images **a**, **d**, **g**, **j**, **p**, **s**, **v**, **y**, **ee**, **hh,** and **nn** with ɣ=0.8. Scale bars are corrected for the expansion factor, (**a-pp**) 200 nm. (**qq-ss**) 1 µm.

SUPP FIG. 5

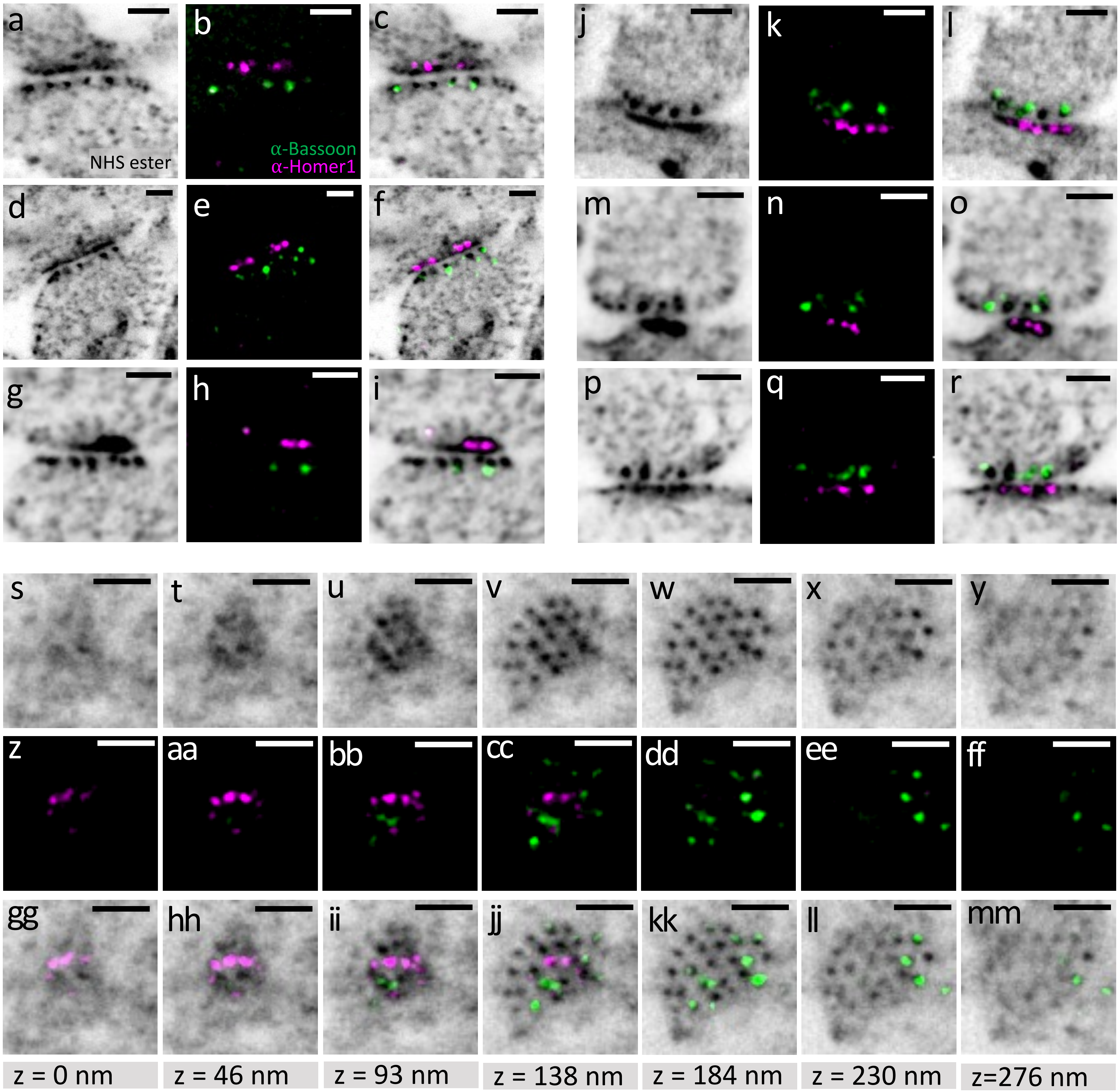


**Supplementary Figure 5: Gallery of Homer1- and Bassoon-immunolabeled and NHS ester pan-stained synapses in dissociated mouse hippocampal neurons. a**, **d**, **g**, **j**, **m**, **p**, axial or top views of synapses pan-stained with NHS ester. **b**, **e**, **h**, **k**, **n**, **q**, Homer1 and Bassoon immunolabeling. **c**, **f**, **i**, **l**, **o**, **r**, overlay. **s-u**, z-stack of a synapse (top view) spanning 276 nanometers and pan-stained with NHS ester. **z-ff**, Homer1 and Bassoon immunolabeling of the same areas as **s-u**. **gg-mm**, overlay. Images **a, d, j** and **s-y** were processed with a gamma value ɣ=0.7. Scale bars are corrected for the expansion factor, (**a-mm**) 200 nm.

### SUPP FIG. 6

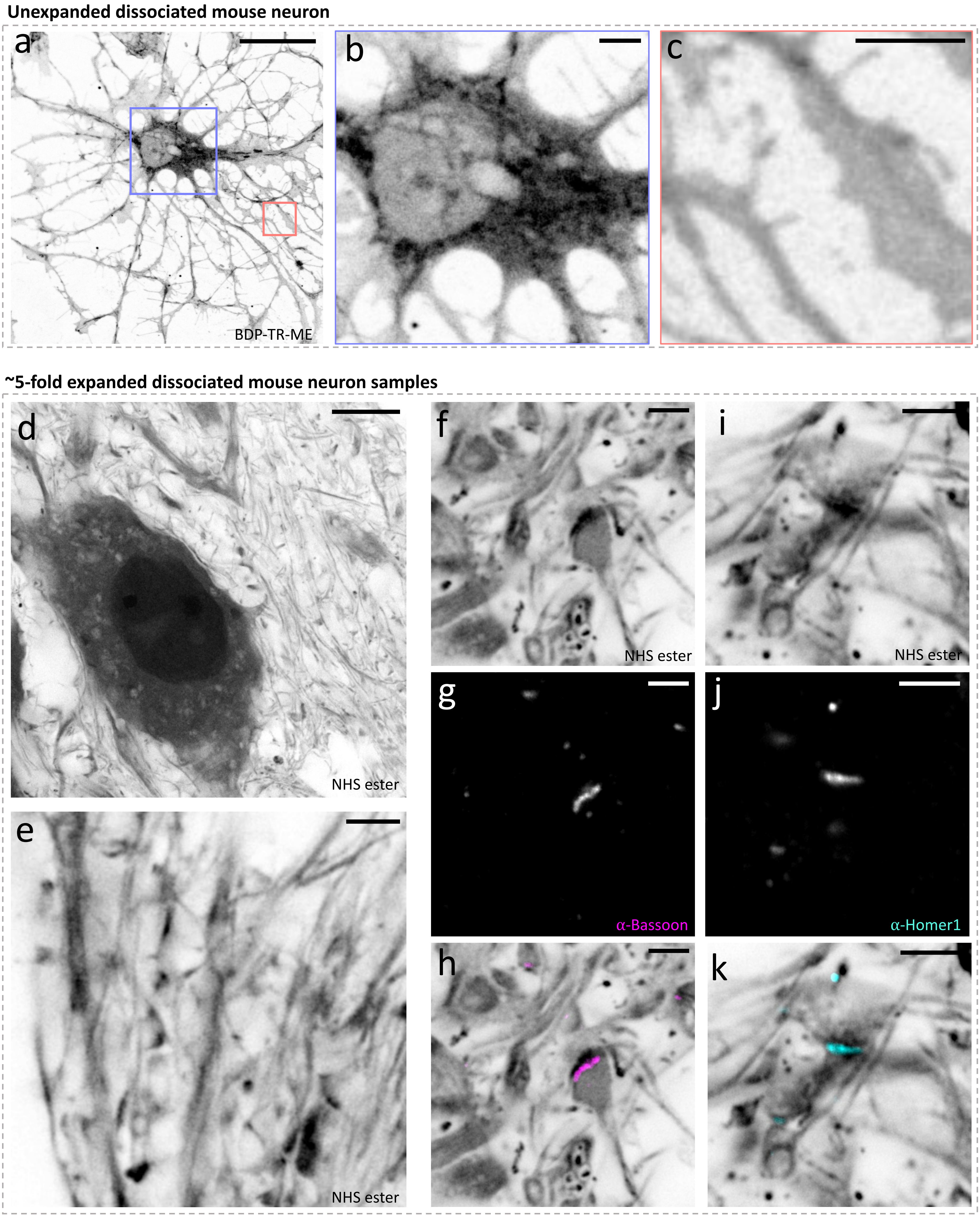

Unexpanded dissociated mouse neuron
a
b
c
BDP-TR-ME
~5-fold expanded dissociated mouse neuron samples
d
NHS ester
e
NHS ester
f
NHS ester
g
α-Bassoon
h
i
NHS ester
j
α-Homer1
k

**Supplementary Figure 6: Ultrastructural features in neurons are not visible in non-expanded and ~5 fold expanded samples. a,** non-expanded neuron pan-stained with BDP-TR-ME. **b,** magnified view of the area outlined in the lavender box in **a** showing a neuron soma. **c,** magnified view of the area outlined by the salmon box in **a** showing neurites. **d**, ~5-fold expanded neurons pan-stained with NHS ester showing a neuron soma. **e**, ~5-fold expanded neurons pan-stained with NHS ester showing neurites. **f**, **i**, ~5-fold expanded neurons pan-stained with NHS ester showing synapses. **g**, **j**, Bassoon and Homer1 immunolabeling of the same areas shown in **f** and **i**. **h**, **k**, overlay. Images **d**, **e**, **f**, and **i** were processed with a gamma value ɣ=0.8. Scale bars are corrected for the expansion factor, (**a**) 20 µm, (**b**, **c**, **e-k**) 1 µm, (**d**) 10 µm.

### SUPP FIG. 7

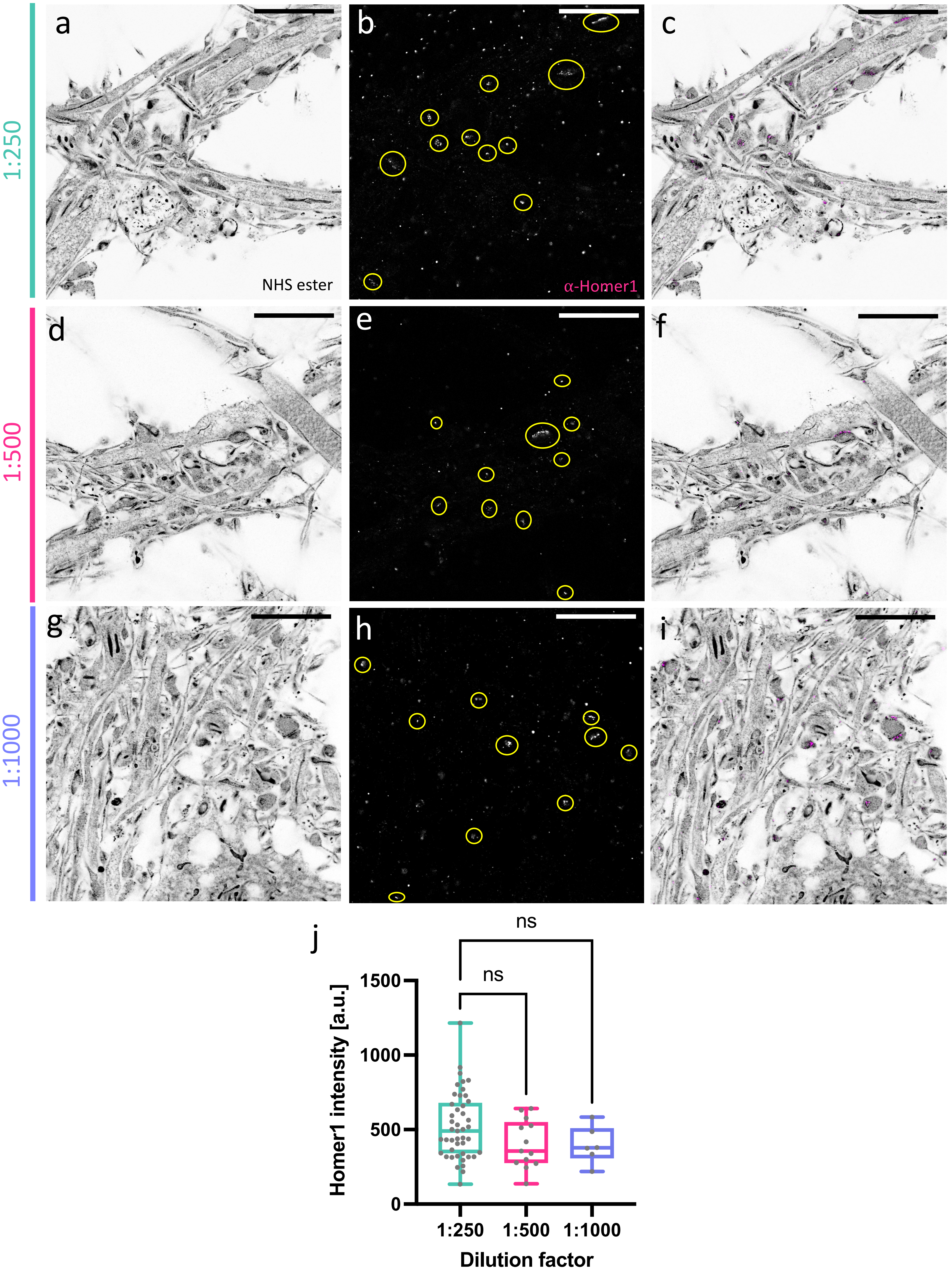

1:250
1:500
1:1000
a
b
c
NHS ester
α-Homer1
d
e
f
g
h
i
j
ns
ns
1500
1000
500
0
Homer1 intensity [a.u.]
1:250
1:500
1:1000
Dilution factor

**Supplementary Figure 7: Effect of antibody dilution factor on background noise in expanded mouse neurons.** **a**, **d**, **g**, pan-ExM processed neurons pan-stained with NHS ester. **b**, **e**, **h**, Homer1 immunolabeling of the areas shown in **a**, **d**, and **g**, respectively. Yellow circles show 'true' Homer1 signal. **c**, **f**, **i**, overlay. **j**, intensity measurements of Homer1 peak signal according to the dilution factors of both primary and secondary antibodies (1:250: n = 45 measurements from 7 FOVs; 1:500: n = 13 measurements from 3 FOVs; 1:1000: n = 6 measurements from 1 FOV). ns: non-significant. Images **a**, **d**, and **g** were processed with a gamma value $\gamma$=0.8. Scale bars are corrected for the expansion factor, (**a-i**) 4 µm.

### SUPP FIG. 8

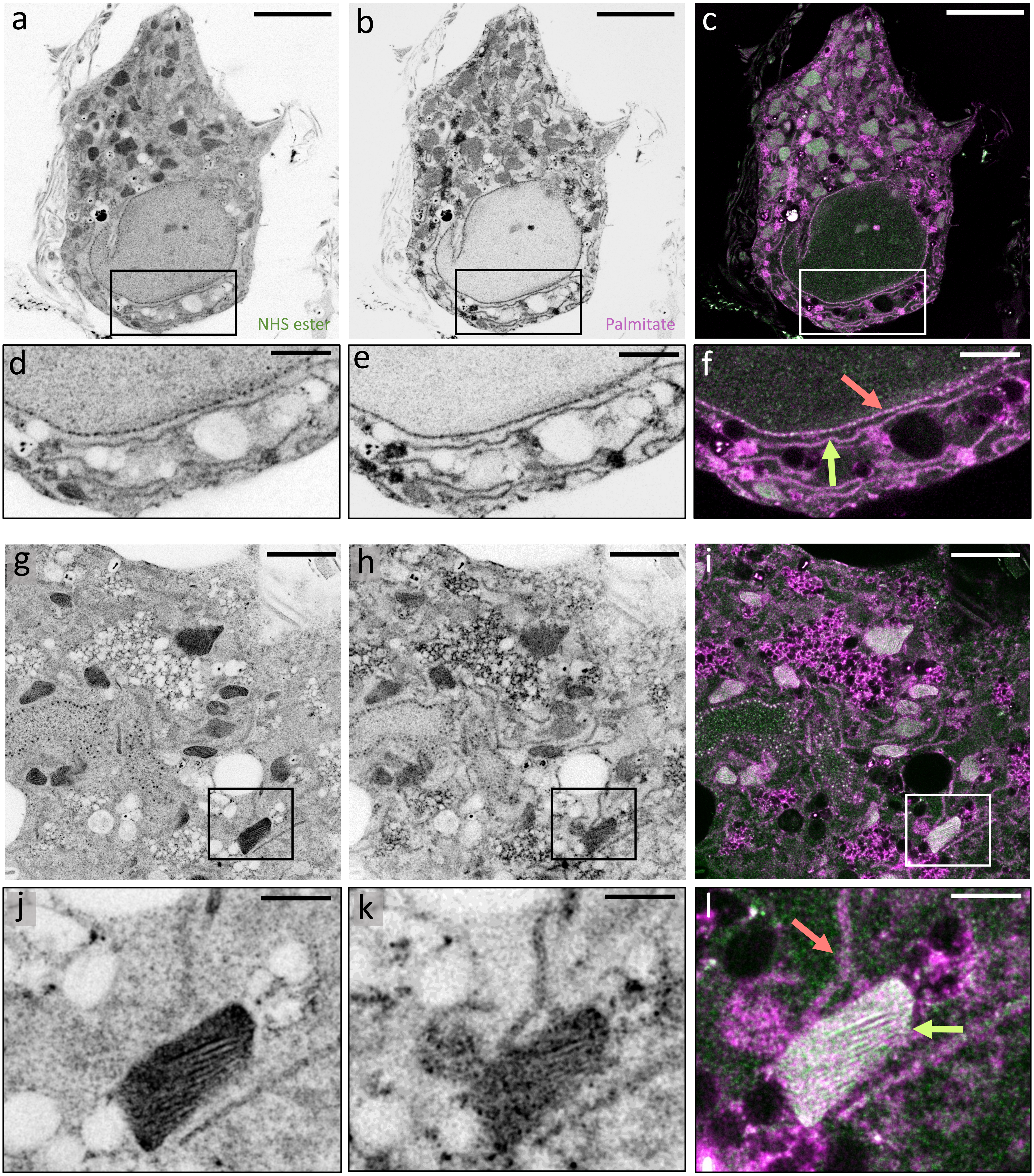

**Supplementary Figure 8: Palmitate and NHS ester differential pan-staining in dissociated mouse neurons.** **a**, NHS ester pan-stained neuron soma. **b**, palmitate pan-staining of the same area shown in **a**. **c**, overlay. **d-f**, magnified views of the areas outlined in the black boxes in **a-c**. Salmon arrow in f points to the nuclear envelope and lime arrow points to an ER tubule. **g**, NHS ester pan-stained neuron soma cytosol. **h**, palmitate pan-staining of the same area shown in **g**. **i**, overlay. **j-l**, magnified views of the areas outlined in the black boxes in **g-i.** Salmon arrow in **l** points to an ER tubule that is contacting a mitochondrion (lime arrow). Image **g** was processed with a gamma value ɣ=0.8. Scale bars are corrected for the expansion factor, (**a-c**) 5 µm, (**d-f**, **g-i**) 2 µm, (**j-l**) 500 nm.

### SUPP FIG. 9

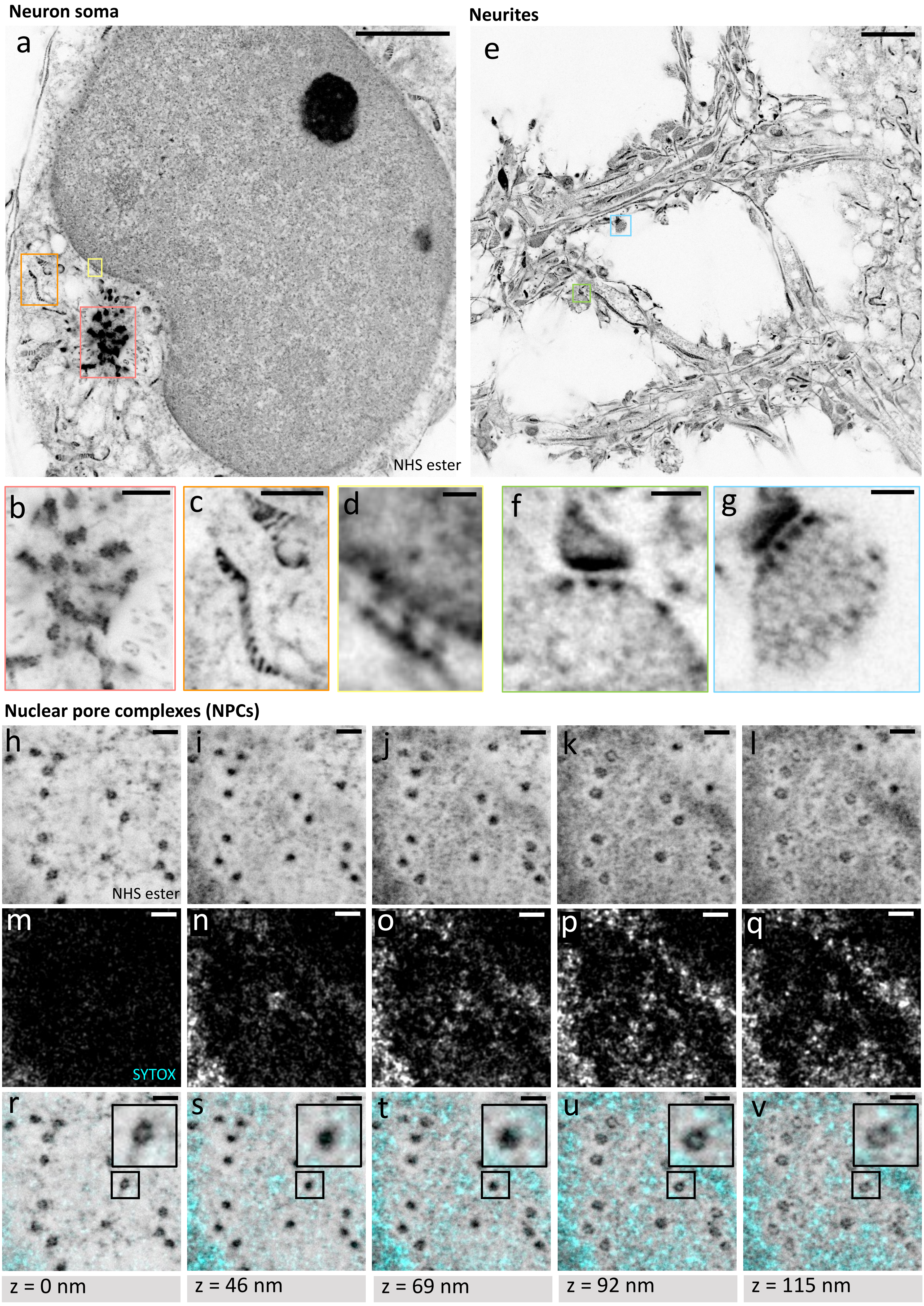

Neuron soma
Neurites
a
e
NHS ester
b
c
d
f
g
Nuclear pore complexes (NPCs)
h
i
j
k
l
NHS ester
m
n
o
p
q
SYTOX
r
s
t
u
v
z = 0 nm
z = 46 nm
z = 69 nm
z = 92 nm
z = 115 nm

**Supplementary Figure 9: Gallery of subcellular features in dissociated mouse hippocampal neurons.** **a**, neuron soma processed with pan-ExM and pan-stained with NHS ester. **b**, **c**, **d**, magnified views of the areas in **a** outlined by the salmon, orange, and yellow boxes showing a centrosome, mitochondria and the side view of nuclear pore complexes (NPCs) respectively. **e**, neuron neurites processed with pan-ExM and pan-stained with NHS ester. **f**, **g**, magnified views of the areas in **e** outlined by the lime and light blue boxes showing synapses. **h-l**, z-stack of nuclear pore complexes pan-stained with NHS ester (top view) spanning 115 nanometers. **m-q**, SYTOX Green staining of the same areas as in **h-l**. **r-v**, overlay. Insets in **r** and **u** show hollow rings corresponding to the cytoplasmic and nuclear surface of the NPC respectively. Image **a** was processed with a gamma value ɣ=0.8 and image **e** with ɣ=0.7. Scale bars are corrected for the expansion factor, (**a**, **e**) 3 µm, (**b-c**) 500 nm, (**d**) 100 nm, (**f-v**) 200 nm.

SUPP FIG. 10

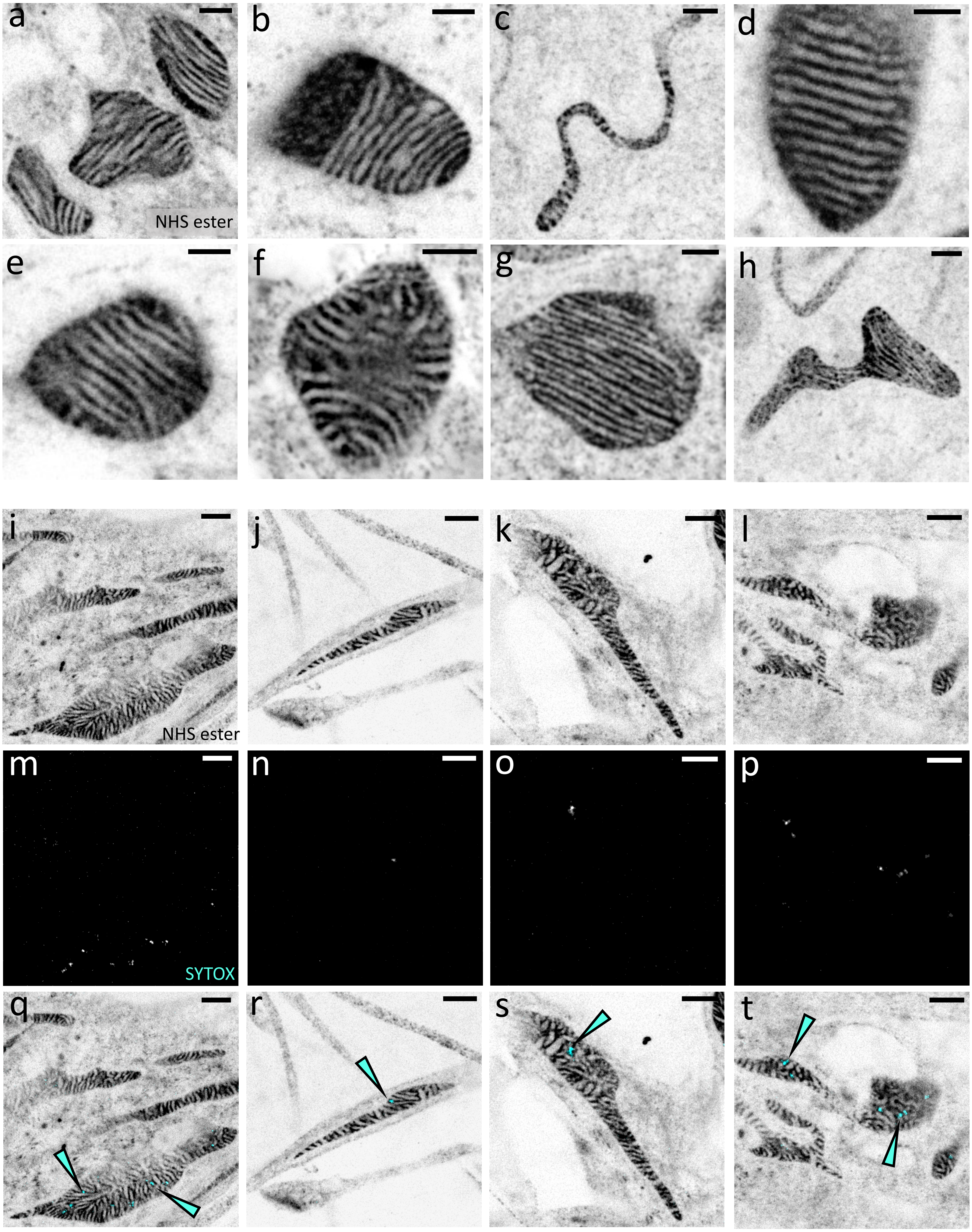


**Supplementary Figure 10: Gallery of mitochondria in dissociated mouse and rat neurons. a-h**, NHS ester pan-stained mitochondria showing resolvable cristae. **i-l**, mitochondria in neuron somas (**i**, **k**, **l**) and a neurite (**j**) pan-stained with NHS ester. **m-p**, SYTOX Green staining of the same areas shown in **i-l**. **q-t**, overlay. Cyan arrows point at SYTOX Green-stained mtDNA. Images **a**, **b**, **d**, **e**, **i**, **j**, **k**, and **l** were processed with a gamma value ɣ=0.8. Scale bars are corrected for the expansion factor, (**a-h**) 200 nm, (**i-t**) 500 nm

SUPP FIG. 11

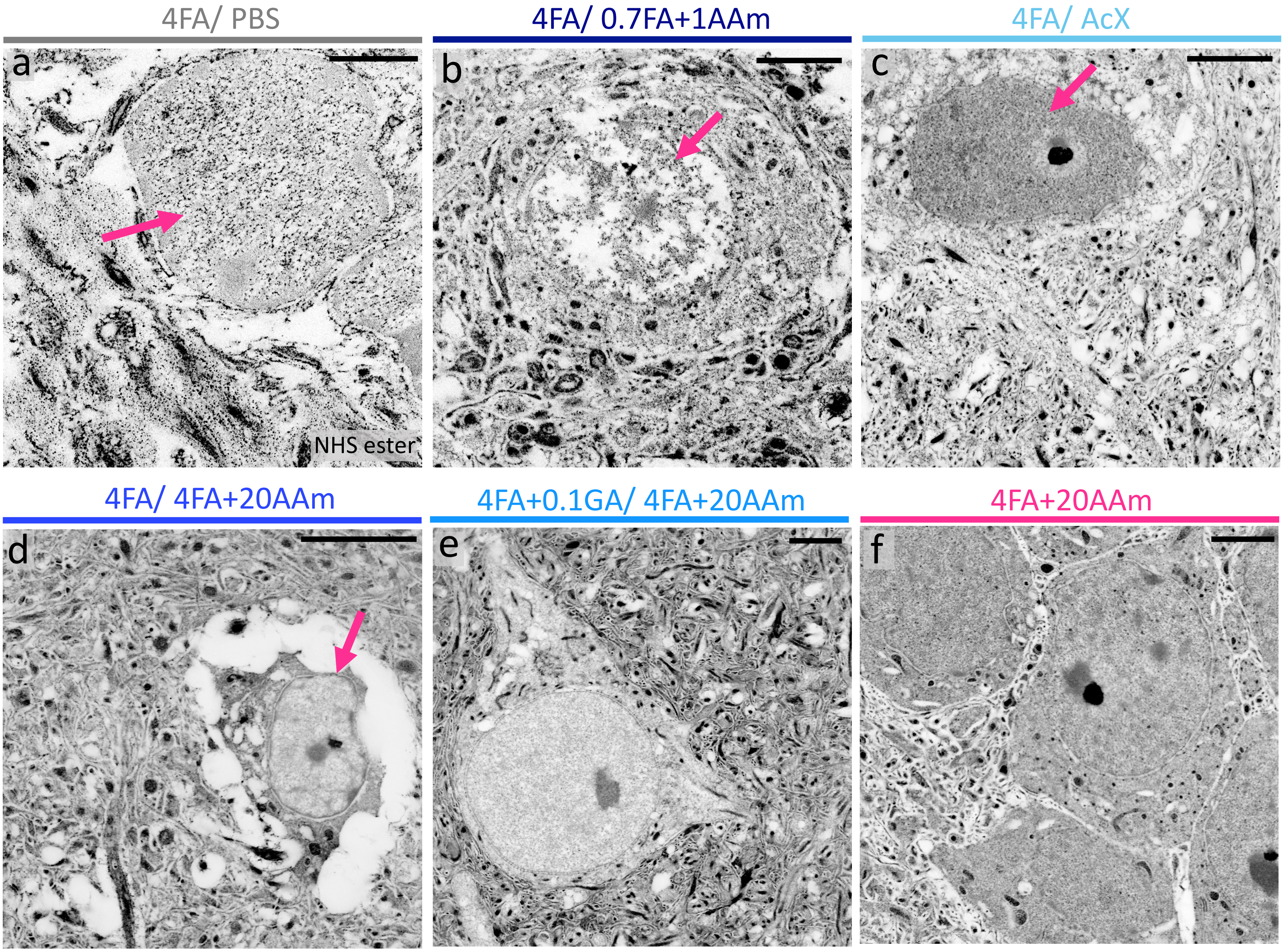


**Supplementary Figure 11: Effect of fixation and post-fixation on cell ultrastructure in expanded mouse brain tissue sections. a**, NHS ester pan-stained brain tissue fixed with 4% FA and not post-fixed (stored in PBS). Only the cell nucleus is discernable (pink arrow). **b**, NHS ester pan-stained brain tissue fixed with 4% FA and post-fixed with 0.7% FA + 1% AAm. Cell nucleus shows major extraction artifacts (pink arrow). **c**, NHS ester pan-stained brain tissue fixed with 4% FA and post-treated with 0.1 mg/mL AcX. Cell nucleus (pink arrow) appears highly stained, but cytosol and surrounding neuropil present extraction artifacts. **d**, NHS ester pan-stained brain tissue fixed with 4% FA and post-fixed with 4% FA + 20% AAm. Cell (pink arrow) is detached from surrounding tissue. **e**, NHS ester pan-stained brain tissue fixed with 4% FA + 0.1% GA and post-fixed with 4% FA + 20% AAm showing adequate cell preservation. **f**, NHS ester pan-stained brain tissue fixed with 4% FA + 20% AAm showing adequate cell preservation. Scale bars are not corrected for the expansion factor (**a-f**) 50 µm.

### SUPP FIG. 12

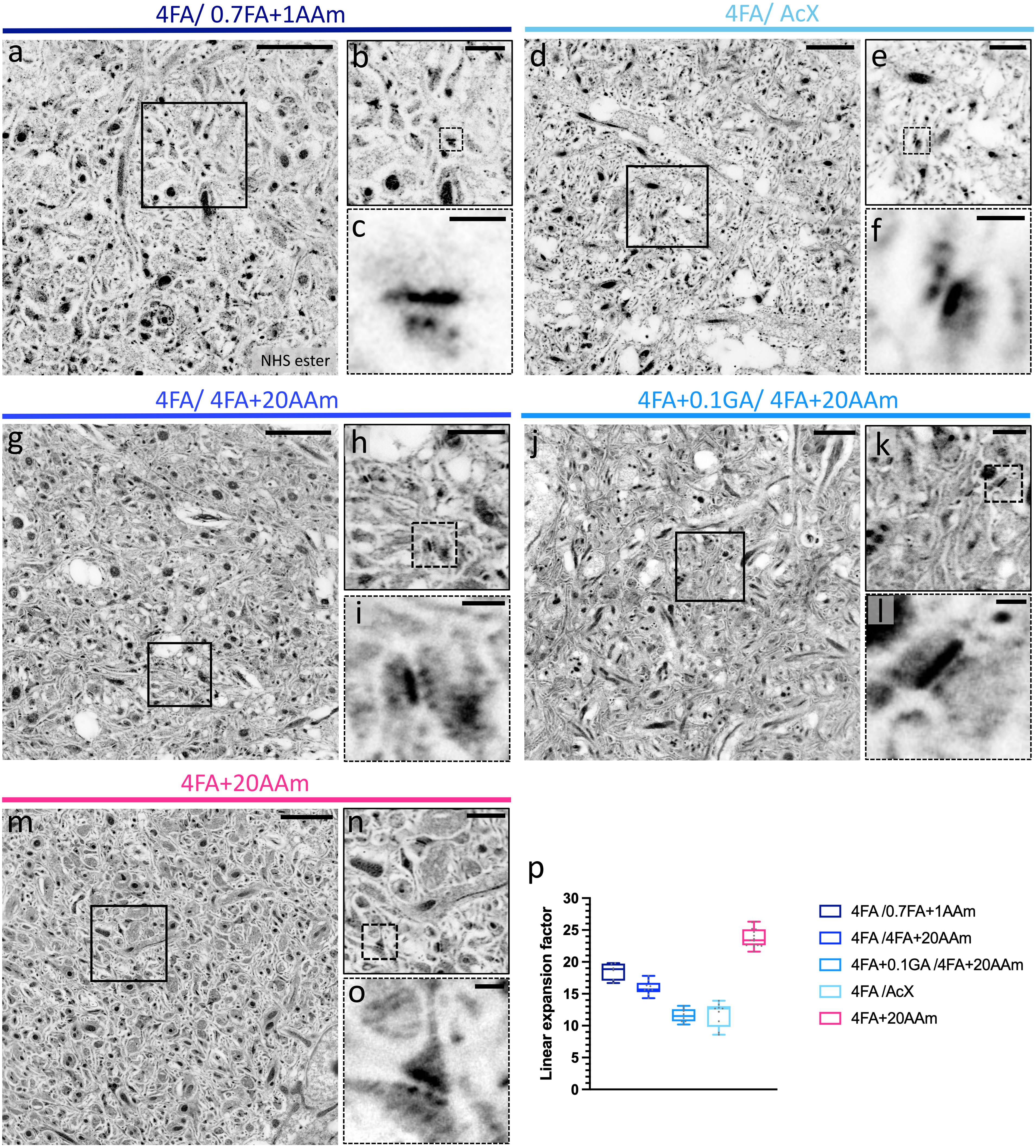


q

| Ref in text | Fix /Post-fix | Neuropil preservation | Cell body preservation | Expansion factor |
|---|---|---|---|---|
| *Fix-2* | 4FA /0.7FA+1AAm | + (low S/N) | + | ++ |
| *Fix-3* | 4FA /4FA+20AAm | ++ | + (detached) | ++ |
| *Fix-4* | 4FA+0.1GA /4FA+20AAm | +++ | +++ | + |
| *Fix-5* | 4FA /AcX | + | + | + |
| *Fix-6* | 4FA+20AAm | +++ | +++ | +++ |

**Supplementary Figure 12: Effect of fixation and post-fixation on neuropil ultrastructure in expanded mouse brain tissue sections.** **a**, NHS ester pan-stained brain tissue fixed with 4% FA and post-fixed with 0.7% FA + 1% AAm. **b**, magnified view of the area outlined in the black box in **a**. **c**, magnified view of the area outlined in the dotted black box in **b** showing a synapse. Neuropil in this condition has low signal to noise (difficult to distinguish neurites). **d**, NHS ester pan-stained brain tissue fixed with 4% FA and post-treated with 0.1 mg/mL AcX . **e**, magnified view of the area outlined in the black box in **d**. **f**, magnified view of the area outlined in the dotted black box in **e** showing a synapse. Neuropil in this condition is poorly preserved (presence of many artifactual gaps). **g**, NHS ester pan-stained brain tissue fixed with 4% FA and post-fixed with 4% FA + 20% AAm. **h**, magnified view of the area outlined in the black box in **g**. **i**, magnified view of the area outlined in the dotted black box in **h** showing a synapse. Neuropil in this condition is somewhat preserved (neurites distinguishable but presence of some artifactual gaps). **j**, NHS ester pan-stained brain tissue fixed with 4% FA + 0.1% GA and post-fixed with 4% FA + 20% AAm. **k**, magnified view of the area outlined in the black box in **j**. **l**, magnified view of the area outlined in the dotted black box in **k** showing a synapse. Neuropil in this condition is adequately preserved. **m**, NHS ester pan-stained brain tissue fixed with 4% FA + 20% AAm. **n**, magnified view of the area outlined in the black box in **m**. **o**, magnified view of the area outlined in the dotted black box in **n** showing a synapse. Neuropil in this condition is adequately preserved. **p**, plot of expansion factors (Fix-2: n = 38 measurements from 9 FOVs; Fix-3: n = 81 measurements from 9 FOVs; Fix-4: n = 77 measurements from 8 FOVs; Fix-5: 74 measurements from 10 FOVs; Fix-6: 254 measurements from 10 FOVs in 3 independent experiments). **q**, table outlining a qualitative assessment of mouse brain tissue ultrastructure preservation according to the different fixation schemes used. Black scale bars are corrected for the expansion factor (**a**, **d**, **g**, **f**, **m**) 3 µm, (**b**, **e**, **h**, **k**, **n**) 1 µm, (**c**, **f**, **i**, **l**, **o**) 200 nm.

### SUPP FIG. 13

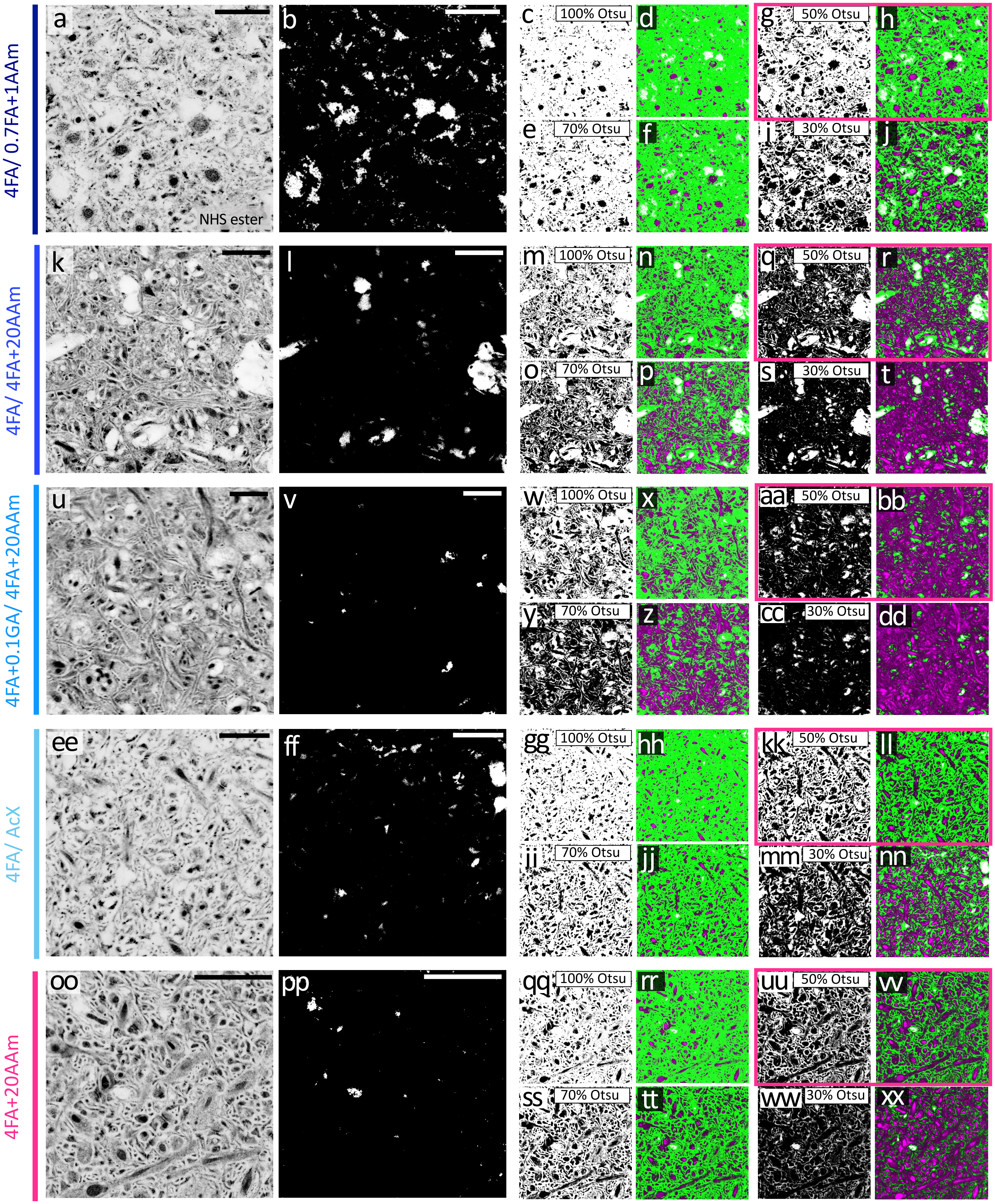


yy

| Ref in text | Fix/Post-fix | ECS + lipid membrane fraction (ECS+) | | | |
|---|---|---|---|---|---|
| | | 100% Otsu | 70% Otsu | 50% Otsu | 30% Otsu |
| *Fix-2* | 4FA /0.7FA+1AAm | 92%± 1% | 84%± 2% | 73%± 3% | 50%± 4% |
| *Fix-3* | 4FA /4FA+20AAm | 75%± 2% | 53%± 2% | 32%± 1% | 12%± 1% |
| *Fix-4* | 4FA+0.1GA /4FA+20AAm | 75%± 5% | 50%± 9% | 29%± 9% | 10%± 6% |
| *Fix-5* | 4FA /AcX | 89%± 2% | 78%± 3% | 64%± 4% | 36%± 4% |
| *Fix-6* | 4FA+20AAm | 79%± 3% | 58%± 5% | 38%± 5% | 14%± 5% |

**Supplementary Figure 13: Effect of fixation and post-fixation on extracellular space (ECS) preservation in expanded mouse brain tissue sections.** **a**, **k**, **u**, **ee**, **oo**, NHS ester pan-stained neuropil-rich regions in brain tissue fixed with the different indicated fixation schemes. **b**, **l**, **v**, **ff**, **pp**, same areas as in **a**, **k**, **u**, **ee**, and **oo** showing pixels (white) that are excluded from the analysis ($A_{excl.}$). **c**, **m**, **w**, **gg**, **qq**, same areas as in **a**, **k**, **u**, **ee**, and **oo**, respectively, where black pixels represent the areas selected by applying 100% of the Otsu threshold. Here, white pixels correspond to the uncorrected ECS + lipid membrane fraction ($A_{ECS}$). **d**, **n**, **x**, **hh**, **rr**, overlay of **a**, **k**, **u**, **ee**, **oo** (magenta), **b**, **l**, **v**, **ff**, **pp** (white), and **c**, **m**, **w**, **gg**, **qq** (green), respectively. **e**, **o**, **y**, **ii**, **ss** same areas as in **a**, **k**, **u**, **ee**, **oo**, respectively, where black pixels represent the areas selected by applying 70% of the Otsu threshold. **f**, **p**, **z**, **jj**, **tt** overlay of **a**, **k**, **u**, **ee**, **oo** (magenta), **b**, **l**, **v**, **ff**, **pp** (white), and **e**, **o**, **y**, **ii**, **ss** (green), respectively. **g**, **q**, **aa**, **kk**, **uu**, same areas as in **a**, **k**, **u**, **ee**, and **oo**, respectively, where black pixels represent the areas selected by applying 50% of the Otsu threshold. **h**, **r**, **bb**, **ll**, **vv**, overlay of **a**, **k**, **u**, **ee**, **oo** (magenta), **b**, **l**, **v**, **ff**, **pp** (white), and **g**, **q**, **aa**, **kk**, **uu** (green), respectively. **i**, **s**, **cc**, **mm**, **ww**, same areas as in **a**, **k**, **u**, **ee**, and **oo**, respectively, where black pixels represent the areas selected by applying 30% of the Otsu threshold. **j**, **t**, **dd**, **nn**, **xx**, overlay of **a**, **k**, **u**, **ee**, **oo** (magenta), **b**, **l**, **v**, **ff**, **pp** (white), and **i**, **s**, **cc**, **mm**, **ww** (green), respectively. **yy**, table summarizing the corrected ECS + lipid membrane (ECS+) values obtained by dividing $A_{ECS}$ by ($A_{total}$-$A_{excl.}$), where $A_{total}$ is the total field of view area (Fix-2: n = 8 FOVs from N = 1 independent experiment; Fix-3: n = 8 FOVs from N = 1 independent experiment; Fix-4: n = 8 FOVs from N = 1 independent experiment; Fix-5: n = 8 FOVs from N = 1 independent experiment; Fix-6: n = 14 FOVs from N = 3 independent experiments). Values in pink report the chosen ECS+ values for this comparison. Scale bars are corrected for the expansion factor (**a**, **b**, **k**, **l**, **u**, **v**, **ee**, **ff**, **oo**, **pp**) 3 µm.

SUPP FIG. 14

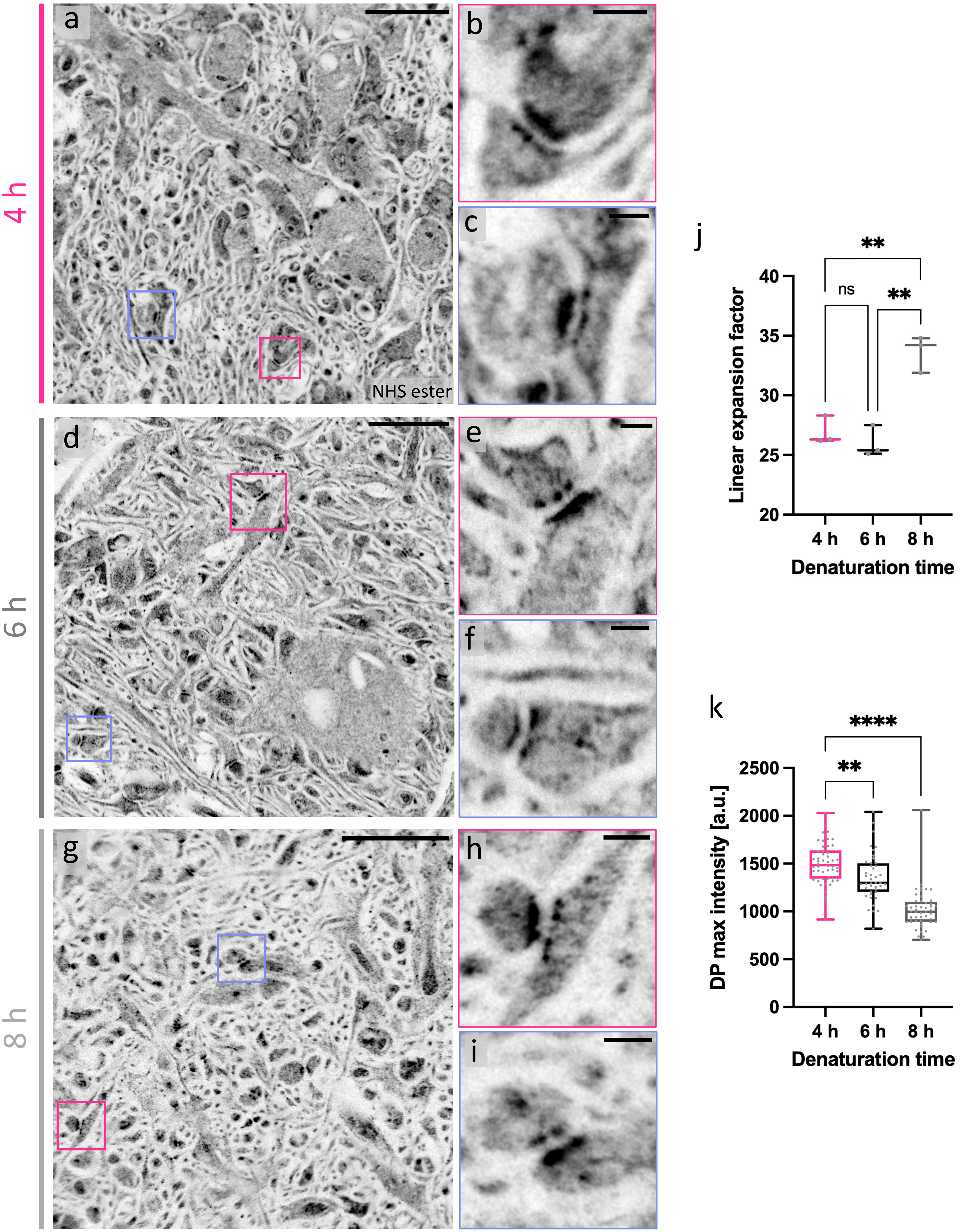
4 h
6 h
8 h
a
b
c
NHS ester
d
e
f
g
h
i
j
Linear expansion factor
40
35
30
25
20
**
ns
**
4 h
6 h
8 h
Denaturation time
k
DP max intensity [a.u.]
2500
2000
1500
1000
500
0
****
**
4 h
6 h
8 h
Denaturation time

**Supplementary Figure 14: Effect of denaturation time on expansion factor and protein retention in expanded mouse brain tissue sections.** **a**, NHS ester pan-stained brain tissue denatured for 4 h. **b**, magnified view of the area outlined in the pink box in **a**. **c**, magnified view of the area outlined in the lavender box in **a**. **d**, NHS ester pan-stained brain tissue denatured for 6 h. **b**, magnified view of the area outlined in the pink box in **d**. **e**, magnified view of the area outlined in the lavender box in **d**. **g**, NHS ester pan-stained brain tissue denatured for 8 h. **h**, magnified view of the area outlined in the pink box in **g**. **i**, magnified view of the area outlined in the lavender box in **g**. **j,** plot of expansion factors (4 h: n = 50 measurements from 3 FOVs; 6 h: n = 42 measurements from 3 FOVs; 8 h: n = 100 from 3 FOVs). **k**, relative protein retention was measured by comparing the peak intensity of dense projection (DP) NHS ester signal (4 h: n = 50 measurements from 3 FOVs; 6 h: n = 42 measurements from 3 FOVs; 8 h: 48 measurements from 3 FOVs). Images were processed with a gamma value ɣ=0.8. Black scale bars are corrected for the expansion factor (**a**, **d**, **g**) 2 µm, (**c**, **d**, **e**, **f**, **h**, **i**) 200 nm.

SUPP FIG. 15

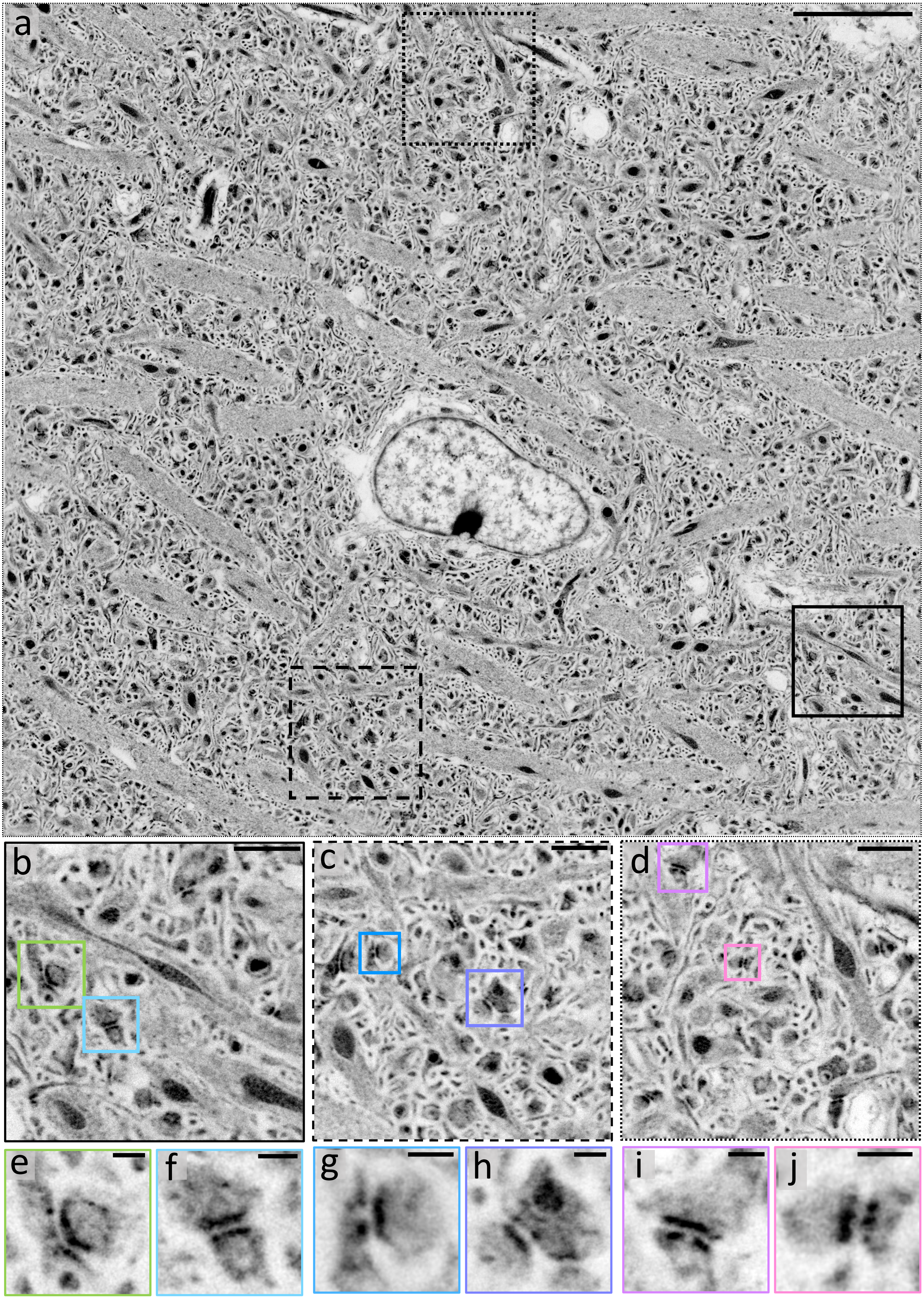

**Supplementary Figure 15: pan-ExM-t reveals mouse brain tissue ultrastructure (overview 1).** **a**, NHS ester pan-stained tissue section in the mouse hippocampus. **b-d**, magnified areas in the solid, dashed, and dotted black boxes in **a**, respectively, showing neuropil. **e**, **f**, magnified areas in the lime and light blue boxes in **b**, respectively, showing synapses. **g**, **h**, magnified areas in the blue and orchid boxes in **c**, respectively, showing synapses. **i**, **j**, magnified areas in the lavender and pink boxes in **d**, respectively, showing synapses. Images were acquired with a 25X/0.95 NA water objective. Images were processed with a gamma value ɣ=0.8. Black scale bars are corrected for the expansion factor (**a**) 5 µm, (**b-d**) 1 µm, (**e-j**) 200 nm.

SUPP FIG. 16

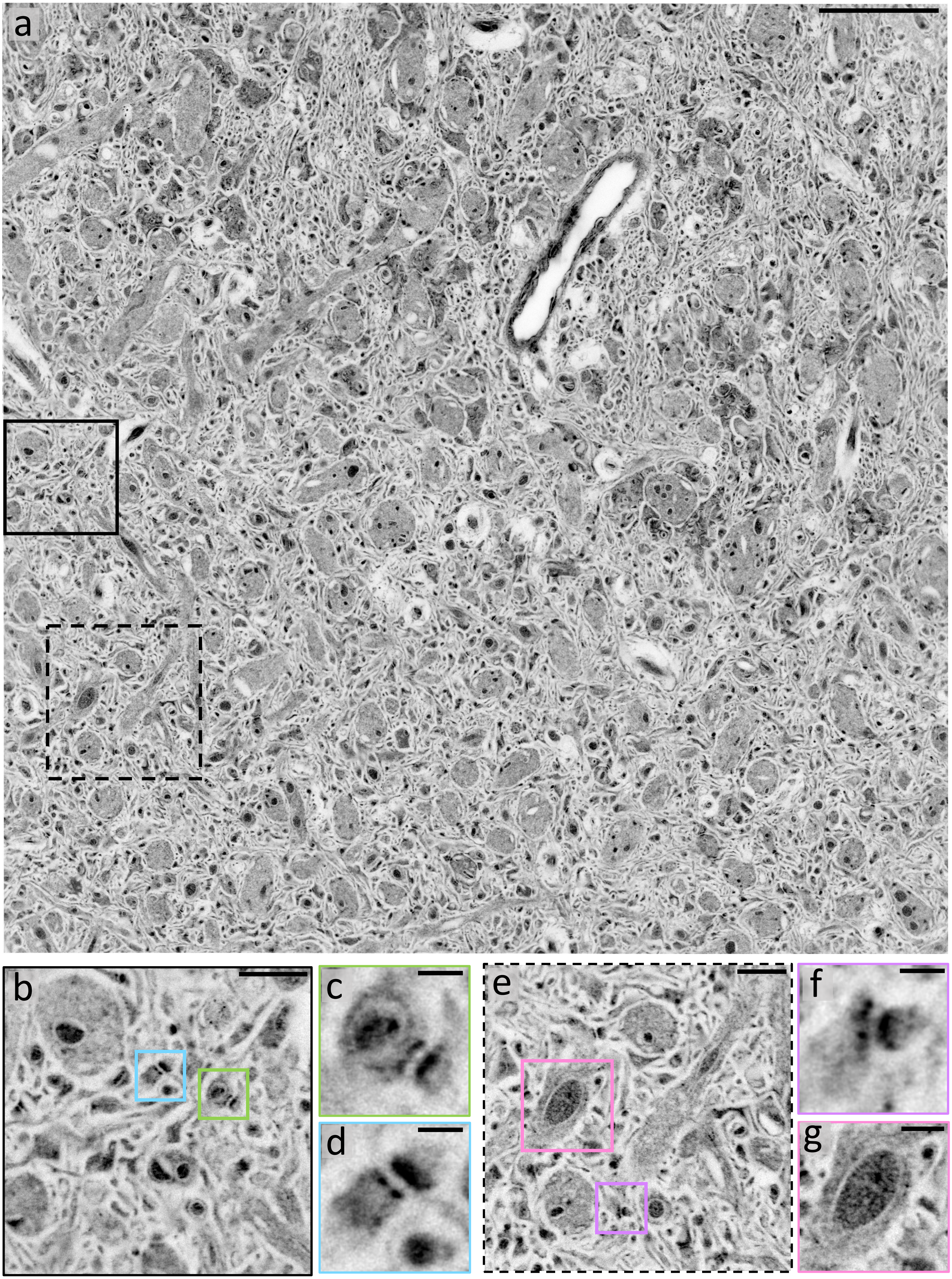

**Supplementary Figure 16: pan-ExM-t reveals mouse brain tissue ultrastructure (overview 2).** **a**, NHS ester pan-stained tissue section in the mouse cortex. **b**, **e**, magnified areas in the solid and dashed black boxes in **a**, respectively, showing neuropil. **c**, **d**, magnified areas in the lime and light blue boxes in **b**, respectively, showing synapses. **f**, **g**, magnified areas in the lavender and pink boxes in **e** showing a synapse and mitochondrion, respectively. Images were acquired with a 25X/0.95 NA water objective. Images were processed with a gamma value ɣ=0.7. Black scale bars are corrected for the expansion factor (**a**) 5 µm, (**b**, **e**) 1 µm, (**c**, **d**, **f**) 200 nm, (**g**) 500 nm.

SUPP FIG. 17

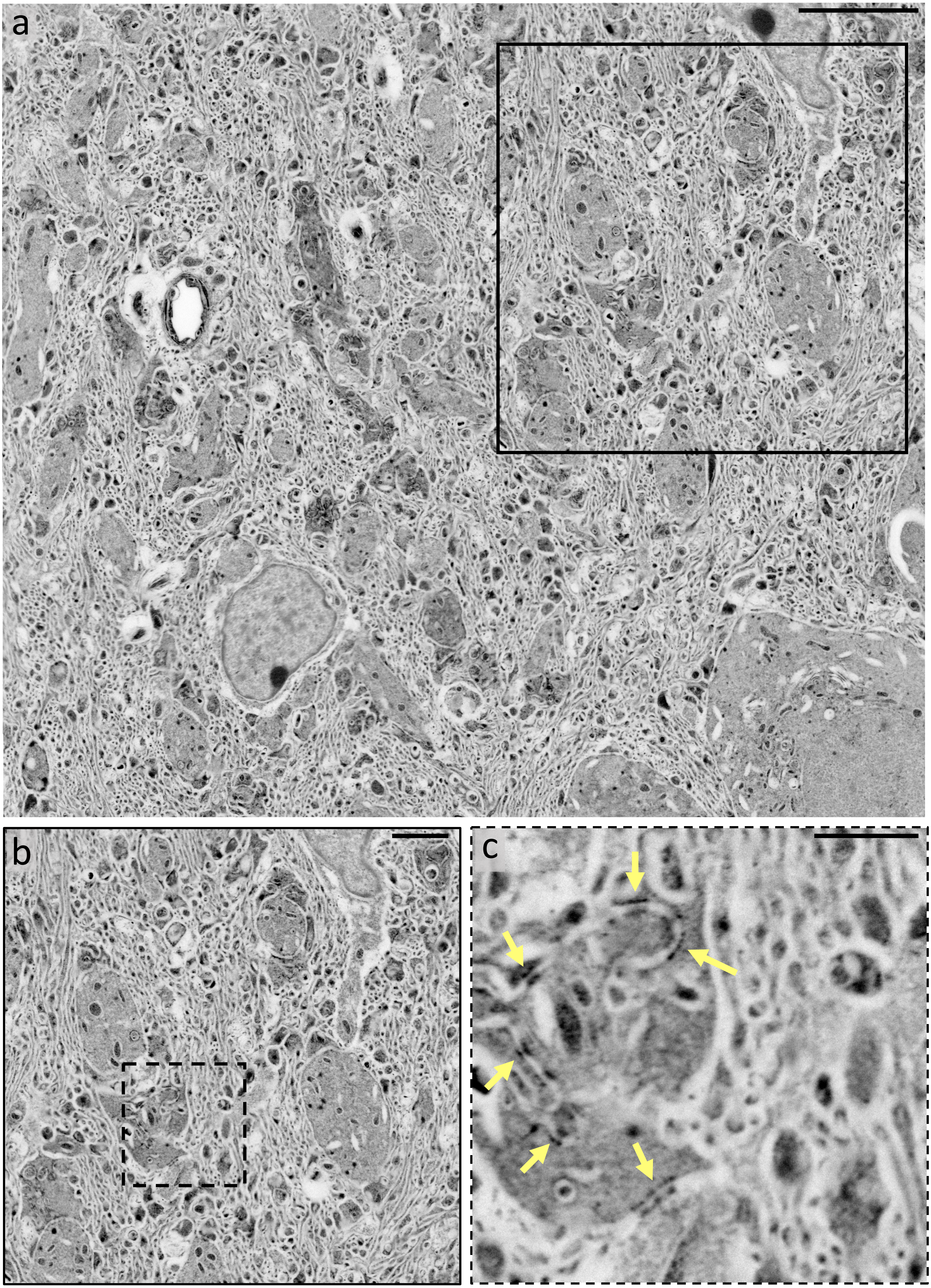

**Supplementary Figure 17: pan-ExM-t reveals mouse brain tissue ultrastructure (overview 3).** **a**, NHS ester pan-stained tissue section in the mouse cortex. **b**, magnified area in the black boxes in **a** showing neuropil. **c**, magnified area in the dotted black box in **b** showing multiple putative synapses (yellow arrows). Images were acquired with a 25X/0.95 NA water objective. Images were processed with a gamma value ɣ=0.7. Black scale bars are corrected for the expansion factor (**a**) 5 µm, (**b**) 2 µm, (**c**) 1 µm.

SUPP FIG. 18

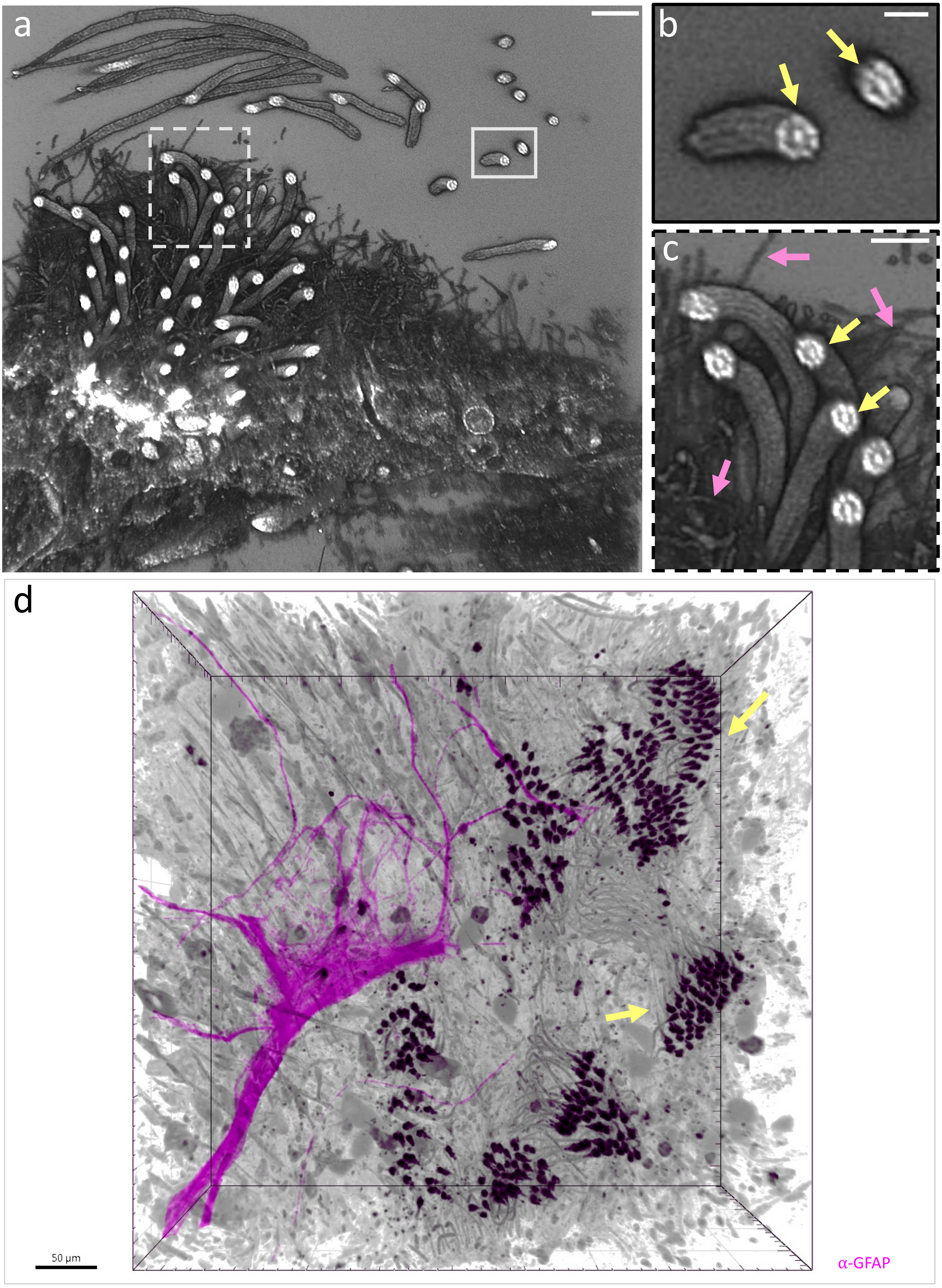

**Supplementary Figure 18: Multiciliated ependymal epithelia lining the lateral ventricles of the mouse brain**. **a**, 3D rendering of NHS ester pan-stained ependymal epithelia showing multiple cilia. **b**, **c**, magnified areas in the solid and dashed gray boxes in **a**, respectively, showing cilia with visible nine-fold microtubule symmetry (yellow arrows) and filopodia-like structures (pink arrows). **d**, 3D rendering of NHS ester pan-stained multi-ciliated ependymal epithelia (grayscale) and anti-GFAP immunolabeling (magenta). Yellow arrows point to cilia. 3D image processing details are described in the **Methods** section. Note that sample **a** was processed with *Fix-2* (4% FA fix and 0.7% FA + 1% AAm postfix). Image **a** was acquired with a 63X/1.2 NA water objective and image **d** was acquired with a 25X/0.95 NA water objective. Black scale bars are corrected for the expansion factor, (**a**, **d**) 1 µm, (**b**) 200 nm, (**c**) 400 nm.

SUPP FIG. 19

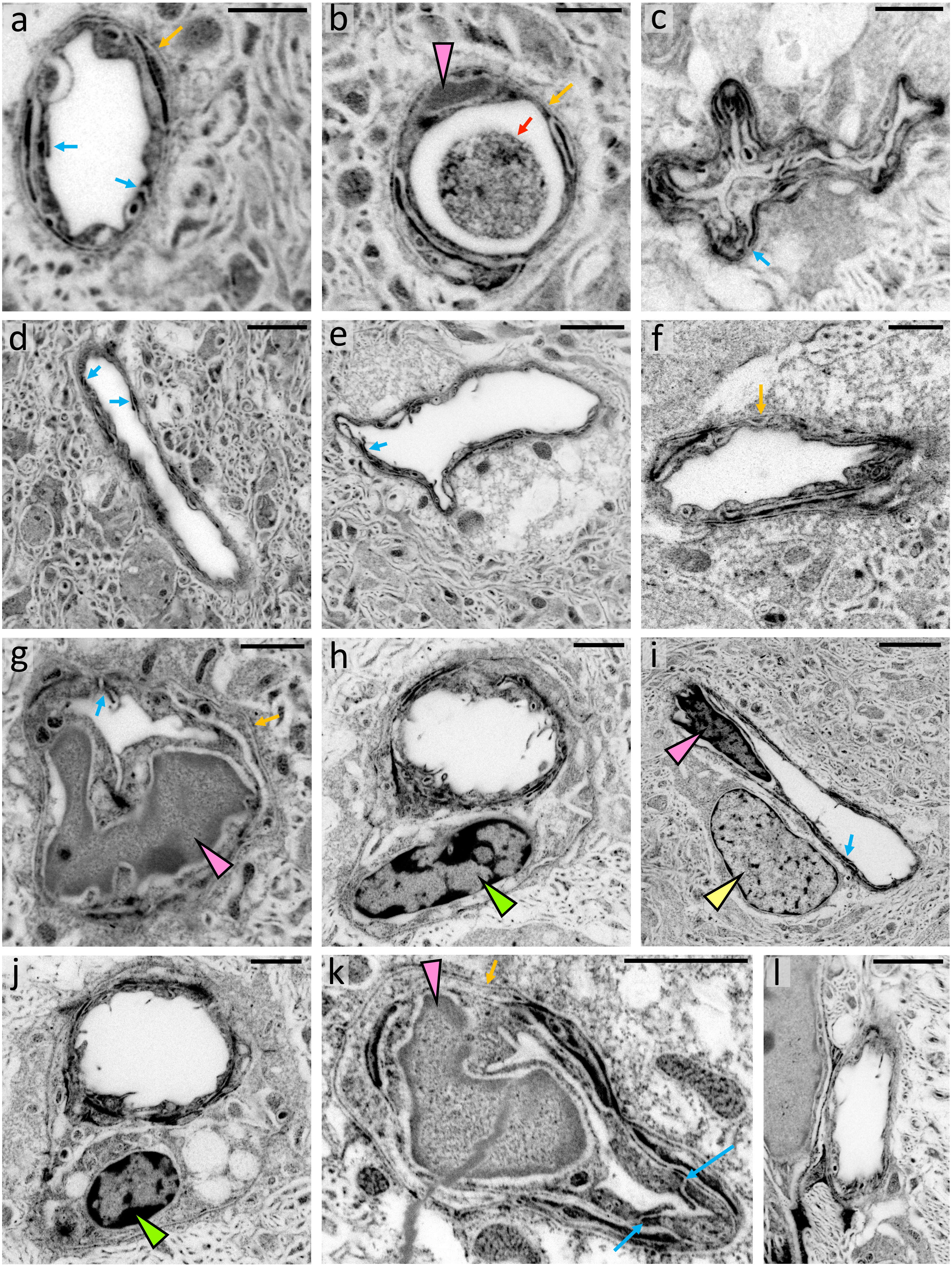

**Supplementary Figure 19: Gallery of mouse brain capillaries**. **a-l**, NHS ester pan-stained brain capillaries. Blue arrows point to tight junctions. Red arrow in **b** points to an erythrocyte. Tangerine arrow points to the basement membrane. Pink arrowheads in **b**, **g**, and **k** point to epithelial cell nuclei. Lime arrowheads in **h** and **j** point to pericyte nuclei. Yellow arrowhead in **i** points to a perivascular glial cell. All images were acquired with a 20X/0.95 NA water objective except for **f** and **k** which were acquired with a 86X/1.2 NA water objective. Images **a**, **b**, **c**, **d**, **e**, **g**, **i**, **j**, **k**, **l** were processed with a gamma value ɣ=0.7 and image **f** with ɣ=0.5. Black scale bars are corrected for the expansion factor, (**a**, **b**, **c**, **f**, **g**, **j**, **k**, **l**) 1 µm, (**d**, **e**, **i**) 2 µm.

### SUPP FIG. 20

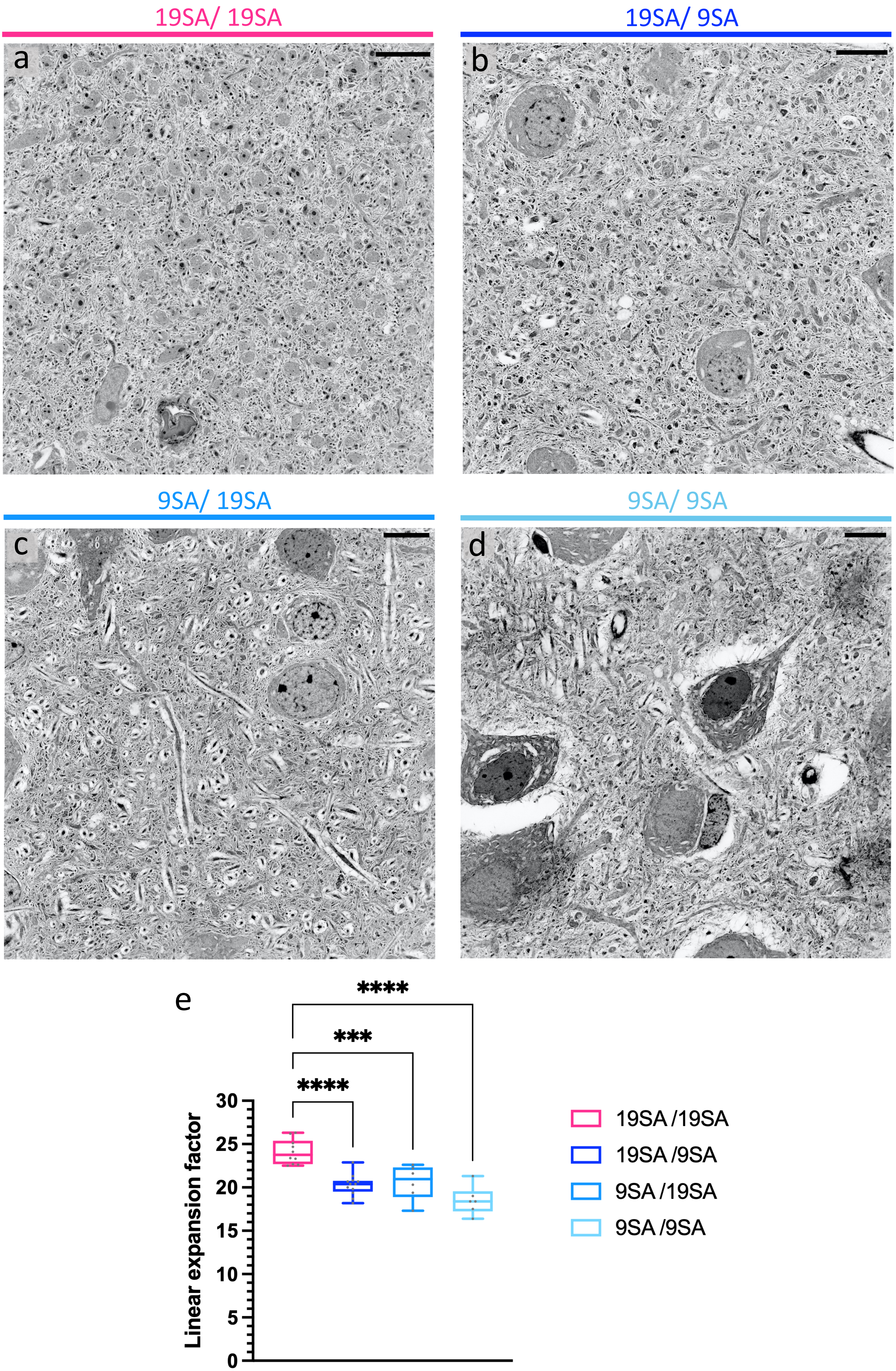
19SA/ 19SA
19SA/ 9SA
9SA/ 19SA
9SA/ 9SA
a
b
c
d
e
****
***
****
Linear expansion factor
0
5
10
15
20
25
30
19SA /19SA
19SA /9SA
9SA /19SA
9SA /9SA

**Supplementary Figure 20: Effect of sodium acrylate (SA) monomer concentration on expanded mouse brain tissue integrity.** **a**, NHS ester pan-stained brain tissue gelled with 1st expansion gel monomer solution containing 19% (w/v) SA and 2nd expansion gel monomer gel monomer solution containing 19% (w/v) SA. **b**, NHS ester pan-stained brain tissue gelled with 1st expansion gel monomer solution containing 19% (w/v) SA and 2nd expansion gel monomer gel monomer solution containing 9% (w/v) SA. **c**, NHS ester pan-stained brain tissue gelled with 1st expansion gel monomer solution containing 9% (w/v) SA and 2nd expansion gel monomer gel monomer solution containing 19% (w/v) SA. **d**, NHS ester pan-stained brain tissue gelled with 1st expansion gel monomer solution containing 9% (w/v) SA and 2nd expansion gel monomer gel monomer solution containing 9% (w/v) SA. **a-c** show neuropil and cell bodies with satisfactory preservation, while **d** shows detached cell bodies and distorted neuropil. **e**, Plot of expansion factors (19SA/ 19SA: n = 254 measurements from 10 FOVs from 3 independent experiments; 19SA/ 9SA: n = 267 measurements from 12 FOVs from 2 independent experiments; 9SA/ 19SA: n = 88 measurements from 6 FOVs from 2 independent experiments; 9SA/ 9SA: n = 86 measurements from 2 independent experiments). Images were processed with a gamma value ɣ=0.7. Black scale bars are corrected for the expansion factor (**a-d**) 5 µm.

SUPP FIG. 21

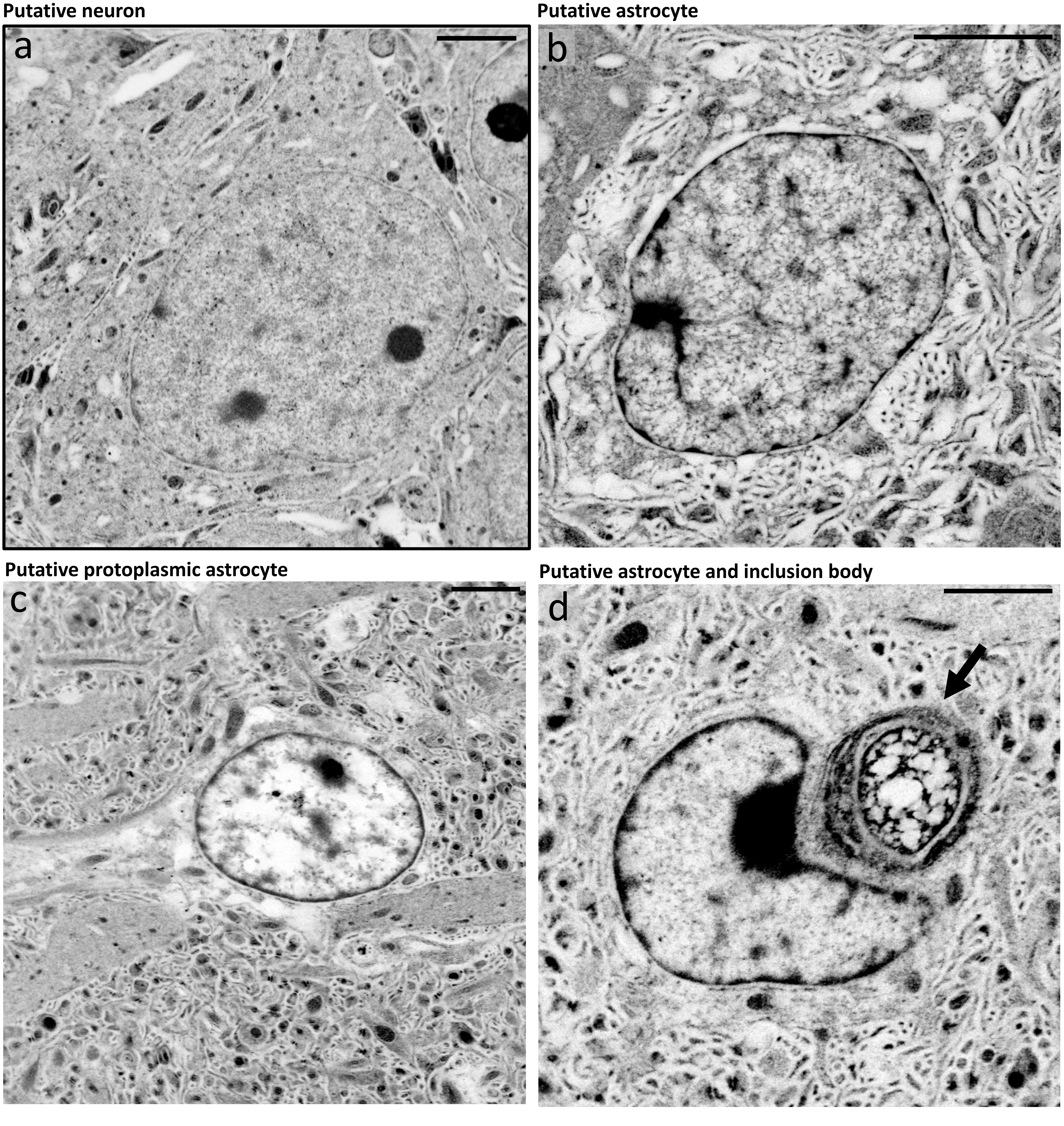


**Supplementary Figure 21: Astrocytes and neurons exhibit distinct NHS ester pan-staining patterns.** **a**, NHS ester pan-stained putative neuron showing a homogenous nuclear chromatin pattern. **b**, NHS ester pan-stained putative astrocyte showing a heterogeneous nuclear chromatin pattern and an irregular cytoplasm. **c**, NHS ester pan-stained putative protoplasmic astrocyte showing a pale nucleus with patches of chromatin, a pale cytoplasm, and triangular projections. **d**, NHS ester pan-stained putative astrocyte showing a heterogenous nuclear chromatin pattern, irregular cytoplasm, and an inclusion body (arrow). All images were processed with a gamma value of ɣ=0.7 Scale bars corrected for the expansion factor (**a-d**) 2 µm.

SUPP FIG. 22

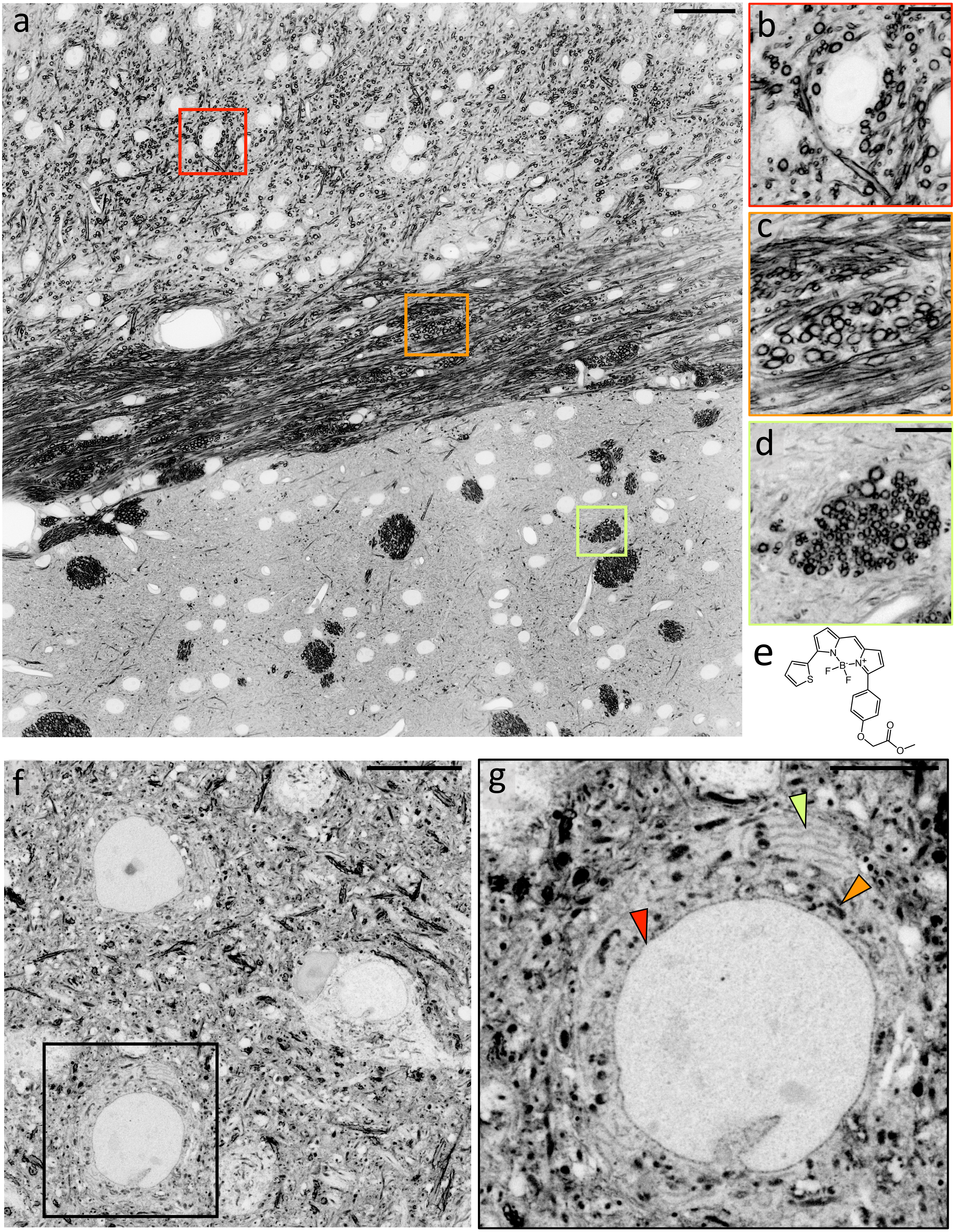

**Supplementary Figure 22: BODIPY-TR-Methyl Ester (BDP) pan-staining reveals lipophilic components in 4-fold expanded mouse brain tissue sections.** **a**, tiled image of a BDP pan-stained tissue section expanded ~4-fold. **b-d**, magnified images of the areas outlined by the red, orange, and lime boxes in **a**, showing highly pan-stained myelinated axons in top and lateral views. **e**, chemical structure of BODIPY-TR-Methyl Ester. **f**, BDP pan-stained tissue section expanded ~4-fold showing neurons. **g**, magnified image of the area outlined by the black box in **f** showing the nuclear envelope (red arrow), the endoplasmic reticulum (orange arrow), and mitochondria (lime arrow). Scale bars are not corrected for the expansion factor. (**a**) 100 µm, (**b-d**) 20 µm, (**f**) 50 µm, (**g**) 20 µm.

### SUPP FIG. 23

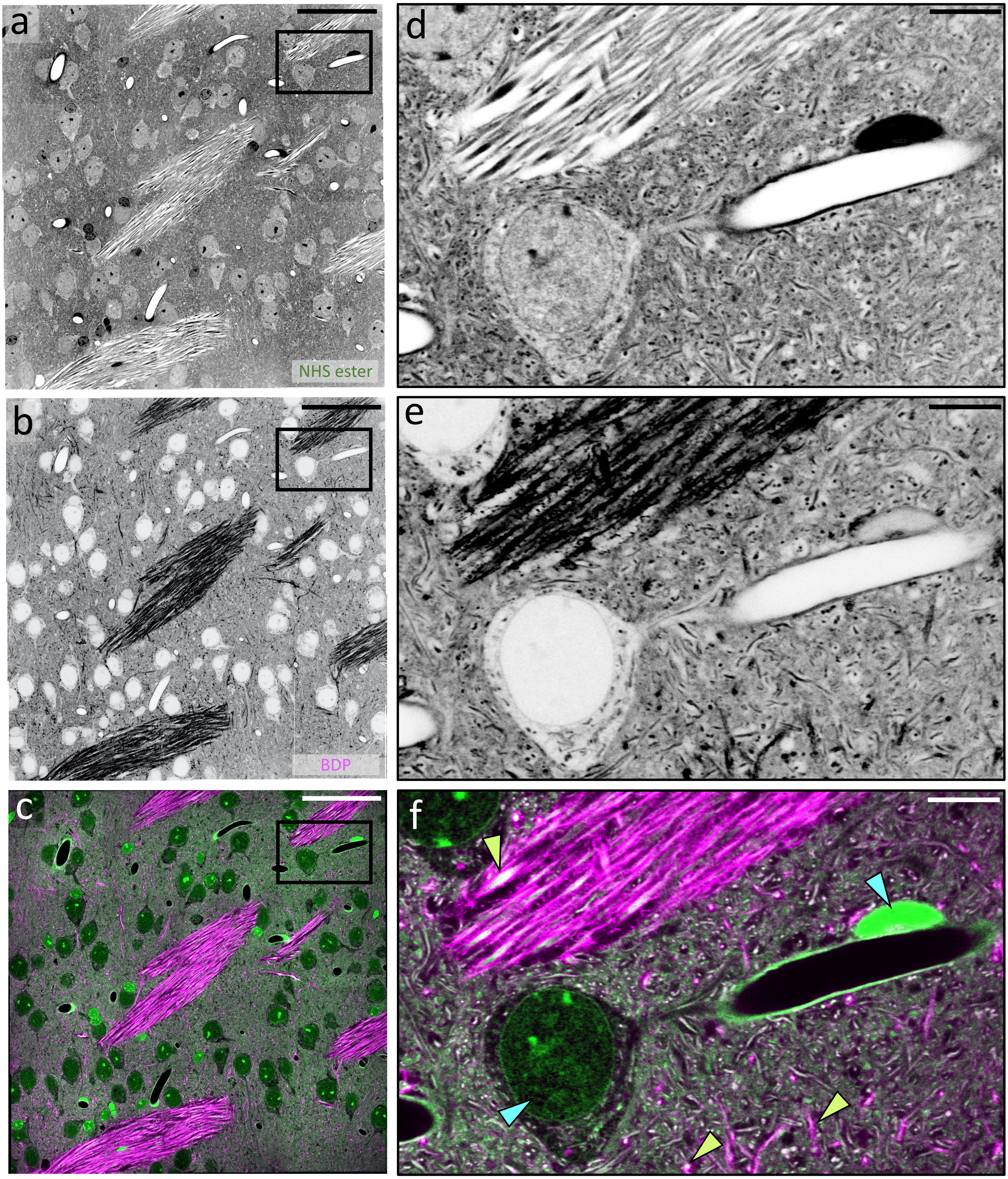


**Supplementary Figure 23: BODIPY-TR-Methyl Ester (BDP) and hydrophilic NHS ester differential pan-staining in a 4-fold expanded mouse brain tissue section. a**, tiled image of an NHS ester pan-stained tissue section expanded ~4-fold. **b**, BDP pan-staining corresponding to area shown in **a**. **c**, overlay of **a** and **b**. **d-f**, magnified images of the areas outlined by the black boxes in **a-c**. Lime arrows point to myelin in axons, cyan arrows point to nuclei stained exclusively by NHS ester. Scale bars are not corrected for the expansion factor. (**a-c**) 200 µm, (**d-f**) 20 µm.

### SUPP FIG. 24

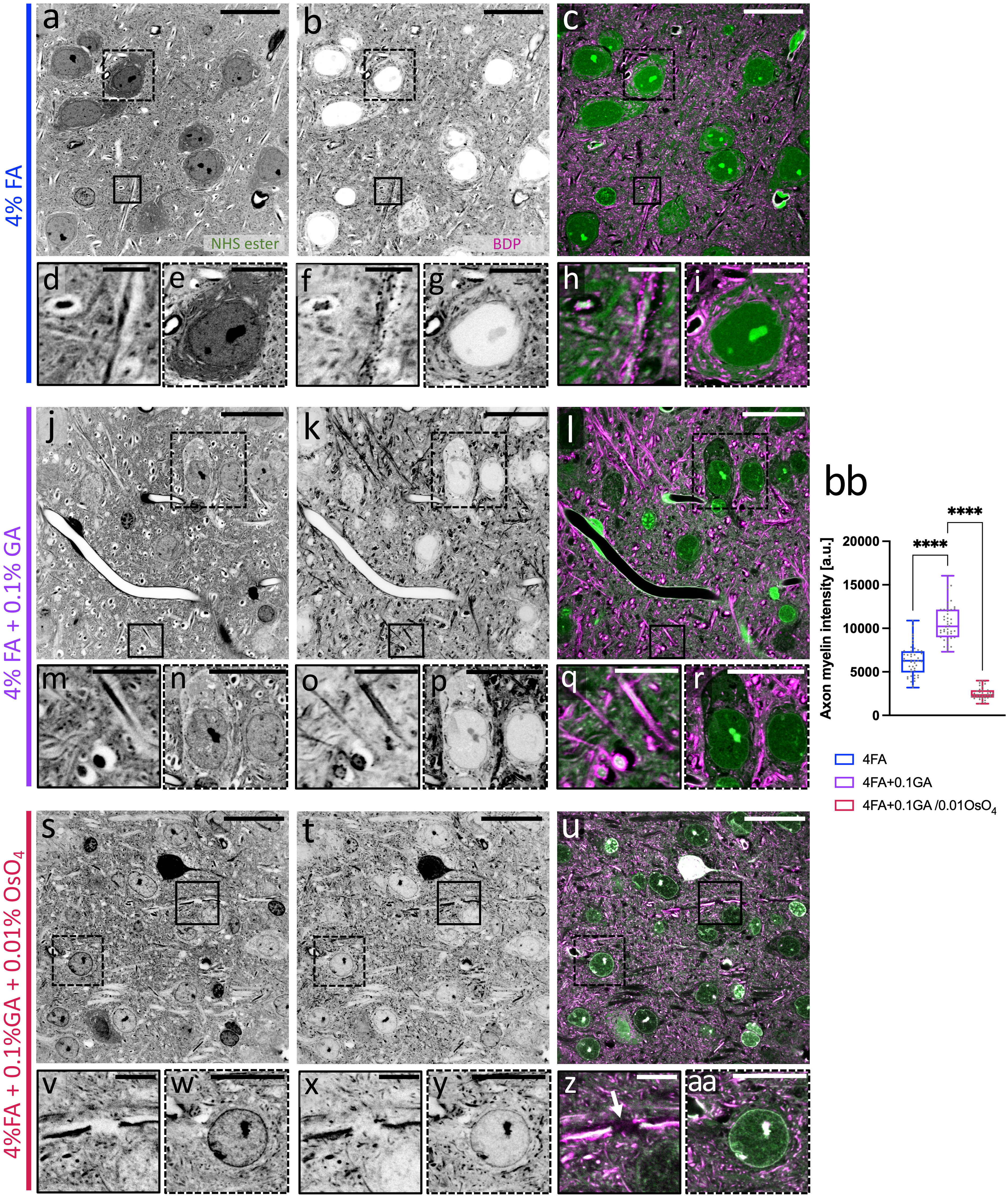

4% FA
4% FA + 0.1% GA
4%FA + 0.1%GA + 0.01% OsO4
a
b
c
d
e
f
g
h
i
j
k
l
m
n
o
p
q
r
s
t
u
v
w
x
y
z
aa
bb
NHS ester
BDP
****
****
Axon myelin intensity [a.u.]
20000
15000
10000
5000
0
4FA
4FA+0.1GA
4FA+0.1GA /0.01OsO4

**Supplementary Figure 24: Effect of fixation on BODIPY-TR-Methyl Ester (BDP) pan-staining in 4-fold expanded mouse brain tissue sections.** **a**, NHS ester pan-stained tissue section expanded ~4-fold fixed with 4% FA. **b**, BDP pan-staining of the same area shown in **a**. **c**, overlay. **d**, **f**, **h**, magnified images of the areas outlined by the black boxes in **a**, **b**, and **c**, respectively, showing myelinated axons. **e**, **g**, **i**, magnified images of the areas outlined by the dashed black boxes in **a**, **b**, and **c**, respectively, showing a cell body. **j**, NHS ester pan-stained tissue section expanded ~4-fold fixed with 4% FA + 0.1% GA. **k**, BDP pan-staining of the same area shown in **j**. **l**, overlay. **m**, **o**, **q**, magnified images of the areas outlined by the black boxes in **j**, **k**, and **l**, respectively, showing myelinated axons. **n**, **p**, **r**, magnified images of the areas outlined by the dashed black boxes in **j**, **k**, and **l**, respectively, showing a cell body. **s**, NHS ester pan-stained tissue section expanded ~4-fold fixed with 4% FA. **t**, BDP pan-staining of the same area shown in **s**. **u**, overlay. **v**, **x**, **z**, magnified images of the areas outlined by the black boxes in **s**, **t**, and **u**, respectively, showing myelinated axons. **w**, **y**, **aa** magnified images of the areas outlined by the dashed black boxes in **s**, **t**, and **u**, respectively, showing a cell body. **bb**, plot comparing axon myelin BDP intensity across the indicated fixation conditions (4% FA: n = 46 measurements from 5 FOVs; 4% FA + 0.1% GA: n = 46 measurements from 5 FOVs; 4% FA + 0.1% GA + 0.01 % $OsO_4$: n = 38 measurements from 5 FOVs). ****: $p<0.0001$. Scale bars are not corrected for the expansion factor. (**a-c**, **j-l**, **s-u**), 100 µm, (**d**, **f**, **h**, **m**, **o**, **q**, **v**, **x**, **z**) 20 µm, (**e**, **g**, **h**, **n**, **p**, **r**, **w**, **y**, **aa**) 50 µm.

SUPP FIG. 25

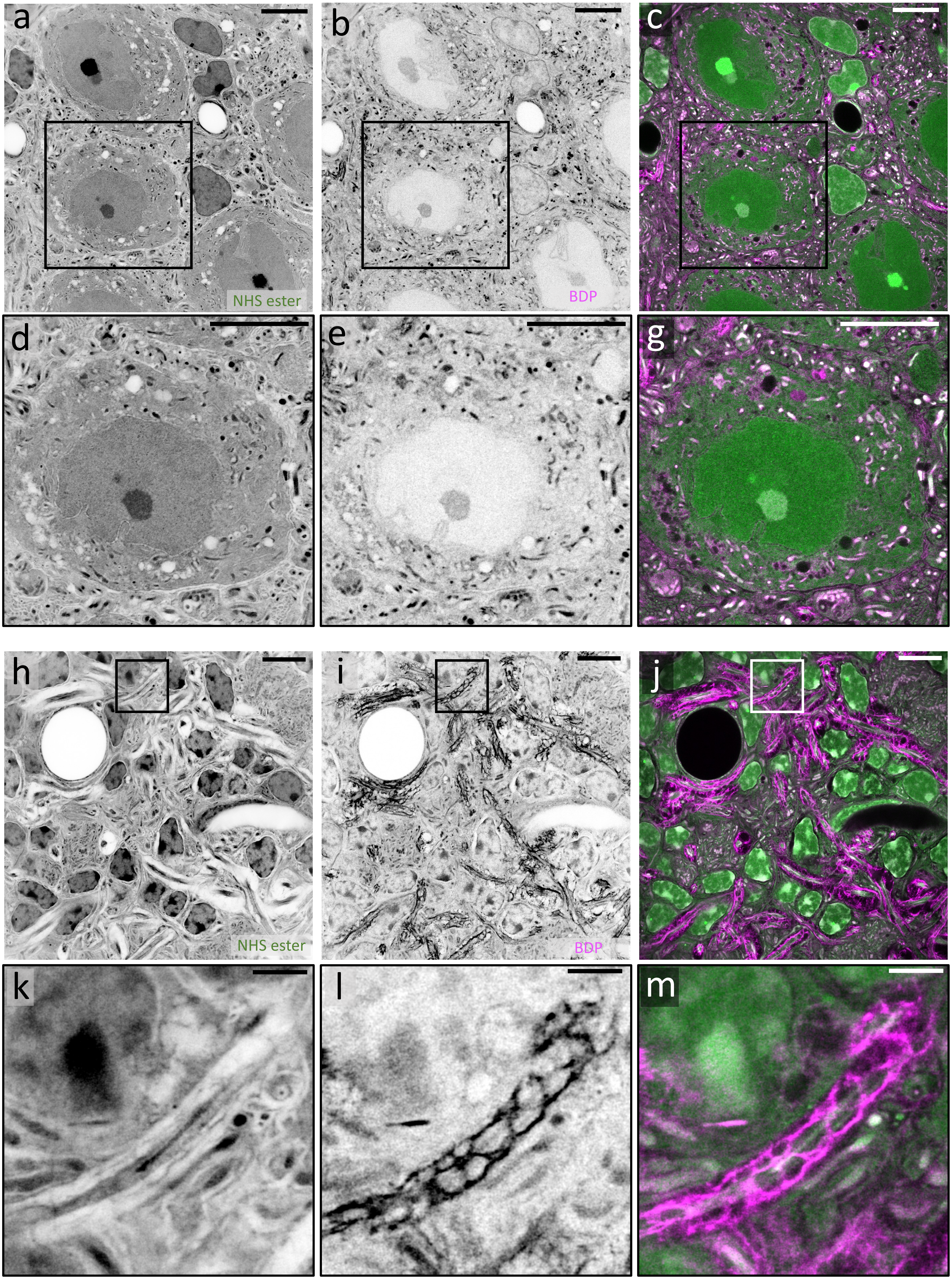

**Supplementary Figure 25: BODIPY-TR-Methyl Ester (BDP) and hydrophilic NHS ester differential pan-staining in pan-ExM-t expanded mouse brain tissue sections.** **a**, NHS ester pan-stained tissue section fixed with 4% FA + 0.1% GA and expanded 13-fold showing neuron somas. **b**, BDP pan-staining corresponding to area shown in **a**. **c**, overlay of **a** and **b**. **d-g,** magnified images of the areas outlined by the boxes in **a-c**. **h**, NHS ester pan-stained tissue section fixed with 4% FA + 0.1% GA and expanded 13-fold showing myelinated axons. **i**, BDP pan-staining corresponding to area shown in **h.** **j**, overlay of **h** and **i.** **k-m**, magnified images of the areas outlined by the boxes in **h-j**. Scale bars are corrected for the expansion factor. (**a-j**) 5 µm, (**k-m**) 1 µm.

### SUPP FIG. 26

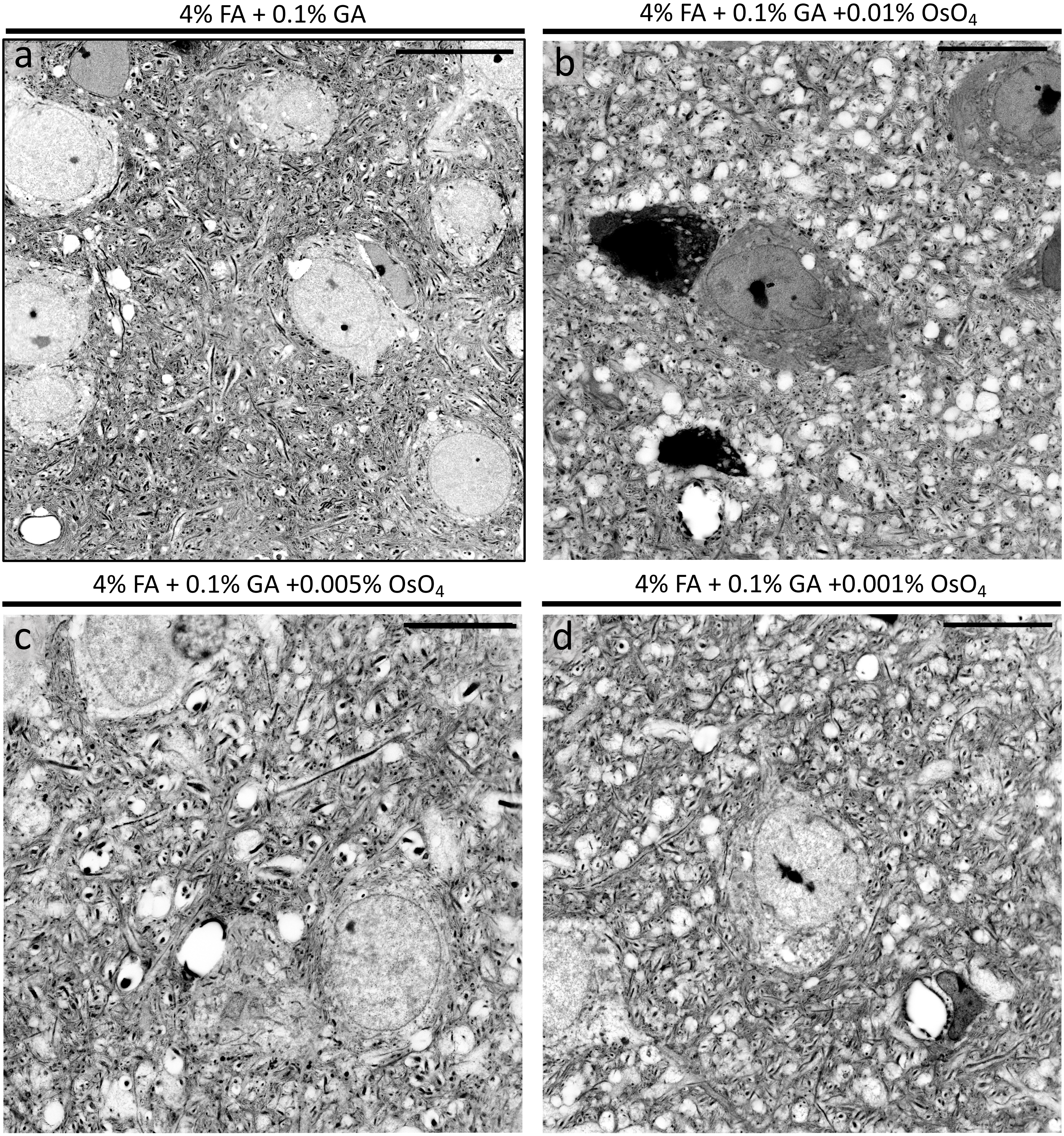


**Supplementary Figure 26: Effect of osmification on pan-ExM-t expanded brain tissue integrity.** **a**, NHS ester pan-stained brain tissue fixed with 4% FA + 0.1% GA. **b**, NHS ester pan-stained brain tissue fixed with 4% FA + 0.1% GA + 0.01% $OsO_4$. **c**, NHS ester pan-stained brain tissue fixed with 4% FA + 0.1% GA + 0.005% $OsO_4$. **d**, NHS ester pan-stained brain tissue fixed with 4% FA + 0.1% GA + 0.001% $OsO_4$. Very dark neuron perikarya (**a**) and multiple tissue perforations (**b-d**) appear to be artifactual. Images **b-d** were processed with a gamma value ɣ=0.7. Scale bars are not corrected for the expansion factor (**a-d**) 200 µm.

SUPP FIG. 27

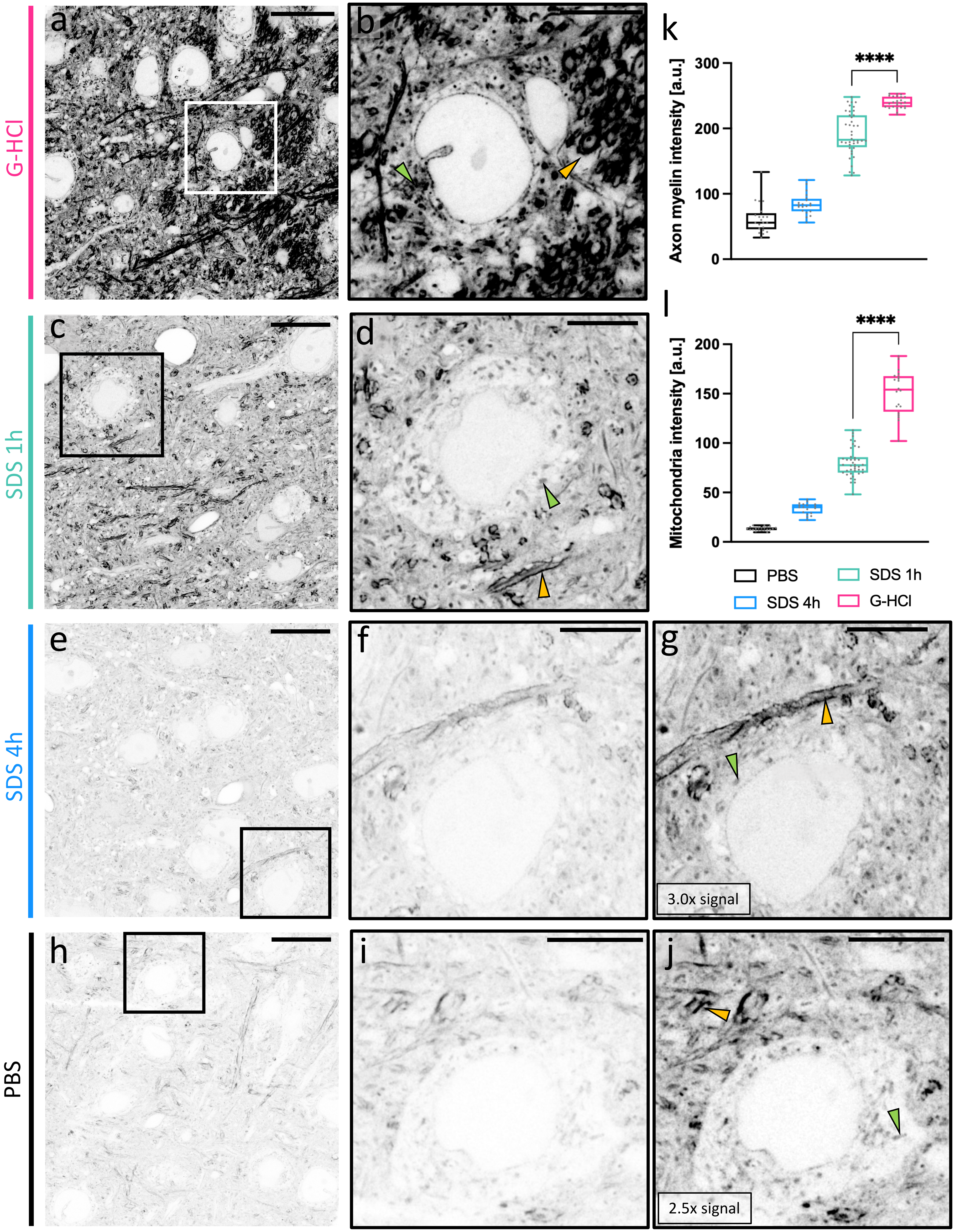

G-HCl
SDS 1h
SDS 4h
PBS
a
b
c
d
e
f
g
h
i
j
k
l
3.0x signal
2.5x signal
Axon myelin intensity [a.u.]
0
100
200
300
****
Mitochondria intensity [a.u.]
0
50
100
150
200
****
PBS
SDS 1h
SDS 4h
G-HCl

**Supplementary Figure 27: BODIPY-TR-Methyl Ester (BDP) signal is higher in G-HCl-denatured than in SDS- and non-denatured ~4-fold expanded mouse brain tissue sections. a,** BDP pan-stained tissue section denatured with G-HCl. **b**, magnified image of the area outlined by the black box in **a**. **c**, BDP pan-stained tissue section denatured with SDS for 1 h. **d**, magnified image of the area outlined by the black box in **c**. **e**, BDP pan-stained tissue section denatured with SDS for 4 h. **f**, magnified image of the area outlined by the black box in **e**. **g**, same area as in **f** with signal increased 2.5-fold. **e**, BDP pan-stained tissue section non-denatured. **i**, magnified image of the area outlined by the white box in **h**. **j**, Same area as in **I** with signal increased 3-fold. Samples in **a-j** were all expanded ~4-fold. Orange arrows point to myelinated axons. Green arrows point to mitochondria in neuron somas. **k**, comparison of BDP intensities in axons across different denaturation conditions (PBS: n = 41 measurements, N = 2 FOVs; SDS 4h: n = 20 measurements, N = 1 FOV; SDS 1h: n = 40 measurements, N = 4 FOVs; G-HCl: n = 24 measurements, N = 1 FOV). **l**, comparison of BDP intensities in mitochondria across different denaturation conditions (PBS: : n = 41 measurements, N = 2 FOVs; SDS 4h: n = 18 measurements, N = 1 FOV; SDS 1h: n = 40 measurements, N = 4 FOVs; G-HCl: n = 20 measurements, N = 1 FOV). Means ± standard deviations are reported. An ANOVA with post-hoc Tukey test was used to analyze the data. ****: $p<0.0001$. Scale bars are not corrected for the expansion factor. (**a**, **c**, **e**, **h**) 50 µm, (**b**, **d**, **f**, **g**, **i**, **j**) 20 µm .

### SUPP FIG. 28

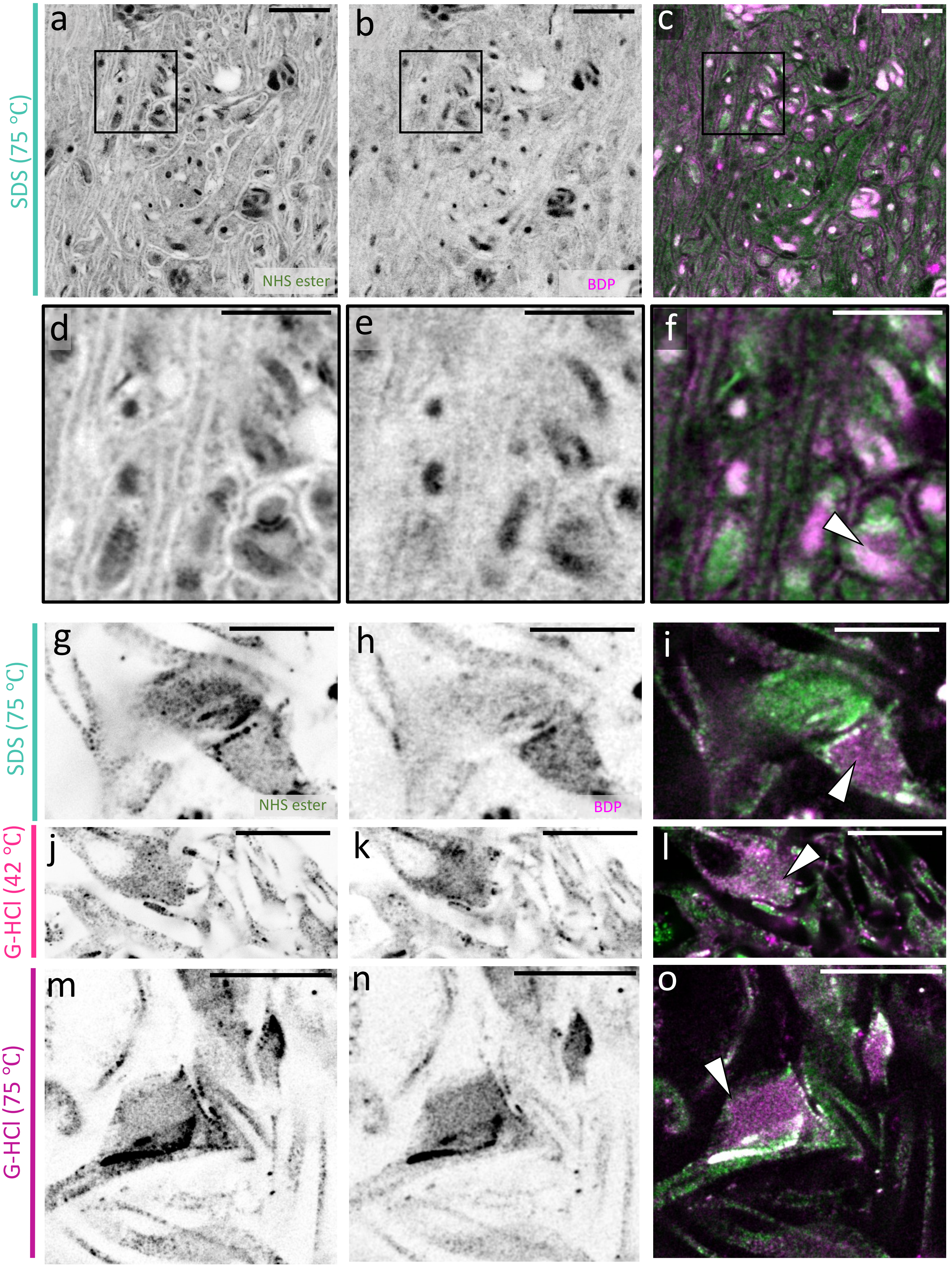

SDS (75 °C)
a
b
c
NHS ester
BDP
d
e
f
SDS (75 °C)
g
h
i
NHS ester
BDP
G-HCl (42 °C)
j
k
l
G-HCl (75 °C)
m
n
o

**Supplementary Figure 28: BODIPY-TR-Methyl Ester (BDP) does not label neurite boundaries specifically.** **a**, NHS ester pan-stained tissue section fixed with 4% FA + 0.1% GA and denatured with SDS. **b**, BDP pan-staining of the same area as in **a**. **c**, overlay of **a** and **b**. **d-f**, magnified images of the areas outlined by the black boxes in **a-c** showing BDP pan-stained mitochondria, including a mitochondrion in a synapse (white arrow). **g**, NHS ester pan-stained synapse in a dissociated neuron sample fixed with 3% FA + 0.1% GA and denatured with SDS at 75℃. **h**, BDP pan-staining of the same area as in **g**. **i**, overlay of **g** and **h**. **j**, NHS ester pan-stained synapse in a dissociated neuron sample fixed with 3% FA + 0.1% GA and denatured with guanidium hydrochloride (G-HCl) at 42℃. **k**, BDP pan-staining of the same area as in **j**. **l**, overlay of **j** and **k**. **m-o**, same as **j-l** but in a dissociated neuron sample denatured with G-HCl at 75℃. White arrows in **i**, **l**, and **o** point at the presynaptic bouton which is differentially pan-stained with BDP. It is worth noting that BDP does not appear to stain cellular membranes at the boundary of neurites regardless of whether the detergent SDS is used. BDP however appears to stain hydrophobic compartments such as mitochondria and the presynaptic compartment (likely because of the abundance of synaptic vesicles). Scale bars are not corrected for the expansion factor. (**a-c**) 20 µm, (**d-f**) 10 µm, (**g-o**) 20 µm.

SUPP FIG. 29

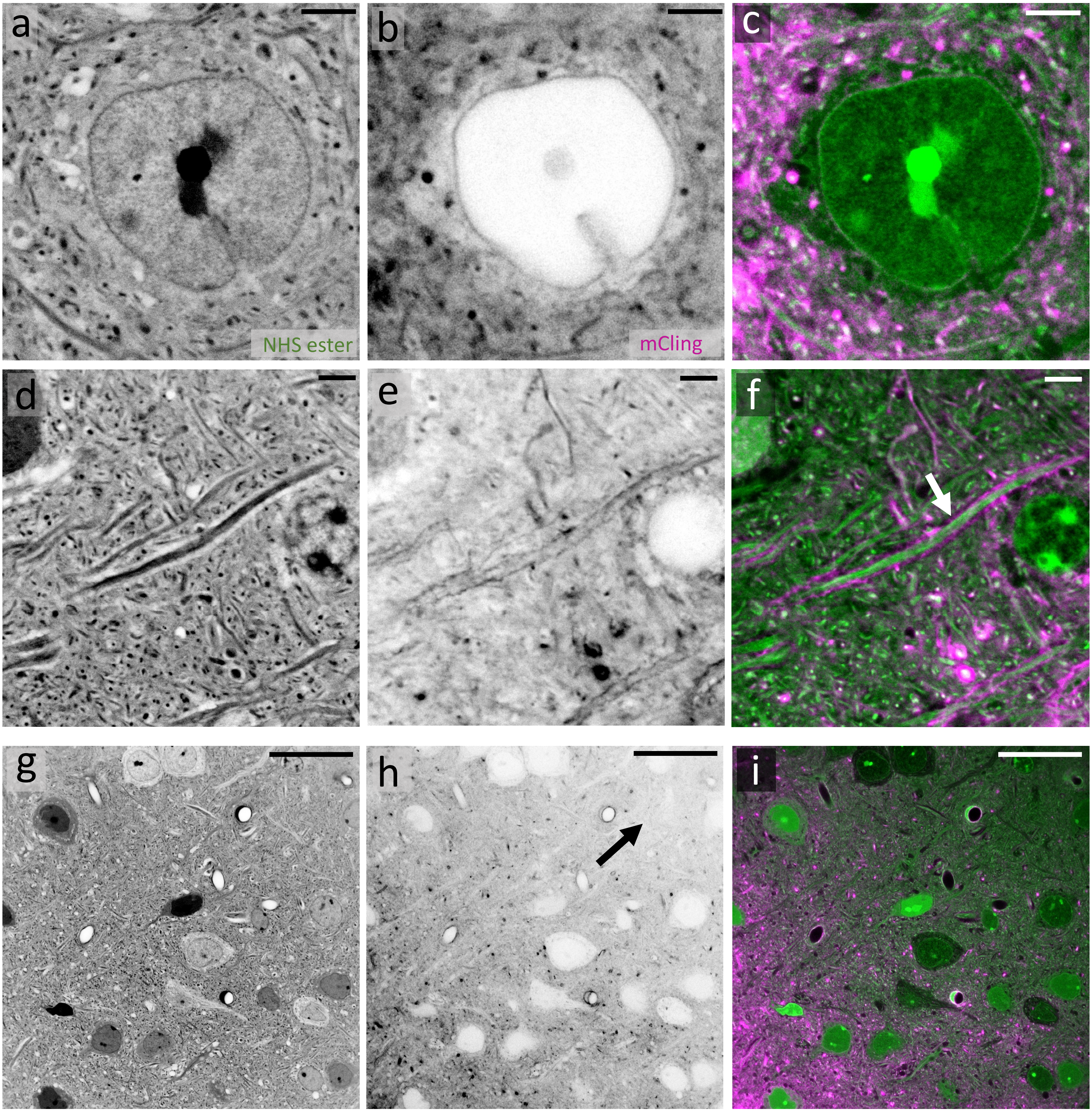


**Supplementary Figure 29: mCling only labels the surface of 4-fold expanded mouse brain tissue sections fixed with 3% FA + 0.1% GA.** **a**, NHS ester pan-stained tissue section expanded ~4-fold showing a neuron soma. **b**, mCling staining of the same area shown in **a**. **c**, overlay. **d**, NHS ester pan-stained tissue section expanded ~4-fold showing a myelinated axon. **e**, mCling staining of the same area shown in **d**. **f**, overlay. White arrow points to myelinated axon. **g**, Large FOV image of an NHS ester pan-stained tissue section expanded ~4-fold. **h**, mCling pan-staining of the same area shown in **a** showing uneven labeling which is indicative of poor ligand penetration. Black arrow points to region of low mCling staining. **i**, overlay. Scale bars are not corrected for the expansion factor. (**a-f**), 10 µm, (**g-i**) 100 µm.

### SUPP FIG. 30

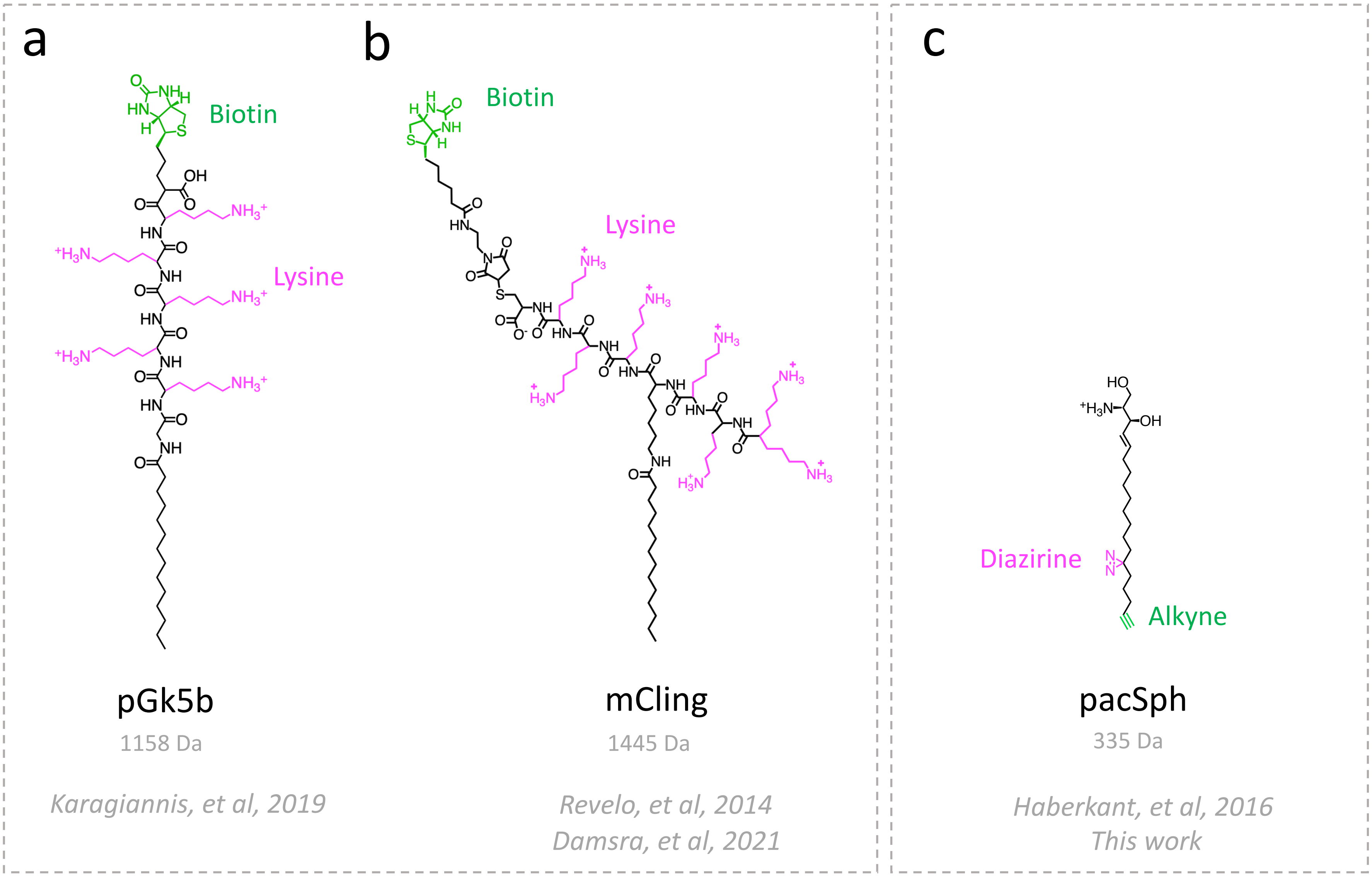


**Supplementary Figure 30: Chemical structures of pGk5b, mCling, and pacSph. a**, chemical structure of palmitoyl-glycine-(penta)lysine-biotin (pGk5b). pGk5b is 1158 Da in size. It has 5 reactive primary amine groups, a palmitoyl group for lipid membrane intercalation, a glycine for mechanical flexibility, and a biotin reporter group. **b**, chemical structure of membrane-binding fluorophore-cysteine-(hepta)lysine-palmitoyl group (mCling). mCling (biotin) is 1445 Da in size. It has 7 reactive primary amine groups, a palmitoyl group for lipid membrane intercalation, and a biotin (or fluorophore) reporter group. **c**, chemical structure of photoactivatable and clickable sphingosine (pacSph). pacSph is 335 Da in size. It is a sphingosine that intercalates lipid membranes. It has a diazirine UV photocrosslinkable group and an alkyne reporter group.

### SUPP FIG. 31

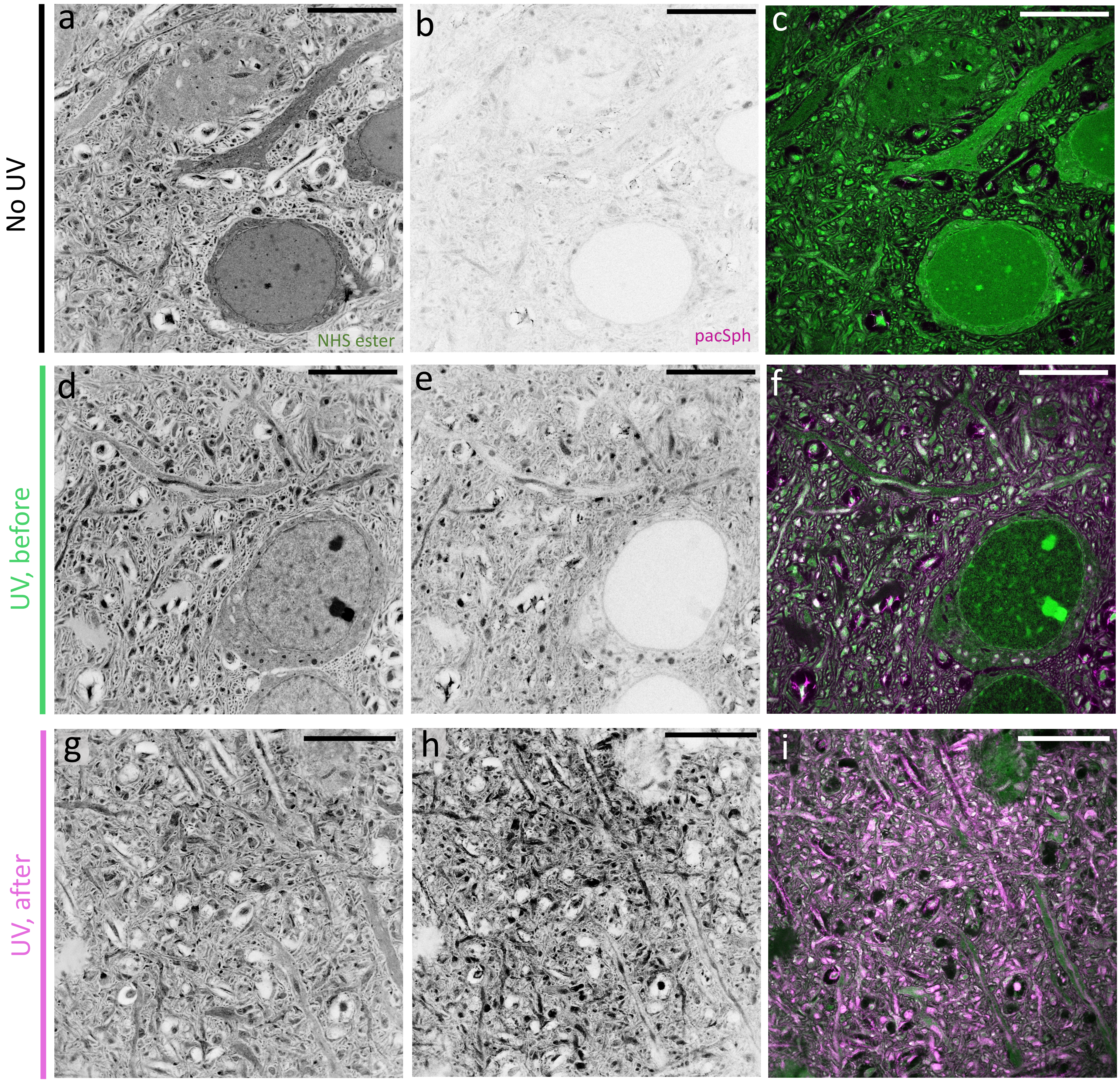


**Supplementary Figure 31: Effect of UV irradiation on pacSph pan-staining in pan-ExM-t expanded mouse brain tissue sections.** **a**, NHS ester pan-stained neuropil. **b**, pacSph pan-staining of the same area shown in **a** without UV irradiation. **c**, overlay. **d**, NHS ester pan-stained neuropil. **e**, pacSph pan-staining of the same area shown in **d** with UV irradiation before hydrogel embedding. **f**, **o**verlay. **g**, NHS ester pan-stained neuropil. **h**, pacSph pan-staining of the same area shown in **g** with UV irradiation after hydrogel embedding and before denaturation. **i**, overlay. Scale bars are corrected for the expansion factor. (**a-i**), 5 µm

### SUPP FIG. 32

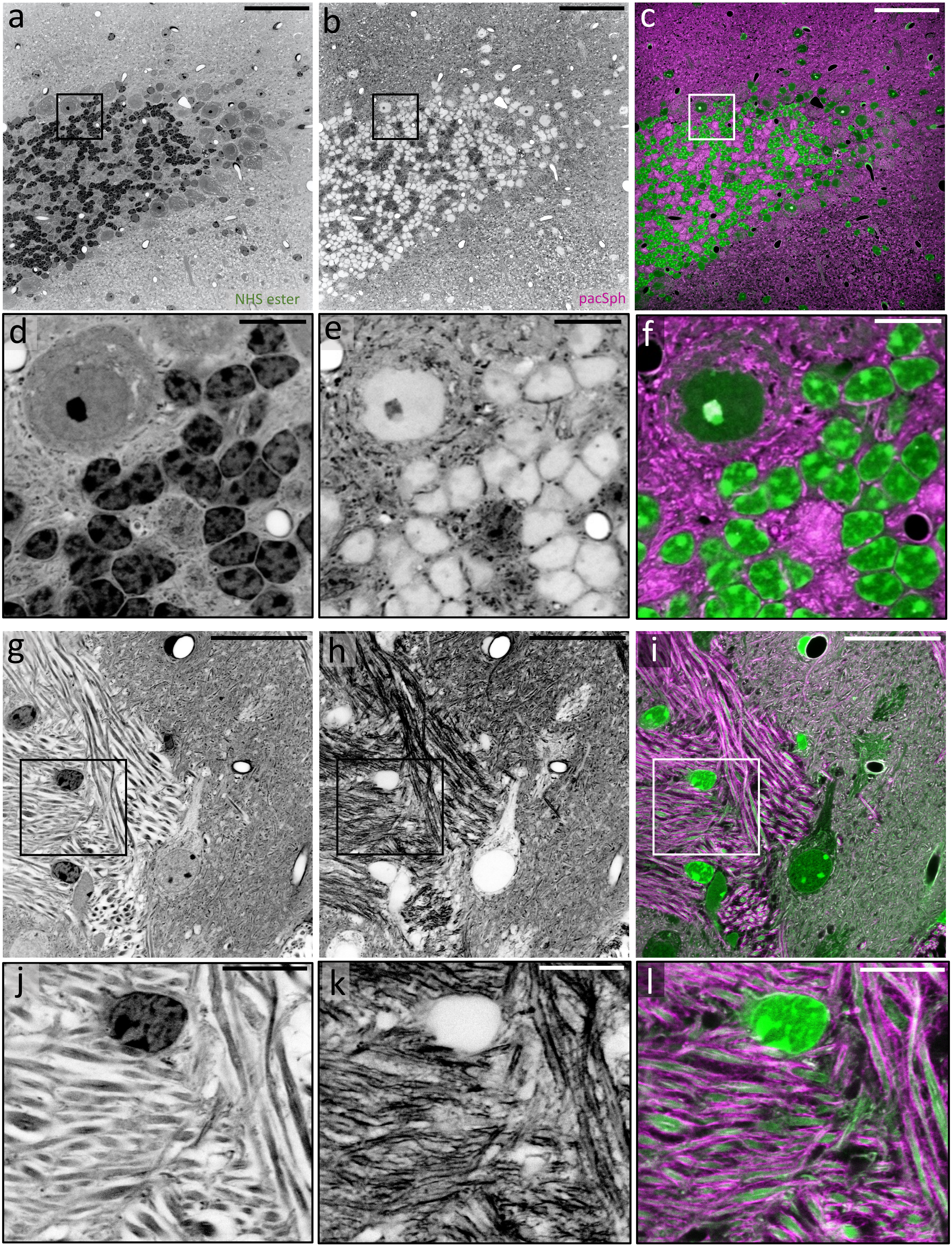

**Supplementary Figure 32: Examples of pacSph and NHS ester differential pan-staining in 4-fold expanded mouse brain tissue sections.** **a**, **g**, NHS ester pan-stained tissue section expanded ~4-fold. **b**, **h**, pacSph pan-staining of the same area shown in **a** and **g**, respectively. **c**, **i**, overlay. **d**, **e**, **f**, magnified images of the areas outlined by the boxes in **a**, **b**, and **c**, respectively, showing cell bodies. **j**, **k**, **l**, magnified images of the areas outlined by the boxes in **g**, **h**, and **I**, respectively, showing myelinated axons. Scale bars are not corrected for the expansion factor. (**a-c**) 200 µm, (**d-f**, **j-l**) 30 µm, (**g-i**) 100 µm.
